# PTP1B is an intracellular checkpoint that limits T cell and CAR T cell anti-tumor immunity

**DOI:** 10.1101/2021.11.11.468140

**Authors:** Florian Wiede, Kun-Hui Lu, Xin Du, Mara N. Zeissig, Rachel Xu, Pei Kee Goh, Chrysovalantou E. Xirouchaki, Samuel J. Hogarth, Spencer Greatorex, Kevin Sek, Roger J. Daly, Paul A. Beavis, Phillip K. Darcy, Nicholas K. Tonks, Tony Tiganis

## Abstract

Immunotherapies aimed at alleviating the inhibitory constraints on T cells have revolutionised cancer management. To date, these have focused on the blockade of cell surface checkpoints such as PD-1. Herein we identify protein-tyrosine-phosphatase 1B (PTP1B) as an intracellular checkpoint that is upregulated in T cells in tumors. We show that the increased PTP1B limits T cell expansion and cytotoxicity to contribute to tumor growth. T cell-specific PTP1B deletion increased STAT-5 signaling and this enhanced the antigen-induced expansion and cytotoxicity of CD8^+^ T cells to suppress tumor growth. The pharmacological inhibition of PTP1B recapitulated the T cell-mediated repression of tumor growth and enhanced the response to PD-1 blockade. Furthermore, the deletion or inhibition of PTP1B enhanced the efficacy of adoptively-transferred chimeric-antigen-receptor (CAR) T cells against solid tumors. Our findings identify PTP1B as an intracellular checkpoint whose inhibition can alleviate the inhibitory constraints on T cells and CAR T cells to combat cancer.

**STATEMENT OF SIGNIFICANCE:** Tumors subvert anti-tumor immunity by engaging checkpoints that promote T-cell exhaustion. Here we identify PTP1B as an intracellular checkpoint and therapeutic target. We show that PTP1B is upregulated in intra-tumoral T-cells and that its deletion or inhibition enhances T-cell anti-tumor activity and increases CAR T-cell effectiveness against solid tumors.

## INTRODUCTION

Immune checkpoints are crucial for modulating the magnitude of immune responses to minimise collateral tissue damage and overt autoimmunity during anti-microbial defence and T cell homeostasis ^1^. Tumors hijack immune checkpoints by upregulating on their cell surface ligands for T cell inhibitory receptors. In particular, the inflammatory tumor microenvironment is instrumental in driving the expression of ligands such as PD-L1, which engage T cell inhibitory receptors, such as PD-1. The engagement of PD-1 on activated T cells by PD-L1 on tumor cells, or other cells in the tumor microenvironment, suppresses anti-tumor responses and promotes T cell exhaustion, a state of deteriorated T cell function ^1,2^. Antibodies blocking the PD-1 immune checkpoint have revolutionised cancer therapy, resulting in durable clinical responses, especially in immunogenic tumors with abundant T cell infiltrates ^3^. However, not all tumors are responsive to PD-1 checkpoint blockade and both primary and adaptive resistance can occur ^3^. An alternative approach by which to alleviate the inhibitory constraints on T cells and to overcome T cell exhaustion in the tumor microenvironment may be to target protein-tyrosine-phosphatases (PTPs) that antagonise T cell function ^4^. Such PTPs might include the closely related SHP-1 and SHP-2 that are thought to act redundantly to mediate PD-1 inhibitory signaling in T cells ^5,6^, or TCPTP (also known as PTPN2), which antagonises T cell receptor (TCR) and cytokine signaling to maintain T cell tolerance ^7–10^ and whose deletion enhances both T cell-mediated immune surveillance and the anti-tumor activity of adoptively transferred T cells ^11,12^.

The prototypic tyrosine-specific phosphatase PTP1B (encoded by *PTPN1*) is expressed ubiquitously and has been implicated in various physiological and pathological processes ^13^. PTP1B shares a high degree of structural and sequence identity with TCPTP and the two phosphatases can function together to coordinate diverse biological processes ^13^, including, for example, the CNS control of energy and glucose homeostasis ^14,15^. However, it remains unknown whether PTP1B has a role in T cells. PTP1B is fundamentally important in metabolism and its global deletion results in obesity resistance and improved glucose homeostasis, attributable to the promotion of CNS leptin and peripheral insulin signaling respectively ^16–18^. As such, PTP1B has long been considered an exciting therapeutic target for combating metabolic syndrome ^13^. Indeed, its systemic inhibition with trodusquemine (MSI-1436), a specific allosteric inhibitor of PTP1B ^19^, promotes weight loss and improves glucose homeostasis in mice ^19,20^. Yet other studies have shown that the inhibition of PTP1B with MSI-1436 might also be useful for the treatment of neurological disorders, including Rett syndrome ^21^. Moreover, several studies point towards PTP1B serving as a potential therapeutic target in solid tumors ^19,22–24^. In particular, PTP1B levels are elevated in breast cancer where it is thought to contribute to tumor growth ^22,23^. Consistent with this, the global deletion of PTP1B attenuates the development of mammary tumors driven by mutant ErbB2 in mice ^22,23^, whereas MSI-1436 attenuates the growth of xenografts in SCID-beige mice and the metastasis of ErbB2-driven mammary tumors in transgenic mice ^19^. These studies have focused on the cell autonomous contributions of PTP1B in breast cancer tumorigenesis. However, the extent to which PTP1B may affect tumor growth by eliciting non-cell autonomous effects remains unclear. Although PTP1B heterozygosity has been reported to attenuate the growth of syngeneic Eμ-myc-driven lymphomas in mice by enhancing the immunogenicity of antigen-presenting dendritic cells (DCs), paradoxically its complete deletion impairs DC maturation and function ^25,26^. Therefore, it remains unclear whether the systemic inhibition of PTP1B might enhance or otherwise repress anti-tumor immunity.

Cytokine-induced JAK (Janus kinase)/STAT (Signal transducer and activator of transcription) signaling is fundamentally important for all aspects of immunity ^27^. In T cells the induction of STAT-5 signaling downstream of common γ (γc) chain cytokines, such as interleukin (IL)-2, is essential for T cell activation, homeostasis, survival and memory formation ^28^. It is well established that PTP1B can attenuate JAK/STAT signaling by dephosphorylating and inactivating JAK-2 and Tyk2 ^29,30^. In this study we show that PTP1B attenuates JAK/STAT-5 signaling and antagonises the expansion and activation of T cells. We show that PTP1B abundance is increased in intratumoral CD8^+^ T cells to repress anti-tumor immunity. Using genetic approaches, we establish that the deletion of PTP1B in T cells promotes STAT-5 signaling to facilitate the antigen-induced expansion, activation and cytotoxicity of CD8^+^ T cells to attenuate the growth of solid tumors. Importantly, we report that the inhibition of PTP1B in T cells enhances not only endogenous T cell-mediated anti-tumor immunity and the response to anti-PD-1 therapy, but also the efficacy of adoptively transferred T cells and CAR T cells to repress the growth of solid tumors. Our findings define a novel intracellular checkpoint and actionable therapeutic target for enhancing the anti-tumor activity of T cells.

## RESULTS

### PTP1B deletion in the hematopoietic compartment represses solid tumor growth

To explore whether PTP1B may elicit non-cell autonomous effects on the growth of tumors we generated syngeneic mammary tumors by implanting ovalbumin (OVA)-expressing AT3 (AT3-OVA) murine mammary tumor cells into the inguinal mammary fat pads of *Ptpn1^+/+^*, *Ptpn1^+/–^* or *Ptpn1^−/−^* C57BL/6 mice ^17^ and assessed tumor growth (**Fig. 1a-b**). We found that mammary tumor growth was markedly attenuated in *Ptpn1^−/−^* mice so that the resultant tumors were less than half the size of those in *Ptpn1^+/+^* control mice; intermediate effects were seen in *Ptpn1^+/–^* C57BL/6 mice indicating that even partial PTP1B deficiency is sufficient to attenuate tumor growth. The repression of tumor growth was accompanied by an increased abundance of tumor-infiltrating lymphocytes (TILs) (**Fig. 1c**), which in many tumors, including breast cancer, are associated with improved survival and response to therapy ^31^. These TILs were predominantly CD44^hi^CD62L^lo^ (CD44 marks activated T cells and CD62L lymphoid-resident T cells ^32,33^) effector/memory CD8^+^ T cells, which can repress tumor growth (**Fig. 1c**; **Fig. S1a**), but also included CD4^+^ T cells, natural killer (NK) cells and DCs (**Fig. 1c**), which promote anti-tumor immunity ^3,34^. In addition, the global deletion of PTP1B was accompanied by the recruitment of CD4^+^ regulatory T cells (T_regs_), tumor-associated macrophages (TAMs) and both granulocytic and monocytic myeloid-derived suppressor cells (MDSCs) (**Fig. 1c**; **Fig. S1b**) that are immunosuppressive ^3^. To determine the extent to which the repression of syngeneic tumor growth may be ascribed to stromal differences, changes in the hormonal milieu, or otherwise differences in the hematopoietic compartment, we repeated these experiments using chimeric mice; lethally irradiated Ly5.1^+^ C57BL/6 recipient mice were reconstituted with congenically-marked Ly5.2^+^ *Ptpn1^+/+^* control or *Ptpn1^−/−^* bone marrow, before the implantation of AT3-OVA tumor cells into the inguinal mammary fat pads (**Fig. 1d**). We found that PTP1B deficiency in the hematopoietic compartment alone completely phenocopied the effects of global PTP1B deletion on the repression of tumor growth (**Fig. 1e**) and the abundance of TILs (**Fig. 1f**). Therefore, the deletion of PTP1B in the immune compartment is sufficient to increase TILs and suppress tumor growth.

**Figure 1.**
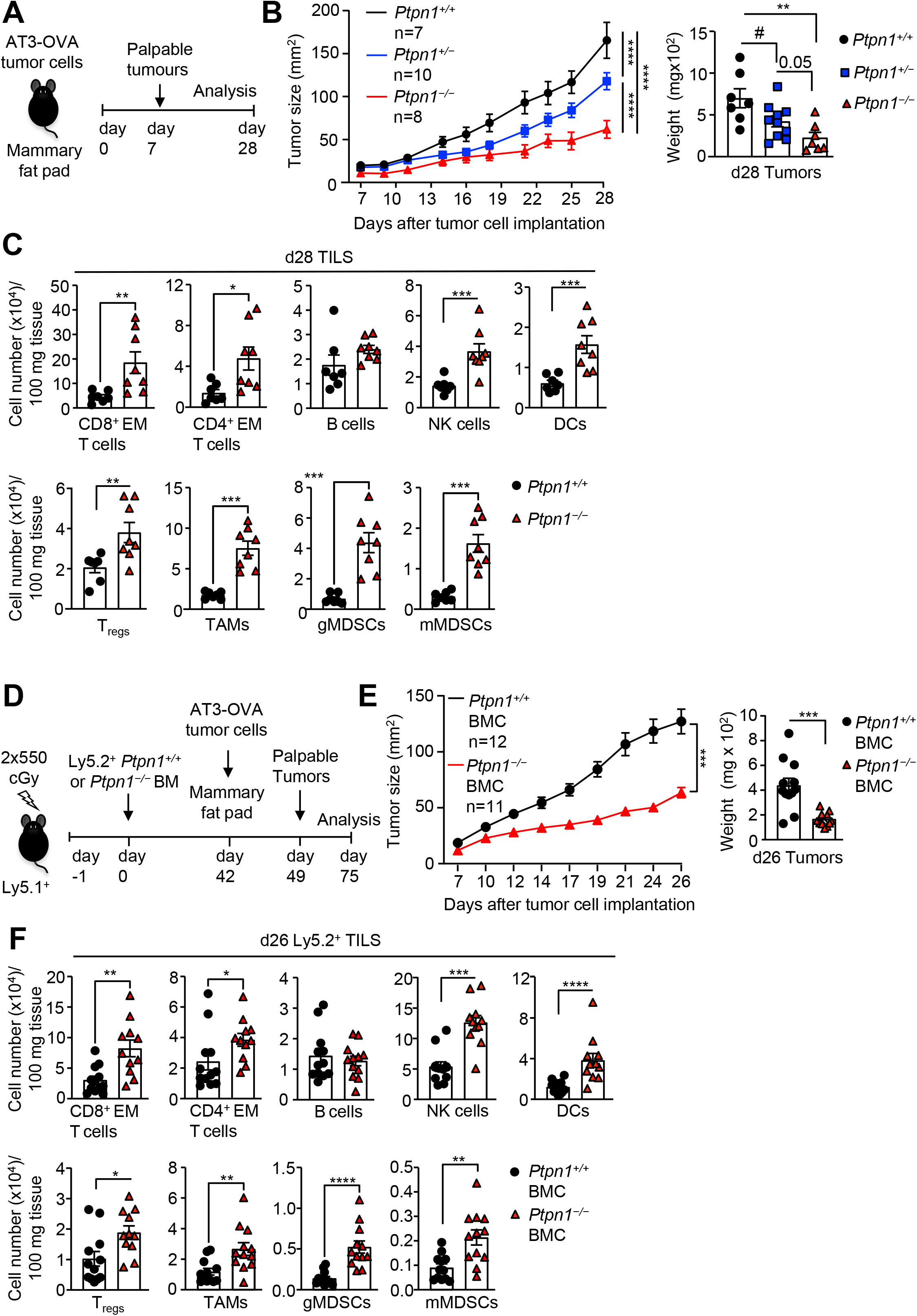
PTP1B deletion in immune cells enhances anti-tumor immunity. **a-c**) AT3-OVA tumor cells were implanted into the fourth inguinal mammary fat pads of *Ptpn1^+/+^*, *Ptpn1^+/−^* or *Ptpn1^−/−^* littermate mice and tumor growth and weights measured. **c**) Tumor-infiltrating lymphocytes (TILs) including CD44^hi^CD62L^lo^ CD8^+^ and CD4^+^ effector/memory (EM) T cells, B220^+^ B cells, NK1.1^+^TCRβ^−^ (NK) cells, CD11c^+^ dendritic cells (DCs), CD4^+^CD25^+^FoxP3^+^ regulatory T cells (T_regs_), CD11b^+^F4/80^hi^Ly6C^−^Ly6G^−^ tumour associated macrophages (TAMs), granulocytic CD11b^+^F4/80^hi/lo^Ly6C^int^Ly6G^+^ (gMDSCs) and monocytic CD11b^+^F4/80^hi/lo^ Ly6C^+^Ly6G^−^ (mMDSCs) myeloid-derived suppressor cells were analysed by flow cytometry. **d-e**) Lethally irradiated Ly5.1 mice were reconstituted with bone marrow cells derived from *Ptpn1^+/+^* or *Ptpn1^−/−^* mice. After 4 weeks of reconstitution, AT3-OVA tumor cells were implanted into the fourth inguinal mammary fat pads of PTP1B-deficient or control bone marrow chimeras (BMC) and tumor growth monitored and final weights measured and **f**) tumor-infiltrating analysed by flow cytometry. Representative results (means ± SEM) from at least two independent experiments are shown. For tumor sizes in (b, e) significance was determined using a 2-way ANOVA Test, for tumor weights in (b, e) and for immune cell infiltrates in (c, f) using 2-tailed Mann-Whitney U Test or 1-way ANOVA Test and for tumour weights in (b) using a 2-tailed Mann-Whitney U Test (^#^p<0.05).

### PTP1B is increased in tumor T cells

Although PTP1B deficiency may potentially affect both the innate and adaptive immune compartments to promote anti-tumor immunity, we focused our attention on CD8^+^ T cells since they were the most abundantly represented TILs in syngeneic tumors in PTP1B-deficient mice (**Fig. 1c**). As PTP1B’s function in T cells remains unknown, we first assessed its expression in lymphoid versus intratumoral T cells. To this end we adoptively transferred congenically-marked (Ly5.2^+^) naive (CD44^lo^CD62L^hi^) CD8^+^ T cells expressing the OT-I TCR, which is specific for OVA, into immunocompetent Ly5.1^+^ mice bearing AT3-OVA mammary tumors and assessed PTP1B protein in both the endogenous and adoptively transferred CD8^+^ T cell subsets in the spleens versus tumors by flow cytometry (**Fig. 2a; Fig. S2; Gating Strategy Fig. S3**). Endogenous splenic T cells from tumor-bearing mice were largely naïve in phenotype, but also included a smaller proportion of CD44^hi^CD62L^hi^ central memory and CD44^hi^CD62L^lo^ effector/memory T cells (**Fig. S3**). By contrast, adoptively transferred OT-I T cells had central memory or effector/memory phenotypes in spleens and an effector/memory phenotype in tumors (**Fig. S3**); the central memory and effector/memory OT-I T cells in the spleens of tumour-bearing mice likely originated from adoptively transferred naïve T cells that had engaged antigen and had recirculated from the tumor to lymphoid organs ^32,33^. In endogenous splenic CD8^+^ T cells, PTP1B protein as assessed by flow cytometry was moderately elevated in central memory and effector/memory T cells when compared to naïve T cells (**Fig. 2a**). By contrast, PTP1B protein was elevated by approximately 2-fold in both endogenous effector/memory CD8+ T cells and adoptively transferred OT-I effector/memory T cells in tumors, when compared to the corresponding effector/memory T cells in spleens (**Fig. 2a**); the increased PTP1B protein in intratumoral versus splenic effector/memory T cells was accompanied by increased *Ptpn1* mRNA expression, as assessed by quantitative real time PCR (**Fig. 2b**). Moreover, PTP1B protein, as assessed by immunoblotting, was elevated by more than 2-fold in tumor effector/memory OT-I T cells when compared to splenic effector/memory OT-I T cells (**Fig. 2c**). Therefore, PTP1B levels are elevated in intratumoral T cells and the induction of PTP1B is not an outcome of T cell differentiation or activation, but rather an outcome of conditions within the tumor microenvironment. In line with our findings in mice, an analysis of publicly available single cell RNA-sequencing data of immune cells isolated from the tumors of melanoma patients, revealed that *PTPN1* mRNA was increased in human intratumoral CD8^+^ T cells with an effector [high for interferon (IFN) γ (*IFNG*), granzyme B (*GZMB*) and perforin (*PRF1*)] or exhausted [high for TIM3 (*HAVCR2*), LAG3 (*LAG3*) and PD-1 (*PDCD1*)] phenotype (**Fig. 2d**), otherwise associated with a poor response to PD-1 checkpoint blockade ^35^, as compared to PTP1B levels in intratumoral T cells with a stem cell or memory phenotype [high for *TCF7*] (**Fig. 2d**), which is associated with a positive clinical outcome to PD-1 blockade ^35^. This suggests that, as in murine models, PTP1B levels are elevated in human intratumoral T cells and may negatively regulate T cell responses and the response to immunotherapy.

**Figure 2.**
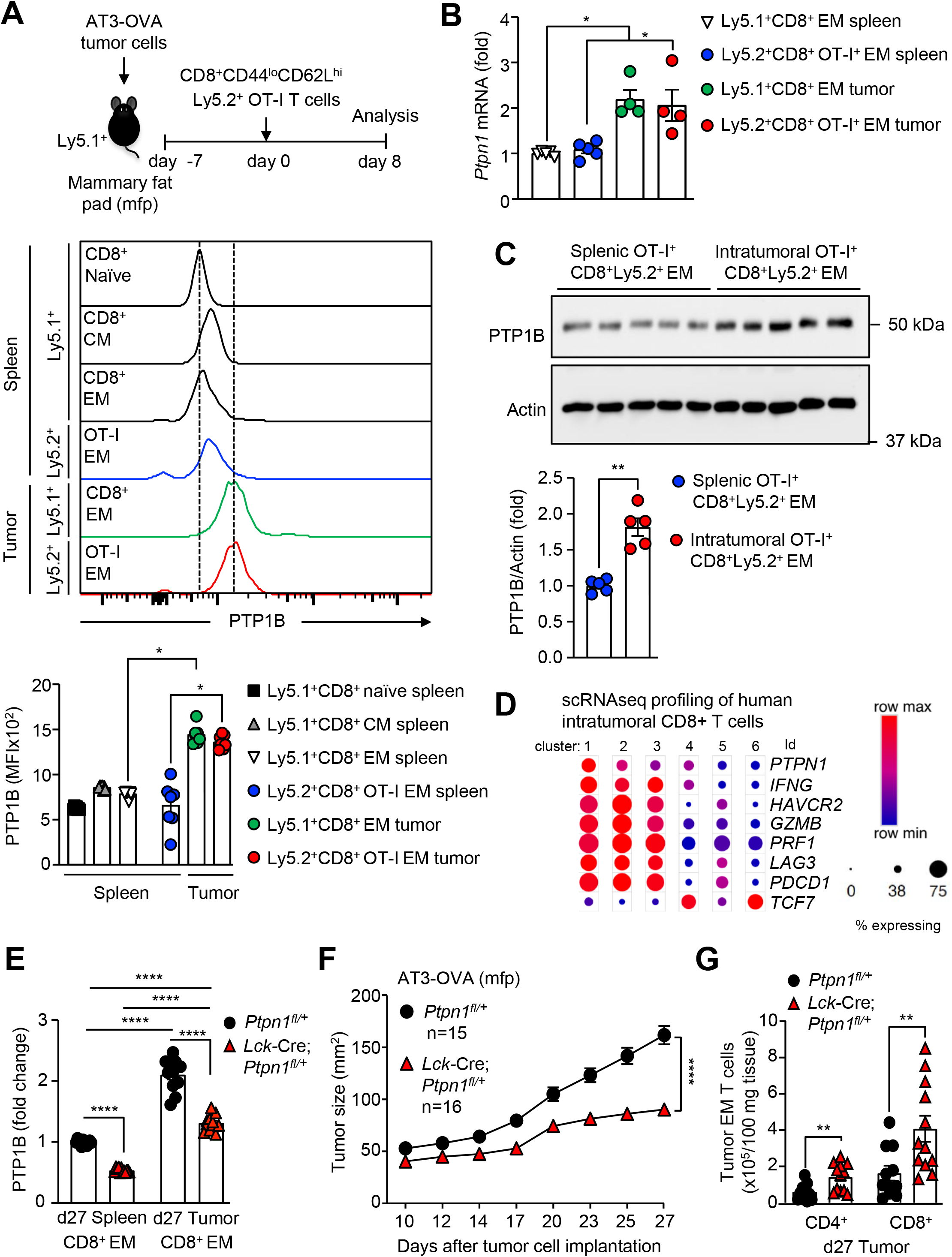
Increased PTP1B in intratumoral CD8^+^ T cells limits anti-tumor immunity. **a-c**) Naïve (CD44^lo^CD62L^hi^) Ly5.2^+^CD8^+^OT-I^+^ cells from OT-I;*Ptpn1^fl/fl^* or T cell-specific PTP1B-deficient OT-I;*Lck*-Cre;*Ptpn1^fl/fl^* mice were adoptively transferred into Ly5.1 mice bearing established (40-50 mm^2^) AT3-OVA mammary tumors. **a**) 8 days post adoptive transfer PTP1B protein levels were determined by flow cytometry in splenic or intratumoral Ly5.1^+^CD8^+^ recipient or Ly5.2^+^CD8^+^OT-I^+^ donor T cells with a naïve, central/memory (CM; CD44^hi^CD62L^hi^) or effector/memory (EM; CD44^hi^CD62L^lo^) phenotype. **b**) *Ptpn1* mRNA in FACS-purified splenic and intratumoral Ly5.1^+^CD8^+^ recipient and Ly5.2^+^OT-I^+^CD8^+^ EM T cells isolated on day 5 after adoptive transfer was assessed by quantitative real time PCR. **c**) PTP1B protein levels in FACS-purified splenic and intratumoral Ly5.2^+^CD8^+^OT-I^+^ donor EM T cells isolated on day 5 after adoptive transfer were assessed by immumoblotting. **d**) The broad institute single cell portal (https://singlecell.broadinstitute.org/single_cell) was used to interrogate publicly available single-cell RNA-sequencing (scRNA-seq) data performed on immune cells isolated from 48 tumor biopsies taken from 32 metastatic melanoma patients treated with checkpoint blockade therapy. Fine clustering of CD8+ T cell subsets revealed 6 clusters. **e-g**) AT3-OVA tumor cells were implanted orthotopically into the fourth inguinal mammary fat pads of control (*Ptpn1^fl/+^*) mice or T cell-specific PTP1B-heterozygous (*Lck*-Cre;*Ptpn1^fl/+^*) mice and **e**) PTP1B protein levels in CD8^+^ EM T cells were analysed by flow cytometry. **f**) Tumor growth was monitored and **g**) tumor-infiltrating CD4^+^ and CD8^+^ EM T cells were analysed by flow cytometry. Representative results (means ± SEM) from at least two independent experiments are shown. Significance in (a, e) was determined using a one-way ANOVA Test, in (b-c) a 2-tailed Mann-Whitney U Test, in (f) a two-way ANOVA Test and in g) a one-way ANOVA Test.

### PTP1B deletion enhances T cell anti-tumor immunity

To explore whether the induction of PTP1B in intratumoral T cells might limit T cell function and anti-tumor immunity, and if enhanced T cell responses in *Ptpn1^−/−^* mice might contribute to the repression of tumor growth, we sought to delete PTP1B in T cells using the *Lck*-Cre transgene; the *Lck*-Cre transgene specifically deletes floxed alleles at the early stages T cell development in the thymus ^7^. As expected, PTP1B was reduced in total thymocytes and efficiently deleted in T cells, but not in myeloid cells, B cells or non-lymphoid organs (**Fig. S4, S5**). The T cell-specific deletion of PTP1B increased total thymocyte numbers at each stage of development without affecting positive selection (**Fig. S6a-e**) and resulted in increased peripheral naïve, effector/memory and central memory T cells in lymphoid organs (**Fig. S6f-i**); the increase in peripheral T cell numbers was also noted in *Ptpn1^−/−^* mice with a global deletion in PTP1B (**Fig. S7a-b**). The increased T cell abundance was not associated with overt pathology, including lymphomas or CD4^+^CD8^+^CD3^−^ leukemias, when mice were aged for 1 year (**Fig. S8a**) or 2 years (**Fig. S9a**). Moreover, there were no signs of systemic inflammation or autoimmunity, as assessed by monitoring for lymphocytic infiltrates in non-lymphoid organs (**Fig. S8b-c**; **Fig. S9b-c**), the presence of inflammatory cytokines and anti-nuclear antibodies in serum (**Fig. S10a-b**; **Fig. S10e-f**), and there were no signs of tissue damage, as assessed histologically (**Fig. S10c**; **Fig. S10g**) and by monitoring for the presence of the liver enzymes ALT and AST in serum (**Fig. S10d**).

To assess the influence of PTP1B deletion on T cell-mediated anti-tumor immunity, we implanted AT3-OVA mammary tumor cells ^36^ into the inguinal mammary fat pads of control (*Ptpn1^fl/fl^*) and T cell-specific PTP1B-deficient mice (*Lck*-Cre;*Ptpn1^fl/fl^*) female mice (**Fig. 3a**). We found that the growth of orthotopic AT3-OVA mammary tumors was significantly repressed (**Fig. 3a**) and this was associated with a significant improvement in survival (**Fig. S11a)**; notably the repression of AT3-OVA mammary tumor growth paralleled that in *Ptpn1^−/−^* mice consistent with the repression of tumor growth in *Ptpn1^−/−^* mice being T cell dependent (**Fig. 1a**). Indeed, the repression of tumor growth in *Lck*-Cre;*Ptpn1^fl/fl^* mice was accompanied by the accumulation of effector/memory CD4^+^ and CD8^+^ T cells (**Fig. S11b**), consistent with T cell-mediated tumour eradication; no overt differences in tumour cell proliferation or apoptosis or tumour angiogenesis were evident (**Fig. S12**). The deletion of PTP1B also significantly repressed the growth of other syngeneic tumors, including OVA-expressing B16F10 (B16F10-OVA) melanoma cells (**Fig. S11c**) and MC38 (MC38-OVA) colorectal cancer cells (**Fig. S11d-e**) implanted into the flanks of male mice. Moreover, the deletion of PTP1B in *Lck*-Cre;*Ptpn1^fl/fl^* mice also repressed the growth of non-OVA expressing AT3, B16F10 and MC38 syngeneic tumors (**Fig. S13a-e**) and the repression of tumor growth was similarly accompanied by the accumulation of effector/memory T cells (**Fig. S13b**); in contrast, the *Lck*-Cre transgene alone had no effect on syngeneic tumor growth (**Fig. S14**). Therefore, PTP1B deficiency can enhance T cell-mediated anti-tumor immunity, irrespective of whether tumors are overtly immunogenic and express OVA or not.

**Figure 3.**
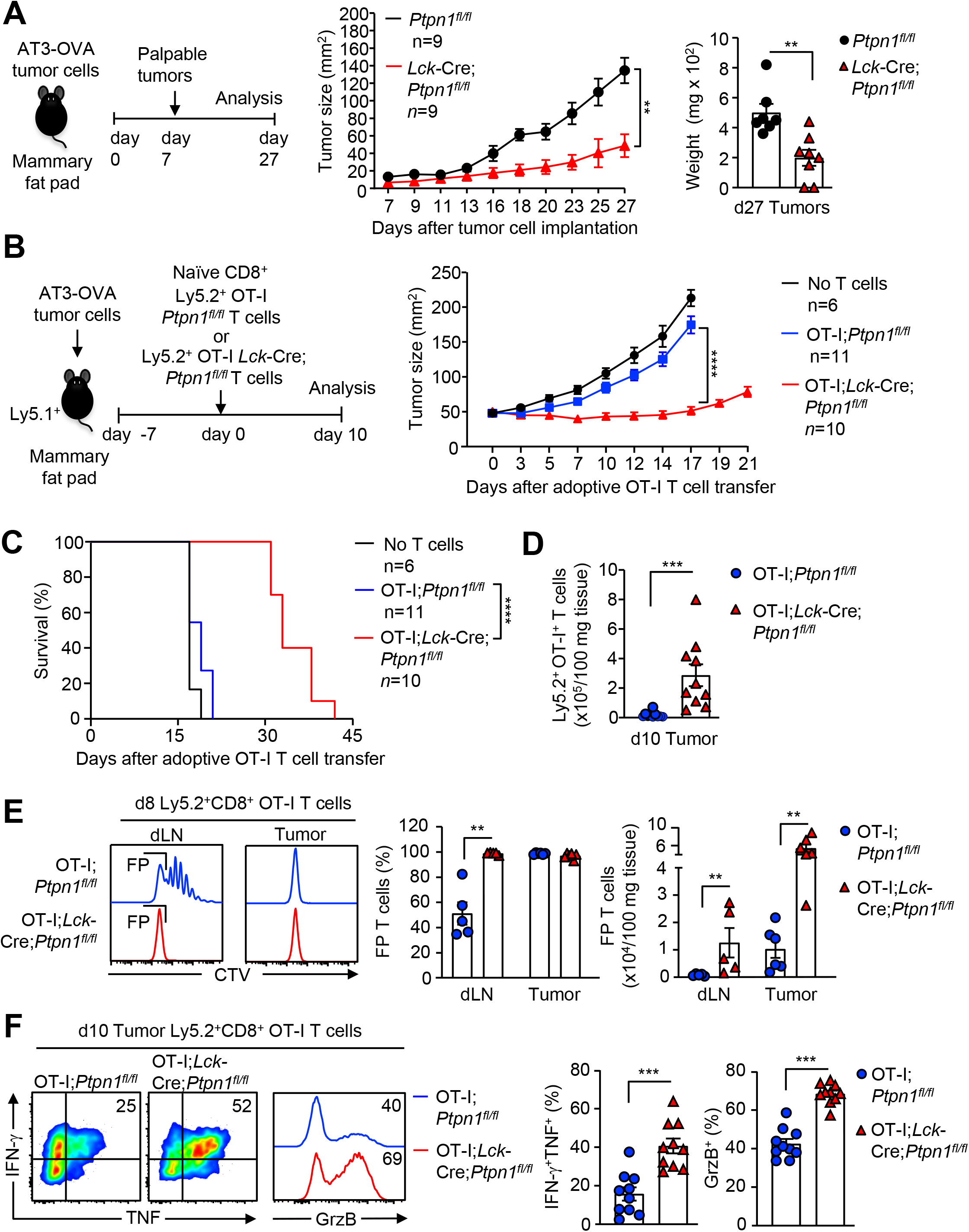
PTP1B deletion in T cells enhances anti-tumor immunity. **a**) AT3-OVA tumor cells were implanted orthotopically into the fourth inguinal mammary fat pads of control (*Ptpn1^fl/fl^*) mice or T cell-specific PTP1B-deficient (*Lck*-Cre;*Ptpn1^fl/fl^*) mice. Tumor growth was monitored and final tumor weights determined. **b-d**) Naïve (CD44^lo^CD62L^hi^) Ly5.2^+^CD8^+^OT-I^+^ T cells from OT-I;*Ptpn1^fl/fl^* and OT-I;*Lck*-Cre;*Ptpn1^fl/fl^* mice were adoptively transferred into Ly5.1 mice bearing established (40-50 mm^2^) AT3-OVA mammary tumors and **b**) tumor growth, **c**) survival and **d**) Ly5.2^+^CD8^+^ OT-I T cell infiltrates (day 10 post adoptive transfer) determined. **e**) CTV-labelled naïve Ly5.2^+^CD8^+^OT-I^+^ T cells from OT-I;*Ptpn1^fl/fl^* or OT-I;*Lck*-Cre;*Ptpn1^fl/fl^* mice were adoptively transferred into Ly5.1 mice bearing established (40-50 mm^2^) AT3-OVA mammary tumors and fast proliferating (FP) T cells monitored in the draining lymph nodes (dLN) and tumors. **f**) Tumor-infiltrating T cells from (d) were stimulated with PMA/Ionomycin in the presence of Golgi Stop/Plug and stained for intracellular IFN-γ and TNF. Intracellular granzyme B (GrzB) was detected in unstimulated tumor-infiltrating CD8^+^ T cells. For tumor growth in (a, b) significance was determined using 2-way ANOVA Test. For tumor weights in (a), T cell infiltrates in (c, e) and for intracellular IFN-γ, TNF and Grzb levels (f) significance was determined using 2-tailed Mann-Whitney U Test. In (c) significance was determined using Log-rank (Mantel-Cox) test.

As PTP1B was induced by approximately 2-fold in intratumoral T cells (**Fig. 2a, c**), we next assessed whether the heterozygous deletion of PTP1B in T cells might be sufficient to alleviate the inhibitory effects on T cell function and enhance anti-tumor immunity. We found that PTP1B heterozygosity largely corrected the increased PTP1B in intratumoral T cells (**Fig. 2e**) and repressed the growth of AT3-OVA mammary tumors in *Lck*-Cre;*Ptpn1^fl/+^* mice (**Fig. 2f**) and this was accompanied by increased CD4^+^ and CD8^+^ effector/memory T cells in tumors (**Fig. 2g**). Taken together these results point towards PTP1B serving as an intracellular checkpoint and its induction repressing T cell-mediated anti-tumor immunity.

To specifically explore the contributions of CD8^+^ T cells, we next asked if the deletion of PTP1B could promote the tumor-specific activity of transgenic OT-I CD8^+^ T cells adoptively transferred into mice bearing OVA-expressing tumors. Naive control (*Ptpn1^fl/fl^*) or PTP1B-deficient (*Lck*-Cre;*Ptpn1^fl/fl^*) congenically-marked (Ly5.2^+^) OT-I CD8^+^ T cells were adoptively transferred into immunocompetent and non-irradiated (Ly5.1^+^) C57BL/6 hosts with AT-3-OVA mammary tumors (**Fig. 3b-f)**. Adoptively transferred control CD8^+^ T cells had little effect on tumor growth (**Fig. 3b**). By contrast, PTP1B-deficient CD8^+^ T cells significantly repressed the growth of AT3-OVA mammary tumors (**Fig. 3b**) and significantly increased the survival of mice (**Fig. 3c**). The repression of tumor growth was accompanied by an increased abundance of PTP1B-deficient tumor-infiltrating effector/memory T cells (**Fig. 3d; Fig. S15**); no overt differences in tumor cell proliferation or apoptosis or tumour angiogenesis were evident (**Fig. S16**). The increased abundance of T cells was associated with a marked increase in T cell proliferation in the draining lymph nodes, as reflected by the presence of ‘fast proliferating’ T cells [assessed by the dilution of the fluorescent dye CellTrace™ Violet (CTV), which defines distinct populations of dividing cells; **Fig. 3e**], increased T cell activation, as assessed by the expression of CD44 and increased cytotoxicity, as assessed by the expression of IFNγ and TNF as well as granzyme B (**Fig. 3f; Fig. S15**), which accumulates in cytolytic granules and mediates tumour cell killing when released into the immunological synapse. These results indicate that PTP1B-deficiency in CD8^+^ T cells enhances the expansion, activation and cytotoxic activity of CD8^+^ T cells to repress tumor growth. Moreover, these results are consistent with elevated PTP1B in intratumoral CD8^+^ T cells repressing T cell-mediated anti-tumor immunity to facilitate tumor growth.

### PTP1B deletion promotes STAT-5-dependent T cell expansion, activation and anti-tumor immunity

To elucidate how PTP1B-deficiency promotes T cell-mediated anti-tumor immunity we first assessed whether PTP1B deletion might enhance αβ T cell receptor (TCR) signaling that is essential for T cell activation and function ^37^. We found that PTP1B-deficiency did not result in gross changes in protein tyrosine phosphorylation or the activation of canonical TCR signaling intermediates after crosslinking the αβ TCR on naïve CD8^+^ T cells with anti-CD3ε (**Fig. S17a-d**). However, PTP1B-deficiency significantly enhanced the TCR-instigated activation of T cells and the generation of effector T cells, as monitored by i) the increased cell surface expression of CD44, CD69 and CD25 [IL-2 receptor (IL2R) α chain] (**Fig. 4a, Fig. S18a, S20a**), markers of T cell activation and responsiveness to IL-2 and ii) the decreased expression of CD62L (**Fig. 4a, Fig. S18a, S20a**), which allows for T cells to exit lymphoid organs and act as effectors. PTP1B deletion was also associated with a marked enhancement in naïve CD8^+^ T cell proliferation and/or resulted in increased T cell numbers across a range of α-CD3ε concentrations, as reflected by the dilution of CTV and the number of resultant cells (**Fig. 4b, Fig. S18b, S20b**). The enhanced proliferative capacity was reaffirmed by monitoring the incorporation of BrdU (measures S phase entry) in α-CD3ε stimulated T cells (**Fig. 18c**). Importantly, even at saturating α-CD3ε concentrations, when differences in CTV dilution where not evident, PTP1B-deficiency resulted in increased cell numbers, consistent with increased survival (**Fig. 4b, S18b, S20b**). In line with this, the deletion of PTP1B was also accompanied by decreased apoptosis in proliferating cells, as assessed by staining for Annexin V (**Fig. S18d**). PTP1B deficiency also enhanced the expansion of naïve OT-I T cells challenged with cognate OVA peptide SII**N**FEKL (N4). Importantly, PTP1B deficiency similarly enhanced responses to the altered peptide ligands SI**Y**NFEKL (Y3) or SII**Q**FEKL (Q4) that have progressively lower affinities (N4>Y3>>Q4) for the OT-I TCR (**Fig. S19**), consistent with PTP1B deficiency promoting T cell expansion without altering the threshold for TCR-instigated responses. The activation and expansion of naive T cells is reliant not only on signaling from the TCR (signal 1) and co-receptors such as CD28 (signal 2), but also cytokines such as IL-2 (signal 3) ^28,37^. IL-2 signals via JAK protein-tyrosine-kinases (PTKs) and STAT-5 to drive the expression of genes that are essential for T cell proliferation, survival, differentiation and cytotoxicity ^28,38,39^. PTP1B deficiency increased basal STAT-5 Y694 phosphorylation (p-STAT-5) as monitored by flow cytometry in naïve and central memory T cells and this was accompanied by an increased abundance of BCL-2, which promotes cell survival (**Fig. 4c**), and Tbet and Eomes (**Fig. 4d**), which are transcriptional targets of STAT-5 that promote CD8^+^ T cell cytotoxicity and CD4^+^ T cell differentiation ^28^. Moreover, PTP1B deficiency enhanced STAT-5 signaling in response to IL-2 in effector T cells (**Fig. 4e**), as well as p-STAT-5 downstream of IL-7 and IL-15 in different T cell subsets (**Fig. S21**). The enhanced p-STAT-5 was accompanied by increased Tyk-2 Y1054/Y1055 (p-Tyk-2) phosphorylation in response to IL-2 and IL-15 (**Fig. S20c**). The activation of JAK PTKs by cytokines also results in phosphatidylinositol 3-kinase (PI3K)/AKT and Ras/mitogen-activated protein kinase (MAPK) signaling. As expected, PTP1B deletion also enhanced IL-2-induced PI3K/AKT signaling (AKT Ser-473 phosphorylation; p-AKT) and Ras/MAPK signaling (phosphorylation of ERK-1/2; p-ERK1/2) (**Fig. S20d**). The promotion of STAT-5 signaling occurred not only as a consequence of increasing JAK activation, but also as a result of increasing STAT-5 phosphorylation, since the inhibition of JAK PTKs with ruxolitinib in T cells pulsed with IL-2 was accompanied by prolonged albeit attenuated p-STAT-5 (**Fig. S20e**). Therefore, the deletion of PTP1B enhances the activation and expansion of T cells and promotes both JAK and STAT-5 signaling, but not TCR signaling.

**Figure 4.**
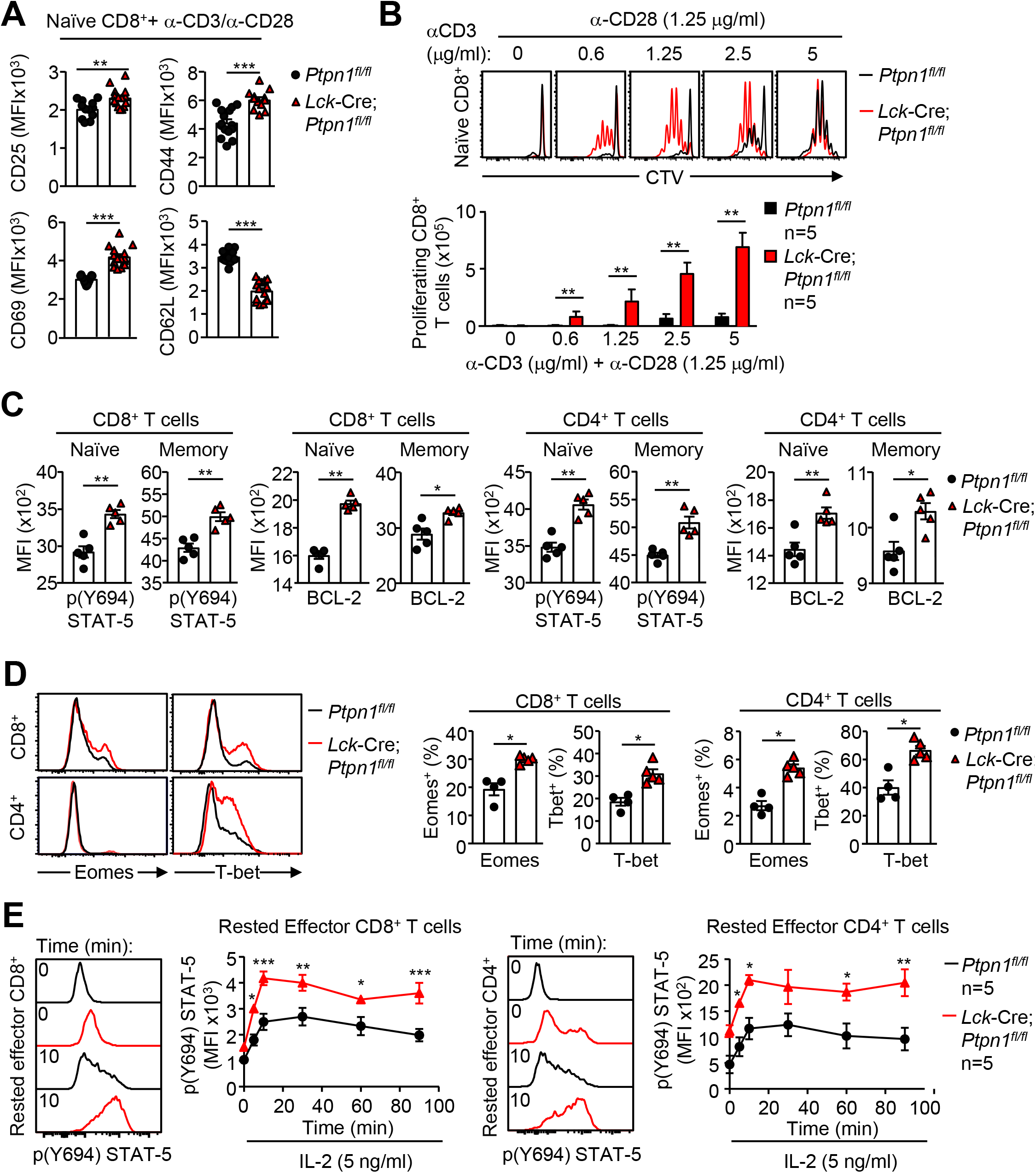
PTP1B deletion enhances T cell activation, proliferation, survival and STAT-5 signaling. **a**) Naïve (CD44^lo^CD62L^hi^) CD8^+^ T cells from *Ptpn1^fl/fl^* and *Lck*-Cre;*Ptpn1^fl/fl^* were stimulated with plate-bound α-CD3 (2.5 μg/ml) and α-CD28 (1.25 μg/ml) for 48 h and CD25, CD44, CD69 and CD62L levels determined by flow cytometry. **b**) CTV-labelled naïve CD8^+^ T cells from *Ptpn1^fl/fl^* and *Lck*-Cre;*Ptpn1^fl/fl^* mice were stimulated with the indicated concentrations of plate-bound α-CD3 plus α-CD28 (1.25 μg/ml) for 72 h and T cell proliferation (CTV dilution) determined by flow cytometry. **c**) Basal intracellular p-STAT-5 and BCL-2 levels were assessed by flow cytometry in naïve (CD44^lo^) and memory (CD44^hi^) CD4^+^ and CD8^+^ T cells from *Ptpn1^fl/fl^* and *Lck*-Cre;*Ptpn1^fl/fl^* mice. **d**) Intracellular Eomes and Tbet protein levels were assessed by flow cytometry in CD8^+^ and CD4^+^ T cells from *Ptpn1^fl/fl^* and *Lck*-Cre;*Ptpn1^fl/fl^* mice. **e**) *In vitro* generated rested effector CD8^+^ or CD4^+^ T cells were stimulated with IL-2 (5 ng/ml) for the indicated times and intracellular STAT-5 Y694 phosphorylation (p-STAT-5) assessed by flow cytometry. Representative results (means ± SEM) from at least two independent experiments are shown. Significance in (a-d) was determined using a 2-tailed Mann-Whitney U Test and in (e) using a 2-way ANOVA Test.

To assess whether PTP1B might have similar roles in human T cells we took advantage of CRISPR-Cas9 genome-editing to delete PTP1B in T cells derived from human peripheral blood mononuclear cells (**Fig. S22a**). As in murine T cells, we found that the deletion of PTP1B resulted in enhanced basal p-STAT-5 and the expression of BCL-2 family members, including BCL-2 and BCL-xL (**Fig. S22b**). PTP1B deletion also enhanced the expansion (as assessed by CTV dilution) and activation (as assessed by CD69 cell surface levels) of T cells after TCR crosslinking with α-CD3 (**Fig. S22c-d)**. Therefore, the deletion of PTP1B promotes STAT-5 signaling and facilitates the expansion and activation of both murine and human T cells.

Although JAK-2 and Tyk2 can signal via multiple STATs, the promotion of STAT-5 signaling in T cells is essential for proliferation after TCR ligation and for effector T cell responses ^28,38,39^. To explore the extent to which heightened STAT-5 signaling may contribute to the enhanced T cell expansion and activation in PTP1B-deficient T cells, we crossed *Lck*-Cre;*Ptpn1^fl/fl^* mice onto a *Stat5^fl/+^* heterozygous background in order to decrease p-STAT-5. *Stat5* heterozygosity largely corrected the elevated basal p-STAT-5, and the associated increase in BCL-2 protein, in naïve and central memory PTP1B-deficient CD4^+^ and CD8^+^ T cells (**Fig. 5a-b; S23a-b**). Moreover, *Stat5* heterozygosity corrected the increased abundance of peripheral CD4^+^ and CD8^+^ naïve and central memory T cells *in vivo* (**Fig. S23c**). Importantly, *Stat5* heterozygosity also largely, if not completely, corrected the enhanced α-CD3ε/α-CD28-induced T cell activation, as assessed by the correction in CD25, CD44 and CD69 cell surface expression, corrected the enhanced proliferation, as assessed by CTV dilution, and T cell numbers (**Fig. 5c-d**). Therefore, the deletion of PTP1B promotes STAT-5 signaling to facilitate the expansion and activation of T cells.

**Figure 5.**
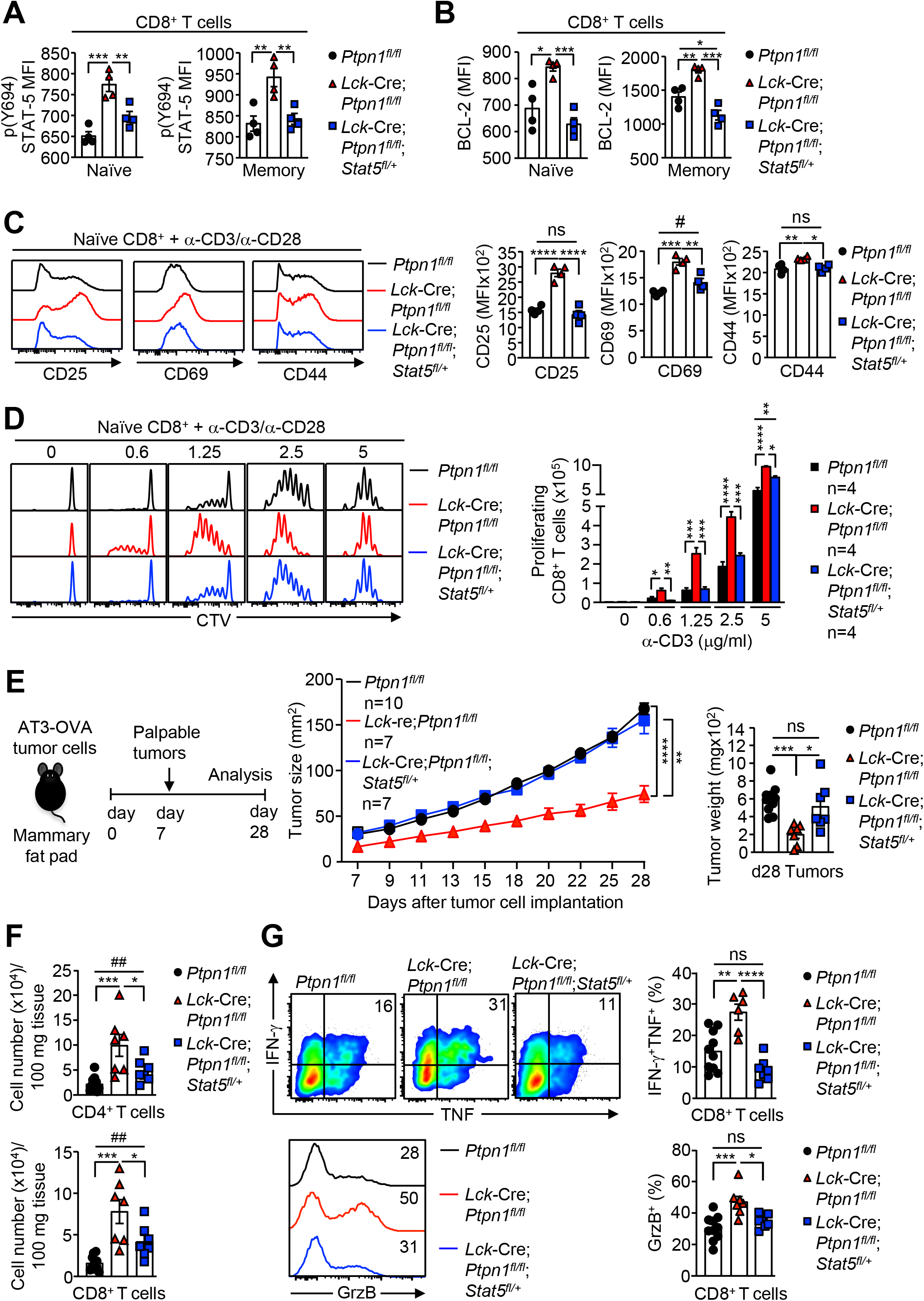
PTP1B-deficiency promotes STAT-5-dependent T cell activation and anti-tumor immunity. **a)** Basal intracellular p-STAT-5 and **b**) BCL-2 levels were assessed in naïve (CD44^lo^) and memory (CD44^hi^) lymph node CD8^+^ T cells from *Ptpn1^fl/fl^*, *Lck*-Cre;*Ptpn1^fl/fl^* and *Lck*-Cre;*Ptpn1^fl/fl^Stat5^fl/+^* mice. **c**) Naïve CD8^+^ (CD44^lo^CD62L^hi^) T cells were stimulated with plate-bound α-CD3 (2.5 μg/ml) plus α-CD28 (1.25 μg/ml) for 48 h and cell surface CD25, CD44 and CD69 levels (MFIs) determined. **d**) CTV-labelled naïve CD8^+^T cells were stimulated with the indicated concentrations of plate-bound α-CD3 plus α-CD28 (1.25 μg/ml) for 72 h and T cell proliferation (CTV dilution) assessed by flow cytometry. **e**) AT3-OVA tumor cells were implanted orthotopically into the fourth inguinal mammary fat pads of *Ptpn1^fl/fl^*, *Lck*-Cre;*Ptpn1^fl/fl^*, or *Lck*-Cre;*Ptpn1^fl/fl^;Stat5^fl/+^* mice and tumor growth, final tumor weights and **f**) the number of tumor-infiltrating CD4^+^ and CD8^+^ T cells were determined. **g**) Intratumoral CD8^+^ T cells were stimulated with PMA/Ionomycin in the presence of Golgi Stop/Plug and stained for intracellular IFN-γ and TNF. Intracellular granzyme B (GrzB) was assessed in unstimulated intratumoral CD8^+^ T cells. Representative results (means ± SEM) from at least two independent experiments are shown. In (a-d; f-g) significance was determined using a one-way ANOVA Test, for tumor sizes in (e) using a 2-way ANOVA Test and for immune cell infiltrates in (c, f) using a 2-tailed Mann-Whitney U Test (^#^p>0.05, ^##^p>0.01).

To determine whether the heightened STAT-5 signaling in PTP1B-deficient T cells may be responsible for the enhanced anti-tumor activity, we implanted AT3-OVA tumor cells into the mammary fat pads of *Ptpn1^fl/fl^* versus *Lck*-Cre;*Ptpn1^fl/fl^* mice, as well as *Lck*-Cre;*Ptpn1^fl/fl^*;*Stat5^fl/+^* mice, where the otherwise enhanced STAT-5 signaling in T cells associated with PTP1B-deficiency was corrected (**Fig. 5e-g**). We found that *Stat5* heterozygosity largely corrected the repression of syngeneic mammary tumor growth and the accumulation of PTP1B-deficient T cells in AT3-OVA mammary tumors (**Fig. 5e-f**). Moreover, *Stat5* heterozygosity completely corrected the enhanced cytotoxic activity of intratumoral CD8^+^ T cells, as assessed by staining for IFNγ, TNF and granzyme B (**Fig. 5g**). Thus, PTP1B deletion enhances the proliferation, activation, cytotoxicity and anti-tumor activity of CD8^+^ T cells by promoting STAT-5 signaling.

### Targeting PTP1B enhances endogenous T cell anti-tumor immunity

To more interrogate more directly PTP1B as a potential therapeutic target for combatting cancer, we next determined if the systemic inhibition of PTP1B with a small molecule might similarly enhance anti-tumor immunity. Trodusquemine (MSI-1436) is a specific allosteric inhibitor of PTP1B ^19,20^ that has been reported to be safe and well-tolerated in humans. Previous studies have shown that at 5-10 mg/kg, MSI-1436 administered intraperitoneally can repress the growth of HER-2^+^ breast cancer xenografts in immunocompromised mice ^19^. Accordingly, we administered C57BL/6 female mice bearing large (50 ^mm2^) AT3-OVA mammary tumors MSI-1436 at 2.5-10 mg/kg (intraperitoneal) every three days and monitored for effects on tumor growth and TILs. At 2.5 mg/kg, MSI-1436 had modest effects on tumour growth, but at 5-10 mg/kg, MSI-1436 markedly suppressed tumor growth (**Fig. 6a, Fig. S24a**). At 5 mg/kg MSI-1436, the repression of tumor growth was accompanied by the infiltration of T cells, NK cells, and DCs that promote anti-tumor immunity, as well as immunosuppressive TAMs, MDSCs and T_regs_ that were also seen in our genetic studies when PTP1B was deleted globally or within the hematopoietic compartment (**Fig. 6b**). The repression of tumor growth at 5 mg/kg MSI-1436 was accompanied by a significant increase in survival so that 33% of mice were alive 29 days post MSI-1436 treatment (**Fig. S24b**). Importantly, the repression of tumor growth by MSI-1436 (5 mg/kg) could not be ascribed to the inhibition of PTP1B in tumor cells as the deletion of PTP1B in AT3-OVA cells using CRISPR-Cas9 genome-editing in itself had no effect on tumor growth (**Fig. S25a-b**). MSI-1436 administration (5 mg/kg) also suppressed the growth of B16F10-OVA melanoma and MC38-OVA colon cancer tumors injected into the flanks of C57BL/6 male mice (**Fig. 6c-d)**. Therefore, the PTP1B inhibitor MSI-1436 can repress the growth of different syngeneic tumors in mice.

**Figure 6.**
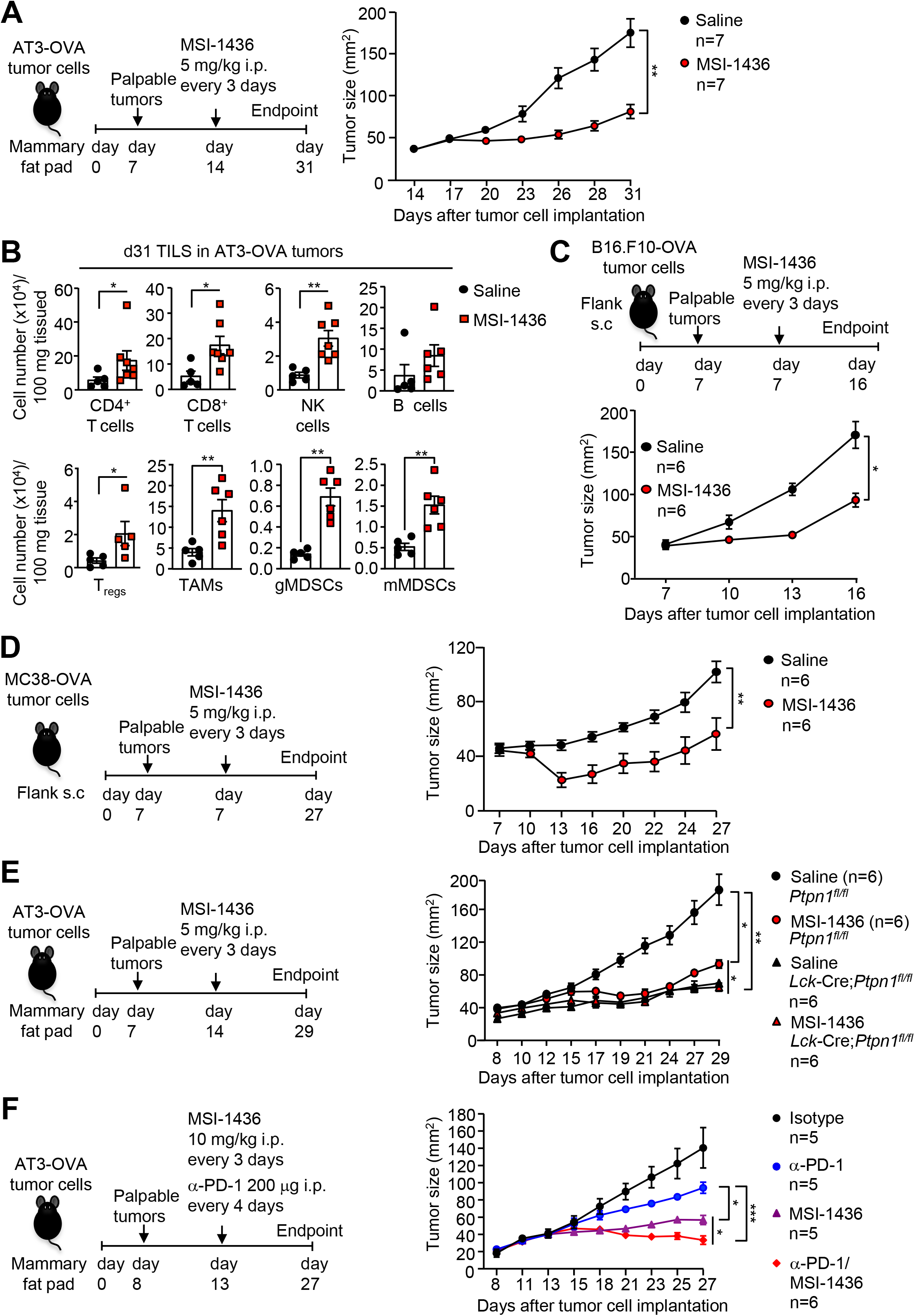
The inhibition of PTP1B in T cells promotes anti-tumor immunity. **a-b**) AT3-OVA mammary tumor cells were implanted into the fourth inguinal mammary fat pads of C57BL/6 mice. Mice were treated with MSI-1436 (5 mg/kg intraperitoneally) or saline on days 14, 17, 20, 23, 26 and 29 after tumor cell implantation. **a**) Tumor growth was monitored and **b**) TILs including CD4^+^, CD8^+^ T cells, CD19^+^ B cells, NK1.1^+^TCRβ^−^ NK cells, CD11c^+^ DCs, CD4^+^CD25^+^FoxP3^+^ regulatory T cells (T_regs)_, CD11b^+^F4/80^hi^Ly6C^−^Ly6G^−^ TAMs, granulocytic CD11b^+^F4/80^hi/lo^Ly6C^int^Ly6G^+^ (gMDSCs) and monocytic CD11b^+^F4/80^hi/lo^Ly6C^+^Ly6G^−^ (mMDSCs) MDSCs were analysed by flow cytometry. **c**) B16F10-OVA or **d**) MC38-OVA tumor cells were xenografted into the flanks of C57BL/6 male mice. Mice were treated every 3 days with saline or MSI-1436 (5 mg/kg i.p.) once tumors (40-50 mm^2^) were established (days 7, 10 and 13 for B16F10-OVA tumors, or days 7, 10, 13, 16 and 19, 22 for MC38-OVA tumors) and tumor growth monitored. **e**) AT3-OVA tumor cells were implanted into the fourth inguinal mammary fat pads of *Ptpn1^fl/fl^* versus *Lck*-Cre;*Ptpn1^fl/fl^* mice. After tumors were established, mice were treated every 3 days with saline or MSI-1436 (5 mg/kg intraperitoneally on days 14, 17, 20, 23, and 26) and tumor growth monitored. **f**) AT3-OVA tumor cells were implanted into the fourth inguinal mammary fat pads of C57BL/6 mice. Mice were treated with MSI-1436 (10 mg/kg intraperitoneally on days 13, 16, 19, 22, and 25) and either α-PD-1 or isotype control (200 μg intraperitoneally in each case on days 13, 17, 21 and 24) alone or MSI-1436 plus α-PD-1. Tumor growth was monitored and tumor weights determined. Representative results (means ± SEM) from at least two independent experiments are shown. Significance for tumor sizes in (a, c-f) was determined using a 2-way ANOVA Test and for tumor-infiltrating lymphocytes in (b) using a 2-tailed Mann-Whitney U Test.

To determine the extent to which the repression of tumor growth by MSI-1436 might be attributable to the inhibition of PTP1B in T cells, we repeated these experiments in control (*Ptpn1^fl/fl^*) versus T cell-specific PTP1B-deficient (*Lck*-Cre;*Ptpn1^fl/fl^*) mice bearing AT3-OVA mammary tumors (**Fig. 6e)**. The deletion of PTP1B in T cells or the administration of MSI-1436 (5 mg/kg) to control mice bearing AT3-OVA tumors similarly repressed tumor growth, but MSI-1436 had no additional effects on tumor growth in T cell-specific PTP1B-deficient mice (**Fig. 6e**). These results are consistent with the effects of MSI-1436 on tumor growth being mediated by the inhibition of PTP1B in T cells. In line with this, we found that the administration of MSI-1436 (5 mg/kg) was accompanied by increased naïve T cells in the spleen, as seen in *Lck*-Cre;*Ptpn1^fl/fl^* mice, as well as increased p-STAT-5 and BCL-2 in naïve, central memory and effector/memory T cells in lymphoid organs (**Fig. S26a-c**). As expected, the administration of MSI-1436 decreased body weight due to reductions in fat mass (**Fig. S27a-b**). This is consistent with PTP1B’s known role in attenuating hypothalamic leptin signaling ^40^ and the ability of MSI-1436 to decrease whole-body adiposity by repressing food intake ^20^. Importantly, MSI-1436 (5 mg/kg) decreased adiposity in both *Ptpn1^fl/fl^* and *Lck*-Cre;*Ptpn1^fl/fl^* mice (**Fig. S27b**) suggesting that the repression of tumor growth was independent of the effects on adiposity. Interestingly, although MSI-1436 treatment and the genetic deletion of PTP1B in T cells similarly repressed tumor growth, MSI-1436 resulted in a greater number of effector/memory T cells within tumors (**Fig. S28**), indicating that MSI-1436, might also elicit T cell-independent effects to affect T cell recruitment and/or activation.

Finally, we determined whether the response to MSI-1436 might be enhanced further by the concomitant blockade of the cell-surface immune checkpoint molecule PD-1. Female mice bearing large AT3-OVA mammary tumors were treated with MSI-1436 (5-10 mg/kg) alone every third day, α-PD-1 alone every fourth day, or MSI-1436 plus α-PD-1, and tumor growth assessed (**Fig. 6f**; **Fig. S29a-b**). At 5 mg/kg MSI-1436 was just as effective, and at 10 mg/kg more effective, at repressing tumor growth than α-PD1 alone, but the addition of α-PD-1 in either case further enhanced the repression of tumor growth (**Fig. 6f**; **Fig. S29a**). Importantly, the combined treatment of MSI-1436 plus α-PD-1 had no additional impact on tumor effector/memory T cell numbers, but decreased the proportion of PD-1^+^Tim3^hi^ CD8^+^ T cells, consistent with the alleviation of T cell exhaustion (**Fig. S29b)**. These findings demonstrate that PTP1B can enhance T cell-mediated anti-tumor immunity and this effect can be enhanced further by targeting the PD-1 checkpoint.

### Targeting PTP1B enhances CAR T cell anti-tumor immunity

CAR T cell therapy has emerged as an exciting immunotherapy approach for non-immunogenic cancers, as it does not rely on endogenous anti-tumor immunity ^41^. Given the marked improvements in T cell-mediated anti-tumor immunity resulting from the deletion or inhibition of PTP1B, we examined whether targeting PTP1B might also enhance the efficacy of adoptively transferred CAR T cells in a solid tumor setting. To this end, we assessed the impact of deleting or inhibiting PTP1B on the activity of second-generation CAR T cells targeting the human orthologue of murine ErbB2/Neu, HER-2 ^42^. The deletion of PTP1B did not affect the generation of CAR T cells *in vitro* (**Fig. S30a**), but enhanced their antigen-specific activation (**Fig. S30b-c**) and their capacity to kill HER-2-expressing 24JK sarcoma cells specifically *in vitro* (**Fig. S30d**). To assess the impact of PTP1B deficiency on the therapeutic efficacy of CAR T cells in *vivo*, we adoptively transferred *Ptpn1^fl/fl^* or *Lck*-Cre;*Ptpn1^fl/fl^* α-HER-2 CAR T cells into sub-lethally irradiated hosts bearing established (25-30 mm^2^) orthotopic tumors arising from the injection of HER-2-expressing E0771 (HER-2-E0771) mammary tumor cells (**Fig. 7a-d**); E0771 mammary tumors resemble human TNBC as they are ER, PR and ErbB2 negative, basal-like and mutant for *Trp53* and are characterised by a robust immunosuppressive tumor microenvironment ^11,43^. HER-2-E0771 cells were grafted into nulliparous human HER-2 transgenic (TG) female mice, where HER-2 expression is driven by the whey acidic protein (WAP) promoter, which induces expression in the lactating mammary gland and the cerebellum ^44^; this ensures that human HER-2-expressing orthotopic tumors are regarded as self, so that host anti-tumour immunity is repressed. Strikingly, whereas the adoptive transfer of *Ptpn1^fl/fl^* control α-HER-2 CAR T cells had little, if any effect on tumor growth, PTP1B-deficient α-HER-2 CAR T cells markedly repressed tumor growth (**Fig. 7a**). This was accompanied by a significant increase in the number of PTP1B-deficient CAR T cells in the tumor, but not in the spleen (**Fig. 7c**), consistent with PTP1B-deficiency driving antigen-induced CAR T cell expansion. Moreover, intratumoral PTP1B-deficient CAR T cells exhibited enhanced cytotoxicity as reflected by the increased expression of TNF, IFNγ and granzyme B (**Fig. 7d**). The enhanced CAR T cell activation/cytotoxicity and repression of tumor growth were accompanied by a significantly enhanced survival of tumor-bearing mice (**Fig. 7b**). Importantly, the adoptive transfer of PTP1B-deficient CAR T cells did not promote systemic inflammation (**Fig. S31a-f)**, as assessed by monitoring for lymphocytic infiltrates in non-lymphoid tissues, including the contralateral mammary glands, lungs and livers (**Fig. S31b**), or for circulating IL-6, IFNγ and TNF (**Fig. S31c**). Moreover, PTP1B-deficient CAR T cells did it promote overt morbidity, as assessed by gross appearance (data not shown) or by measuring food intake, ambulatory activity, voluntary wheel running and energy expenditure (**Fig. S31d**). Furthermore, although the WAP promoter in nulliparous HER-2 TG mice drives HER-2 expression in the cerebellum ^11,44^, there were no overt signs of cerebellar tissue damage (**Fig. S31e**), and the cerebellar control of neuromotor function, as assessed in rotarod tests, was not affected (**Fig. S31f**).

**Figure 7.**
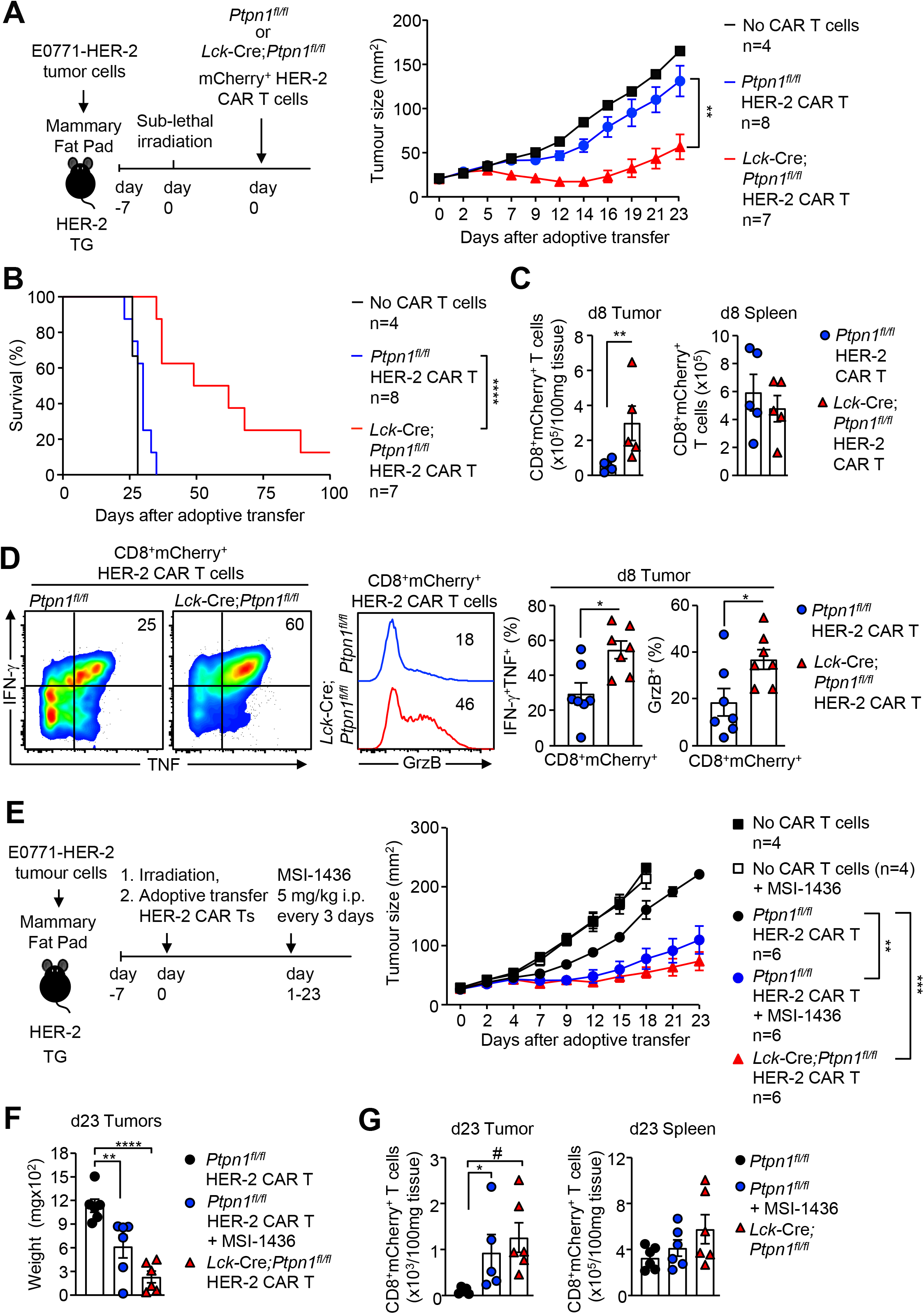
Targeting PTP1B enhances CAR T cell cytotoxicity and efficacy. **a-d**) HER-2-E0771 mammary tumor cells were injected into the fourth inguinal mammary fat pads of female HER-2 TG mice. Seven days after tumor injection HER-2 TG mice received total body irradiation (4 Gy) followed by the adoptive transfer of 20×10^6^ HER-2 CAR T cells generated from *Ptpn1^fl/fl^* or *Lck*-Cre;*Ptpn1^fl/fl^* splenocytes. **a**) Tumor growth and **b**) survival was monitored and **c**) CD45^+^CD8^+^mCherry^+^ CAR T cell infiltrates were determined by flow cytometry at day 8 post adoptive transfer. **d**) Tumor-infiltrating HER-2 CAR T cells were stimulated with PMA/Ionomycin in the presence of Golgi Stop/Plug and stained for intracellular IFN-γ and TNF. Intracellular granzyme B (GrzB) was detected in unstimulated CD8^+^ HER-2 CAR T cell tumor infiltrates. **e-g**) Tumor bearing HER-2 TG mice were treated every 3 days with saline or MSI-1436 (5 mg/kg intraperitoneally) on days 1, 4, 7, 10, 13, 16 and 19 after adoptive HER-2 CAR T cell transfer and tumor growth was monitored and **f**) tumor weights determined. **g**) CD8^+^mCherry^+^ HER-2 CAR T cell infiltrates were determined by flow cytometry at day 23 post adoptive transfer. Representative results (means ± SEM) from at least two independent experiments are shown. Significance for tumor sizes in (a, e) was determined using 2-way ANOVA Test. Significance for tumor-infiltrating lymphocytes and intracellular IFN-γ, TNF and GrzB levels in (c, d) was determined using a 2-tailed Mann-Whitney U Test, for tumor weights in (f) and tumor-infiltrating lymphocytes in (g) using a 1-way ANOVA Test or 2-tailed Mann-Whitney U Test (^#^p<0.05) and in (b) using a Log-rank (Mantel-Cox) test.

To determine whether the inhibition of PTP1B might similarly enhance CAR T cell efficacy, we administered tumor (HER-2-E0771)-bearing HER-2 TG mice that had been sublethally irradiated and then left untreated or treated with control (*Ptpn1^fl/fl^*) or PTP1B-defcient (*Ptpn1^fl/fl^* or *Lck*-Cre;*Ptpn1^fl/fl^*) α-HER-2 CAR T cells with MSI-1436 (5 mg/kg) every three days (**Fig. 7e-g**). In mice not treated with CAR T cells, the inhibitor had little effect, probably because the sublethal irradiation depletes T cells (**Fig. S32**). By contrast, the inhibitor markedly enhanced the ability of α-HER-2 CAR T cells to repress tumor growth, so that they were just as effective as those treated with PTP1B-deficient CAR T cells (**Fig. 7e-f**). The repression of tumor growth in mice treated with MSI-1436 was accompanied by a significant increase in the number of CAR T cells in the tumor (**Fig. 7g**). Taken together, our results demonstrate that targeting PTP1B can not only enhance the anti-tumor activity of endogenous T cells and synergise with the blockade of cell surface checkpoints such as PD-1, but can also enhance the efficacy of adoptively transferred CAR T cells.

## DISCUSSION

Our increasing understanding of how cancer evades the immune system has been instrumental in designing therapies that induce or even reinstate the patient’s immune response to tumor cells. The clinical successes of checkpoint inhibitors directed against PD-1 and cytotoxic T lymphocyte antigen-4 in this regard stem from decades of research focused transferred CAR T cells, without promoting autoimmunity and systemic inflammation.

Our results indicate that the induction of PTP1B in intratumoral T cells in mice occurs as a consequence of the tumor microenvironment, rather than T cell differentiation or activation *per se*, since PTP1B was elevated similarly in both endogenous and adoptively transferred (OT-I) effector/memory CD8^+^ T cells within tumors, when compared to the corresponding T cells in the spleen. Although different aspects of the tumour microenvironment may be responsible for the induction of PTP1B in infiltrating T cells, inflammation is likely to be an important contributing factor. It is well established that inflammation can facilitate malignant growth by both directly promoting the growth of tumor cells and by shaping the immune response. Previous studies have shown that PTP1B expression can be induced by the inflammatory cytokine TNF and via the recruitment of NFκB to the *PTPN1* promoter ^45^. TNF acts by binding to TNFR1, which is expressed widely, and TNFR2, which is expressed on immune cells, including activated T cells. In tumors, TNF may limit T cell responses, since the blockade of TNF-TNFR1 signaling can increase tumor-specific CD8^+^ T cell responses ^46^, whereas monoclonal antibodies targeting TNFR2 enhance T cell-mediated anti-tumor immunity ^47^. It would be interesting to determine the extent to which TNF might be responsible for the induction of PTP1B in intratumoral T cells and whether the benefits of TNFR blockade on anti-tumor immunity ^46,47^ might be reliant on the repression of PTP1B in T cells. Irrespective, we found that the deletion or inhibition of PTP1B enhanced the antigen-induced expansion and activation of naive T cells and the cytotoxicity of effector CD8^+^ T cells to inhibit the growth of syngeneic tumors. Importantly, correcting the increased PTP1B levels in intratumoral T cells in *Lck*-Cre;*Ptpn1^fl/+^* heterozygous mice was sufficient to facilitate the accumulation of effector T cells within tumors and repress tumor growth. As such, our studies are consistent with PTP1B acting as an intracellular checkpoint, whose inhibition can enhance CD8+ T cell-mediated anti-tumor immunity to repress tumor growth.

Our results show that the ability of PTP1B-deficient T cells to repress tumor growth in mice is reliant on the promotion of JAK/STAT-5 signaling. Early studies established that PTP1B could directly dephosphorylate and inactivate JAK-2 and Tyk2 to attenuate cytokine-induced JAK/STAT signaling; there is also evidence that PTP1B can directly dephosphorylate some STAT family members ^29,30,48^. Consistent with this, basal STAT-5 Y694 phosphorylation and cytokine-induced Tyk-2 Y1054/Y1055 and STAT-5 Y694 phosphorylation were enhanced by the deletion of PTP1B. This was attributable to both increased JAK activation and prolonged STAT-5 Y694 phosphorylation, consistent with PTP1B acting directly on both JAKs and STAT-5. The enhanced STAT-5 signaling was evident in both naïve and memory CD4^+^ and CD8^+^ T cells and was accompanied by the induction of STAT-5 transcriptional targets, including T-bet and Eomes, which are important for CD8^+^ T cell cytotoxicity and anti-tumor immunity ^49^, BCL-2, which promotes cell survival, and CD25 (IL2R α chain), which together with IL2Rβ and the γc chain constitute the high affinity receptor for IL2 ^28^. PTP1B-deficiency also enhanced cytokine-induced STAT-5 signaling, especially IL-2-induced STAT-5 signaling in CD8^+^ effector T cells, where IL-2 promotes cytotoxicity by increasing the expression of IFNγ, TNF and granzyme B ^28^. The heightened basal and cytokine-induced STAT-5 signaling were accompanied by an increase in overall thymocyte and T cell numbers, including thymic and peripheral CD4^+^ T_regs_ whose generation and maintenance is reliant on IL-2 ^28^, as well as heightened TCR-induced naive CD4^+^ and CD8^+^ T cell expansion and activation and effector CD8^+^ T cell cytotoxicity and anti-tumor immunity. As in murine T cells, the deletion of PTP1B also increased STAT-5 signaling and the TCR-instigated expansion and activation of human T cells. The enhanced T cell responses and superior tumor control in mice were attenuated when the heightened STAT-5 signaling was corrected by crossing *Lck*-Cre;*Ptpn1^fl/fl^* mice onto a *Stat5^fl/+^* background. Although we cannot exclude that PTP1B might also influence other tyrosine phosphorylation-dependent signaling pathways, our studies indicate that its effects on STAT-5 signaling are nonetheless instrumental in its capacity to act as intracellular checkpoint and influence tumor growth.

We demonstrated that the effects of PTP1B deletion on the T cell-mediated repression of tumor growth phenocopied those in mice in which PTP1B was deleted globally or throughout the hematopoietic compartment. Moreover, we found that the inhibition of PTP1B with MSI-1436 repressed tumor growth by targeting PTP1B in T cells, as its administration had no additional impact in *Lck*-Cre;*Ptpn1^fl/fl^* mice. However, it is possible that in some circumstances, the inhibition PTP1B in tumor cells, or other components of the immune system, might also contribute to the repression of tumor growth. PTP1B is required for the growth and metastasis of HER-2/ErbB2-expressing tumors ^19,22,23^. As such, it would be interesting to determine whether the combined targeting of PTP1B in ErbB2-driven mammary tumors and T cells might yield synergistic outcomes. Furthermore, previous studies have shown that the heterozygous deletion of PTP1B enhances DC maturation and immunogenicity and represses the growth of B cell lymphomas ^25^. Specifically, Eμ-myc-driven B cell lymphomas are attenuated in *Ptpn1^+/–^* mice and this is accompanied by enhanced DC maturation and immunogenicity, as well as repressed T_reg_ and MDSC recruitment ^25^. By contrast, DCs are more tolerogenic in *Ptpn1^−/−^* mice ^26^. We found that repression of syngeneic tumor growth, resulting from the global or hematopoietic-specific deletion of PTP1B, or the systemic inhibition of PTP1B, was accompanied by the recruitment of not only T cells and NK cells, but also DCs, T_regs_ and MDSCs. Although we did not assess DC maturation, any potential effect on DCs appeared to be of limited importance, as the repression of tumor growth was phenocopied in *Lck*-Cre;*Ptpn1^fl/fl^* mice, whereas MSI-1436 neither enhanced nor antagonised the repression of tumor growth in *Lck*-Cre;*Ptpn1^fl/fl^* mice. Nonetheless, MSI-1436 treatment increased intratumoral T cells even in *Lck*-Cre;*Ptpn1^fl/fl^* mice and this could have occurred as a consequence of PTP1B inhibition in DCs to drive licensing and T cell expansion. Therefore, depending on the tumor, there may be additional tumor cell autonomous and non-cell autonomous benefits of systemically targeting PTP1B in cancer.

Our studies indicate that the inhibition PTP1B can be more effective at suppressing tumour growth than the blockade of PD-1. Moreover, we found that combining PTP1B inhibition with PD-1 checkpoint blockade led to a greater tumour repression than either treatment alone. These results suggest that PTP1B acts independently of PD-1 inhibitory signaling. Whereas PD-1 functions to attenuate TCR and potentially co-receptor signaling ^6^, we demonstrate that PTP1B functions to attenuate cytokine signaling, including that mediated by IL-2. Durable clinical responses with PD-1 monotherapy are only evident in a subset of patients, with most developing transient responses or no responses at all ^1,3^. Some studies suggest that inhibitory PD-1 signaling in T cells and CAR T cells can be overcome by promoting IL-2 responses and signaling via STAT-5 ^50,51^. Therefore, targeting PTP1B may afford a means for promoting IL-2/STAT-5 signaling and alleviating PD-1 inhibitory signalling to extend the utility and/or durability of PD-1 checkpoint blockade. In our studies we found that the combined targeting of PTP1B and PD-1 decreased T cell exhaustion, but the precise mechanisms involved remain to be determined.

Beyond the potential to enhance endogenous anti-tumor immunity and the response to anti-PD-1 therapy, our studies also establish the therapeutic potential of targeting PTP1B in adoptive cellular immunotherapy. In particular, we demonstrated that the genetic or pharmacological targeting PTP1B in CAR T cells enhanced the activation and therapeutic efficacy of CAR T cells *in vivo*. To date, CAR T cells have been highly effective in combating hematological malignancies. In particular, CAR T cells targeting the B cell lineage-restricted protein CD19 have transformed the treatment of B cell acute lymphoblastic leukemia ^41^. However, CAR T cells have been largely ineffective against solid tumors ^41^. Herein we demonstrate that targeting PTP1B can enhance the efficacy of adoptively transferred CAR T cells and repress the growth of highly aggressive and otherwise highly immunosuppressive E0771 mammary tumors to significantly extend survival. Notably the inhibition of PTP1B with MSI-1436 was just as effective at enhancing the efficacy of CAR T cells as the genetic deletion of PTP1B. In these studies, the administration of MSI-1436 had no impact on endogenous anti-tumor immunity, since the mice were first lymphodepleted by irradiation to allow for the effective expansion of adoptively transferred CAR T cells. Although lymphodepletion typically is also common in the clinic, an added potential benefit of systemically targeting PTP1B with MSI-1436 might be the alleviation of inhibitory constraints imposed on endogenous tumor-resident T cells, which might serve to prevent the emergence of antigen loss tumor variants during CAR T cell therapy ^52^. Similar outcomes have been sought when CAR T therapies have been combined with PD-1 blockade or oncolytic vaccines ^43,52,53^. Although further studies are required to explore such possibilities, targeting PTP1B with specific inhibitors stands to transform CAR T cell therapy and readily extend the utility of CAR T cells to the effective therapy of recalcitrant solid tumors.

In summary, our studies identify PTP1B as an integral negative regulator of T cell function and an intracellular T cell checkpoint that limits the anti-tumor immunity of tumor-infiltrated CD8^+^ T cells. We demonstrate that the inhibition of PTP1B in cancer can enhance both endogenous T cell-mediated anti-tumor immunity, akin to that seen by targeting the cell surface checkpoint PD-1 that has revolutionised cancer therapy. Furthermore, the inhibition or genetic deletion of PTP1B can overcome a major hurdle that has thus far limited the effectiveness of CAR T cells against solid tumors. Our studies demonstrate that targeting PTP1B with inhibitors such as MSI-1436 may provide an alternative therapeutic strategy for enhancing T cell-mediated anti-tumor immunity to combat cancer.

## ACKNOWLEDGMENTS

We thank Alexandra Ziegler for technical support, Axel Kallies for critical review of the manuscript and DepYmed for providing MSI1436. This work was supported by the National Health and Medical Research Council (NHMRC) of Australia (to T.T.), Cancer Council Victoria (to F.W.) and the National Institutes of Health (CA53840; to N.K.T); P.K.D is a NHMRC Principal Research Fellow.

## AUTHOR CONTRIBUTIONS

Conceptualization: T.T.; Methodology: T.T., F.W., R.D, P.K.D., N.K.T.; Data acquisition: F.W., K-H.L., X.D., M.Z., P.K.G., C.E.X., S.H., S.G. R.X., R.D; Data analysis: F.W., K-H.L., X.D., M.Z., P.K.G., C.E.X., S.H., R.X., K.S., and T.T.; Data interpretation: F.W., K-H.L., M.Z., K.S. R.D., and T.T.; Writing-Original Draft: T.T.; Writing-Review & Editing: F.W., K-H.L., X.D., M.Z., K.S, P.B., P.K.D., R.D., N.K.T., and T.T. Funding acquisition: T.T. and F.W.

## DATA AVAILABILITY STATEMENT

The datasets generated during and/or analysed during the current study are available from the corresponding author on reasonable request.

## METHODS

### Materials

For T cell stimulations, CD3ε (BD Biosciences Cat# 553058, RRID:AB_394591) and CD28 (BD Biosciences Cat# 557393, RRID:AB_396676) antibodies were purchased from BD Biosciences. p-(Y418) SFK (Thermo Fisher Scientific Cat# 14-9034-82) was purchased from Invitrogen. Antibodies to p-(1054/Y1055) TYK2 (Cell Signaling Technology Cat# 9321, RRID:AB_2303972), p-(Y493) ZAP70 (Cell Signaling Technology Cat# 2704, RRID:AB_2217457), p-ERK1/2 (Cell Signaling Technology Cat# 4376, RRID:AB_331772), LCK (Cell Signaling Technology Cat# 2752, RRID:AB_2234649), ZAP-70 (Cell Signaling Technology Cat# 3165, RRID:AB_2218656), ERK1/2 (Cell Signaling Technology Cat# 9102, RRID:AB_330744), and p-(Y694) STAT-5 (D47E7) XP^®^ (Cell Signaling Technology Cat# 4322, RRID:AB_10544692) were from Cell Signaling; pTyr (4G10) antibody from the Monash Antibodies Technologies Facility; anti-actin (Thermo Fisher Scientific Cat# MA5-11866, RRID:AB_10985365) and anti-vinculin (Thermo Fisher Scientific Cat# MA1-25018, RRID:AB_795706) were from Thermo Fisher Scientific, rabbit monoclonal PTP1B (Abcam Cat# ab244207, RRID:AB_2877148) antibody for flow cytometry was from Abcam and goat polyclonal PTP1B (R and D Systems Cat# AF3954, RRID:AB_2174947) antibody for immunoblotting was from R&D Systems. Recombinant human IL-2, murine IL-7 and human IL-15 used for T cell stimulations or the generation of CAR T cells were purchased from the PeproTech. Recombinant murine IFN-α was from BioLegend. *InVivo*MAb anti-mouse PD-1 (Bio X Cell Cat# BE0146, RRID:AB_10949053) and *InVivo*MAb rat IgG2a isotype control (Bio X Cell Cat# BE0251, RRID:AB_2687732) were purchased from Bio X Cell. MSI-1436 was provided by DepYmed Inc, RetroNectin was purchased from Takara, ruxolitinib from Selleckchem and Dnase I from Sigma-Aldrich. The mouse anti-nuclear antibodies Ig’s (total IgA+G+M) ELISA Kit was from Alpha Diagnostic Int., the Transaminase II Kit from Wako Pure Chemicals, the BD Pharmingen APC BrdU Flow Kit and the PE Annexin V Apoptosis Detection Kit II were from BD Biosciences. FBS was purchased from Thermo Scientific, RPMI 1640, DMEM, MEM, HEPES, non-essential amino acids, penicillin/streptomycin and sodium-pyruvate were from Invitrogen and collagenase type IV was purchased from Worthington Biochemical.

### Mice

Mice were maintained on a 12 h light-dark cycle in a temperature-controlled high barrier facility with free access to food and water. 6-12-week-old female B6.SJL-*Ptprc^a^Pepc^b^*/BoyJ (Ly5.1) and human HER-2 transgenic (TG) recipient mice and 6-10-week-old female donor mice were used for adoptive transfers. Aged- and sex-matched littermates were used in all experiments; unless otherwise indicated, female donor mice were used for all in *vivo* experiments, whereas mice of the same sex were used for *ex vivo* experiments. *Ptpn1^fl/fl^* (C57BL/6; MMRRC Cat# 032243-JAX, RRID:MMRRC_032243-JAX), *Lck*-Cre (C57BL/6; IMSR Cat# JAX:006889, RRID:IMSR_JAX:006889), OT-I (IMSR Cat# JAX:003831, RRID:IMSR_JAX:003831), *Stat5^fl/fl^* (C57BL/6; MMRRC Cat# 032053-JAX, RRID:MMRRC_032053-JAX), *Ptpn1^−/−^* (C57BL/6; MMRRC Cat# 032240-JAX, RRID:MMRRC_032240-JAX) and human HER-2 TG (C57BL/6) mice have been described previously ^7,11,17,40^. *Lck*-Cre mice were mated with *Ptpn1^fl/fl^* mice and thereon interbred to generate *Lck*-Cre;*Ptpn1^fl/fl^* mice that were subsequently mated with either OT-1 or *Stat5^fl/fl^* mice to generate OT-I;*Lck*-Cre;*Ptpn1^fl/fl^* or *Lck*-Cre;*Ptpn1^fl/fl^*;*Stat5^fl/+^* mice respectively. Ly5.1 (IMSR Cat# JAX:002014, RRID:IMSR_JAX:002014), C57BL/6J (IMSR Cat# JAX:000664, RRID:IMSR_JAX:000664) mice were purchased from the WEHI Animal Facility (Kew, Australia) or from the Animal Resource Centre (ARC, Perth, Australia).

### Cell lines

The C57BL/6 mouse mammary tumor cell line AT3 (Millipore Cat# SCC178, RRID:CVCL_VR89), the mouse melanoma cell line B16F10 (ATCC Cat# CRL-6475, RRID:CVCL_0159) and the mouse colon carcinoma cell line MC38 (RRID:CVCL_B288) and those engineered to express chicken ovalbumin (AT3-OVA, B16F10-OVA and MC38-OVA) have been described previously ^11,54^. The C57BL/6 mouse breast carcinoma cell line E0771 (ATCC Cat# CRL-3461, RRID:CVCL_GR23), the C57BL/6 mouse sarcoma cell line 24JK and those engineered to express truncated human HER-2 (HER-2-E0771) have been described previously ^11^. All tumor cell lines were maintained in high-glucose DMEM supplemented with 10% (v/v) FBS, L-glutamine (2 mM), penicillin (100 units/mL)/streptomycin (100 μg/mL) and HEPES (10 mM). All tumour cell lines were provided by the PMCC and authenticated for their antigen/marker expression by flow cytometry. 24JK, 24JK-HER-2 and E0771-HER-2 cell lines were received in March 2017. AT3 and AT3-OVA cell lines were received in August 2017. B16F10-OVA and MC38-OVA cell lines were received in April 2020. The B16F10 and MC38 cell lines were received in April 2021. Cells were routinely tested for *Mycoplasma* contamination by PCR and maintained in culture for less than four weeks. All tumour cell lines used came from early frozen batches, between passages two and five after receipt.

### Flow cytometry

Single cell suspensions from thymus, spleen and lymph nodes were obtained as described previously ^7^. For the detection of intracellular cytokines, granzyme B and PTP1B, T cells were fixed and permeabilized with the BD Cytofix/Cytoperm kit according to the manufacturer’s instructions. Briefly, T cells were surface stained with fluorophore-conjugated antibodies and then fixed and permeabilised on ice for 20 min. Cells were stained intracellullarly with the PTP1B monoclonal rabbit antibody (1/200, 45 min, room temperature) which was detected by the secondary antibody against rabbit IgG (H+L) F(ab’)_2_ fragment conjugated to DyLight™ 649 (Jackson ImmunoResearch) (30 min, room temperature). Alternatively, fixed and permeabilised T cells were stained for IFN-γ, TNF and granzyme B for 45 min at room temperature. For the detection of intracellular FoxP3, Eomes and T-bet the Foxp3/Transcription Factor Staining Buffer Set (eBioscience) was used according to the manufacturer’s instructions. Briefly, T cells were fixed and permeabilised for 1 h on ice and intracellular staining was performed at room temperature for 45 min. For the detection of serum cytokines, the BD CBA Mouse Inflammation Kit™ was used according to the manufacturers’ instructions. The BD Pharmingen BrdU Flow Kit from BD Biosciences was used to detect intracellular BrdU in T cells according to the manufacturer’s instructions. The PE Annexin V Apoptosis Detection Kit II from BD Biosciences was used to detect apoptotic T cells according to the manufacturer’s instructions.

Cells were stained with the specified antibodies on ice for 30 min and analyzed using a LSRII, Fortessa, Symphony (BD Biosciences) or CyAn™ ADP (Beckman Coulter). For FACS sorting, cells were stained for 30 min on ice and purified using either a BD Influx cell sorter, or the BD FACSAria II, BD FACSAria Fusion 3 or BD FACSAria Fusion 5 instruments. Data was analyzed using FlowJo10 (Tree Star Inc.) software. For cell quantification, a known number of Calibrite™ Beads (BD Biosciences) or Nile Red Beads (Prositech) or Flow-Count Fluorospheres (Beckman Coulter) were added to samples before analysis. Dead cells were excluded with propidium iodide (1 μg/ml; Sigma-Aldrich). Paraformaldehyde fixed dead cells were excluded with the LIVE/DEAD™ Fixable Near-IR Dead Cell Stain Kit (Thermo Fisher Scientific).

The following antibodies from BD Biosciences, BioLegend, Invitrogen or eBioscience were used for flow cytometry: Phycoerythrin (PE) or peridinin-chlorophyll cyanine 5.5 (PerCP-Cy5.5)-conjugated CD3 (BD Biosciences Cat# 553063, RRID:AB_394596; BD Biosciences Cat# 551163, RRID:AB_394082); PE, BUV805, PerCP-Cy5.5 or phycoerythrin-cyanine 7 (PE-Cy7)-conjugated CD4 (BD Biosciences Cat# 553049, RRID:AB_394585; BD Biosciences Cat# 741912, RRID:AB_2871226; BD Biosciences Cat# 550954, RRID:AB_393977; BD Biosciences Cat# 561099, RRID:AB_2034007); BV711, Pacific Blue-conjugated (PB), allophycocyanin (APC)-Cy7 or APC-conjugated CD8 (BioLegend Cat# 100759, RRID:AB_2563510; BD Biosciences Cat# 558106, RRID:AB_397029; BD Biosciences Cat# 557654, RRID:AB_396769; BD Biosciences Cat# 553035, RRID:AB_398527); BV711-conjugated CD11b (BioLegend Cat# 101241, RRID:AB_11218791); APC-conjugated CD11c (BioLegend Cat# 117309, RRID:AB_313778); PE-Cy7-conjugated CD19 (BD Biosciences Cat# 552854, RRID:AB_394495); PerCP-Cy5.5, PE or APC-Cy7-conjugated CD25 (BD Biosciences Cat# 551071, RRID:AB_394031; BD Biosciences Cat# 553866, RRID:AB_395101; BD Biosciences Cat# 561038, RRID:AB_2034002); Fluorescein isothiocyanate (FITC), V450, BUV737, BV786, Alexa Fluor 700 or PE-Cy7-conjugated CD44 (BD Biosciences Cat# 553133, RRID:AB_2076224; BD Biosciences Cat# 560451, RRID:AB_1645273; BD Biosciences Cat# 612799, RRID:AB_2870126; BD Biosciences Cat# 563736, RRID:AB_2738395; BD Biosciences Cat# 560567, RRID:AB_1727480; BD Biosciences Cat# 560569, RRID:AB_1727484); APC-conjugated CD45 (Thermo Fisher Scientific Cat# 17-0451-83, RRID:AB_469393); BUV395, FITC, V450, APC or PE-conjugated CD45.1 (BD Biosciences Cat# 565212, RRID:AB_2722493; BD Biosciences Cat# 553775, RRID:AB_395043; BD Biosciences Cat# 560520, RRID:AB_1727490; BD Biosciences Cat# 558701, RRID:AB_1645214; BD Biosciences Cat# 553776, RRID:AB_395044); FITC, PE or PerCP-Cy5.5-conjugated CD45.2 (BD Biosciences Cat# 553772, RRID:AB_395041; BD Biosciences Cat# 560695, RRID:AB_1727493; BD Biosciences Cat# 561096, RRID:AB_2034008); biotinylated CD49d (BD Biosciences Cat# 557419, RRID:AB_396692); APC, BV421, BV786 or PE-conjugated CD62L (BD Biosciences Cat# 553152, RRID:AB_398533; BioLegend Cat# 104435, RRID:AB_10900082; BD Biosciences Cat# 564109, RRID:AB_2738598; BD Biosciences Cat# 553151, RRID:AB_394666); V450-conjugated CD69 (BD Biosciences Cat# 560690, RRID:AB_1727511); FITC or PE-Cy7-conjugated CD279 (BioLegend Cat# 135213, RRID:AB_10689633; BioLegend Cat# 135216, RRID:AB_10689635); PE-Cy7 or BV711-conjugated CD366 (Thermo Fisher Scientific Cat# 25-5870-80, RRID:AB_2573482; BioLegend Cat# 119727, RRID:AB_2716208); FITC or APC-Cy7 conjugated B220 (BD Biosciences Cat# 553088, RRID:AB_394618; BD Biosciences Cat# 552094, RRID:AB_394335); Brilliant Violet 421 (BV421)-conjugated TCR-β (H57-597; BioLegend Cat# 109229, RRID:AB_10933263); PE-Cy7-conjugated NK1.1 (PK136; BD Biosciences Cat# 552878, RRID:AB_394507); APC-Cy7-conjugated Ly6C (BD Biosciences Cat# 560596, RRID:AB_1727555); PE-conjugated Ly6G (BD Biosciences Cat# 551461, RRID:AB_394208); APC or PE-conjugated F4/80 (BD Biosciences Cat# 566787, RRID:AB_2869866; BD Biosciences Cat# 565410, RRID:AB_2687527); APC-conjugated KLRG1 (Thermo Fisher Scientific Cat# 17-5893-82, RRID:AB_469469); PE-Cy7-conjugated IFNγ (BD Biosciences Cat# 557649, RRID:AB_396766); FITC or APC-conjugated TNF (BD Biosciences Cat# 554418, RRID:AB_395379; BD Biosciences Cat# 561062, RRID:AB_2034022); AlexaFluor 647-conjugated Granzyme B (BioLegend Cat# 515405, RRID:AB_2294995), APC-conjugated T-bet (BioLegend Cat# 644813, RRID:AB_10896913); Alexa Fluor 488-conjugated Eomes (Thermo Fisher Scientific Cat# 53-4875-80, RRID:AB_10853025) and Alexa Fluor 647-conjugated BCL-2 (BioLegend Cat# 633510, RRID:AB_2274702). PE-Cy7-conjugated Streptavidin was used to detect biotinylated CD49d (9C10).

### TCR signaling

Naïve (CD44^lo^CD62L^hi^) CD8^+^ T cells were purified using the EasySep™ Mouse Naïve CD8^+^ T cell Isolation Kit (Stemcell Technologies) according to the manufacturer’s instructions. Naïve CD8^+^ T cells were incubated with anti-CD3ε (5 μg/ml) for 15 min on ice in RPMI 1640 supplemented with 1% (v/v) FBS. Cells were washed with RPMI 1640 and then incubated with 20 μg/ml rabbit anti-hamster IgG (H+L) (Sigma-Aldrich) in RPMI 1640 for the indicated times at 37°C. Cells were washed with ice-cold PBS and lysed in ice-cold RIPA lysis buffer [50 mM HEPES pH 7.4, 1% (v/v) Triton X-100, 1% (vol/vol) sodium deoxycholate, 0.1% (v/v) SDS, 150 mM NaCl, 10% (v/v) glycerol, 1.5 mM MgCl2, 1 mM EGTA, 50 mM sodium fluoride, 1 µg/ml pepstatin A, 5 μg/ml aprotinin, 1 mM benzamadine, 1 mM phenylmethysulfonyl fluoride, 1 mM sodium vanadate], clarified by centrifugation (16,000 x g, 10 min, 4°C) and processed for immunoblotting.

To assess intracellular ERK1/2 T185Y187 phosphorylation (p-ERK1/2) by flow cytometry 3×10^6^ T cells isolated from inguinal, brachial, axillary, superficial cervical and deep cervical lymph nodes were incubated with anti-CD3ε (1 or 5 μg/ml) for 15 min on ice in RPMI 1640 supplemented with 1% (v/v) FBS. Cells were washed with RPMI 1640 and then incubated with 20 μg/ml rabbit anti-hamster IgG (H+L) (Sigma-Aldrich) in RPMI 1640 for the indicated times at 37°C. Cells were stained with fluorophore-conjugated antibodies against CD4, CD8, CD44 and CD62L, fixed in 2% (w/v) paraformaldehyde, permeabilised with 90% (v/v) methanol and then intracellularly stained with anti-phospho-p44/42 MAPK (Erk1/2; Cell Signaling) for 30 min at room temperature and then with secondary antibodies against rabbit IgG (H+L) F(ab’)_2_ fragment conjugated to DyLight™ 649 (Jackson ImmunoResearch). p-ERK1/2 mean fluorescence intensities (MFIs) in CD4^+^ and CD8^+^ T cell subsets were analysed by flow cytometry.

To assess calcium flux after TCR stimulation, T cells isolated from inguinal, brachial, axillary, superficial cervical and deep cervical lymph nodes were stained with fluorophore-conjugated antibodies against CD8, CD44 and CD62L and then incubated with 5 μM Fluo-4 AM (Molecular Probes) for 15 min at 37°C. T cells were washed three times in complete T cell medium [RPMI 1640 supplemented with 10% (v/v) FBS, L-glutamine (2 mM), penicillin (100 units/ml)/streptomycin (100 μg/ml), MEM non-essential amino acids (0.1 mM), sodium-pyruvate (1 mM), HEPES (10 mM) and 2-β-mercaptoethanol (50 μM)] and incubated then with anti-CD3ε (1 μg/ml) for 15 min on ice in RPMI 1640 supplemented with 1% (v/v) FBS. Cells were washed with RPMI 1640 supplemented with 1% (v/v) FBS, 20 μg/ml rabbit anti-hamster IgG (H+L) (Sigma-Aldrich) added and the median calcium response at 37°C acquired immediately using a BD FACSAria II. Peak time, slope and peak intensity of the calcium response in CD8^+^ T cell subsets were calculated using FlowJo10 (Tree Star Inc.) software.

### Cytokine signaling

For assessing IL-15-induced TYK2 and STAT-5 phosphorylation in CD8^+^ central memory T cells, total CD8^+^ T cells were first enriched using the EasySep™ Mouse CD8^+^ T cell isolation kit (Stemcell Technologies) and then CD8^+^CD44^hi^CD62L^lo^ central memory T cells purified by FACS. Purified central memory T cells (4 x 10^5^) in RPMI plus 1% (v/v) FBS were stimulated with human IL-15 (50 ng/ml) for the indicated times at 37°C. Cells were washed with ice-cold PBS and lysed in RIPA lysis buffer, clarified by centrifugation (16,000 x g, 10 min, 4°C) and processed for immunoblotting.

For assessing IL-2-induced TYK2 and STAT-5 phosphorylation in *in vitro* generated CD8^+^ rested effector T cells by immunoblotting, naïve (CD44^lo^CD62L^hi^) CD8^+^ T cells were purified using the EasySep™ Mouse Naïve CD8^+^ T cell Isolation Kit (Stemcell Technologies) and stimulated with plate-bound α-CD3 (5 μg/ml) and α-CD28 (5 μg/ml) for 72 h at 37°C in complete T cell medium in the presence of human IL-2 (10 ng/ml), washed and then further expanded for two additional days in the presence of human IL-2 (10 ng/ml) and murine IL-7 (0.2 ng/ml). Activated CD8^+^ T cells were then rested overnight in RPMI 1640 supplemented with 1% (v/v) FBS in the presence of murine IL-7 (0.2 ng/ml). Rested effector CD8^+^ T cells were stimulated with human IL-2 (5 ng/ml) for the indicated times at 37°C. Cells were washed with ice-cold PBS, lysed in RIPA lysis buffer, clarified by centrifugation (16,000 x g, 10 min, 4°C) and processed for immunoblotting.

For IL-2 pulse/chase experiments, *in vitro* generated CD8^+^ rested effector T cells were serum starved in RPMI 1640 supplemented with 0.1% (v/v) FBS for 3 h, pulsed with 5 ng/ml human IL-2 for 10 min at 37°C, washed and then incubated in RPMI 1640 supplemented with 0.1% FBS with or without ruxolitinib (250 nM) for the indicated times at 37°C. Cells were twashed with iced-cold PBS, lysed in RIPA lysis buffer, clarified by centrifugation (16,000 x g, 10 min, 4°C) and processed for immunoblotting.

For assessing IL-2-induced STAT-5 phosphorylation in CD8^+^ ‘rested effector’ cells by flow cytometry, T cells isolated from inguinal, brachial, axillary, superficial cervical and deep cervical lymph nodes were stimulated with plate-bound α-CD3 (5 μg/ml) and α-CD28 (5 μg/ml) for 72 h at 37°C in the presence of human IL-2 (10 ng/ml). Activated CD8^+^ T cells were rested overnight in RPMI 1640 supplemented with 1% (v/v) FBS. Rested effector CD8^+^ T cells were then stimulated with human IL-2 (5 ng/ml) for the indicated time points at 37°C. T cells were stained with fluorophore-conjugated antibodies against CD4 and CD8, fixed in 2% (w/v) paraformaldehyde, permeabilised with acetone/methanol (50:50) and then stained intracellularly with anti-p-(Y694) STAT-5 (D47E7) XP^®^ followed by secondary antibodies against rabbit IgG (H+L) F(ab’)_2_ fragment conjugated to DyLight™ 649 (Jackson ImmunoResearch). p-ERK1/2 MFIs in rested effector CD8^+^ T cells were analysed by flow cytometry.

### T cell proliferation

T cells isolated from inguinal, brachial, axillary, superficial cervical and deep cervical lymph nodes were stained with fluorochrome-conjugated antibodies for CD8, CD62L and CD44 and naïve CD8^+^CD44^lo^CD62L^hi^ T cells purified by FACS. For assessing *ex vivo* T cell proliferation by CellTrace™ Violet (CTV; Molecular Probes, Thermo Fisher Scientific) dilution, T cells were incubated with CTV in D-PBS at a final concentration of 2 μM for 10 min at 37°C. Cells were then washed three times with D-PBS supplemented with 10% (v/v) FBS. T cells (2 x 10^5^) were incubated with the indicated amounts of α-CD3 and α-CD28 coated onto 96-well round bottom plates for 3 days in complete T cell medium at 37°C and proliferation determined by assessing CTV dilution by flow cytometry. For cell quantification, a known number of Calibrite™ Beads (BD Biosciences) or Nile Red Beads (Prositech) or Flow-Count Fluorospheres (Beckman Coulter) were added to samples before analysis.

For assessing OT-I T cell proliferation *in vivo*, FACS-purified naïve OT-I^+^ CD8^+^CD44^lo^CD62L^hi^ T cells were labelled with 5 μM CTV and adoptively transferred (2 x 10^6^) into female Ly5.1 mice bearing established AT3-OVA tumors (40-50 mm^2^). On day 8 post adoptive transfer, T cells were harvested from draining lymph nodes and stained with fluorophore-conjugated antibodies against CD8, CD45.1 (Ly5.1) and CD45.2 (Ly5.2) and the proliferation of CD8^+^Ly5.1^−^Ly5.2^+^ T cells was determined by flow cytometry.

### Tumor studies

Male mice were injected subcutaneously with 1 x 10^5^ B16F10 or B16F10-OVA cells or 5 x 10^5^ MC38 or MC38-OVA tumor cells resuspended in 100 μl D-PBS into the flank. Alternatively, female mice were injected orthotopically with 5 x 10^5^ AT3 or AT3-OVA cells resuspended in 20 μl D-PBS into the fourth mammary fat pad.

For CAR T cell studies, female human HER-2 TG mice were injected orthotopically with 2 x 10^5^ HER-2-E0771 cells resuspended in 20 μl D-PBS into the fourth mammary fat pad. At day 7 post tumor cell injection, human HER-2 TG mice were pre-conditioned with total-body irradiation (4 Gy) prior to the adoptive transfer of 20 × 10^6^ CAR T cells. Tumor sizes were measured in two dimensions (mm^2^) using electronic callipers three times per week as soon as tumors were palpable (20-25 mm^2^). Mice were sacrificed at the ethical experimental endpoint (200 mm^2^ tumor size) and individual tumors were removed to measure tumor weights or to assess T cell infiltrates by flow cytometry.

### Generation of murine CAR T cells

Murine CAR T cells were generated as previously described ^12^. Briefly, splenocytes were cultured overnight with anti-CD3ε (0.5 μg/ml) and anti-CD28 (0.5 μg/ml) in the presence of 10 ng/ml human IL-2 and 0.2 ng/ml murine IL-7 in complete T cell medium. Retroviruses encoding a second-generation CAR targeting HER-2 was obtained from the supernatant of the GP+E86 packaging line spun together with T cells onto RetroNectin-coated (10 μg/ml) 6-well plates (Takara Bio) and incubated overnight before the second viral transduction. T cells were maintained in IL-2/IL-7-containing complete media and used 7-8 days after transduction.

### CAR T cell cytotoxicity assays

For CAR T cell cytotoxicity assays, HER-2 expressing 24JK sarcoma cells (24JK-HER-2) and HER-2 negative 24JK sarcoma cells were labelled with 5 mM or 0.5 mM CellTrace™ Violet (CTV; Invitrogen-Molecular Probes) respectively in D-PBS supplemented with 0.1% (v/v) BSA for 15 min at 37°C. Tumor cells were then washed three times with D-PBS supplemented with 10% (v/v) FBS and mixed at a 1:1 ratio. HER-2-specific CAR T cells were added at different concentrations to HER-2-expressing (5 x 10^4^) and HER-2 negative (5 x 10^4^) 24JK cells and incubated for 4 h at 37°C in complete T cell medium. Antigen-specific target cell lysis, assessed by the specific depletion of CTV^hi^ 24JK-HER-2 cells, but not CTV^dim^ 24JK cells, was monitored by flow cytometry.

### PTP1B-inhibition and immune checkpoint blockade

Mice with established tumors (40-50 mm^2^) where randomly grouped to receive 0.9% (w/v) saline only or MSI-1436 dissolved in 0.9% (w/v) saline. MSI-1436 was administered at 5 or 10 mg/kg body weight in 100 μl every third day intraperitoneally for up to 29 days. Control animals received an injection at the same time of 100 μl of saline. Solutions were freshly prepared before each administration. Mice were sacrificed at the experimental endpoint (200 mm^2^ tumor size) and individual tumors were removed to measure tumor weights or to assess T cell infiltrates by flow cytometry.

To assess the combined effects of PTP1B-inhibition with MSI-1436 and immune checkpoint blockade with anti-PD-1, tumor-bearing mice (40-50 mm^2^) were treated with MSI-1436 as described above every third day. In addition, mice received four intraperitoneal injections of anti-PD-1 (200 μg in 200 μl D-PBS; clone RMP1-14), or rat IgG2a isotype control (100 μg in 200 μl D-PBS, Bio X Cell) every four days for up to 27 days. Mice were sacrificed at the experimental endpoint (200 mm^2^ tumor size) and individual tumors were removed to measure tumor weights or to assess T cell infiltrates by flow cytometry.

### Adoptive OT-I T cell therapy

Female Ly5.1 mice were injected orthotopically with 5×10^5^ AT3-OVA cells resuspended in 20 μl D-PBS into the fourth mammary fat pad. At day 7-12 post tumor injection, mice with established tumors (40-50 mm^2^) where randomly grouped and 2 x 10^6^ naïve Ly5.2^+^CD8^+^CD44^lo^CD62L^hi^ OT-I T cells [purified by FACS or using the EasySep™ murine naïve CD8^+^ T cell isolation kit (Stemcell Technologies)] were injected intravenously. Mice were sacrificed at the indicated times and Ly5.1^−^Ly5.2^+^CD8^+^ OT-I T cell infiltrates in draining lymph nodes and tumors were determined by flow cytometry.

### Analysis of tumor-infiltrating T cells

Tumor-bearing mice were sacrificed and tumors excised and digested at 37°C for 30 min using a cocktail of 1 mg/ml collagenase type IV (Worthington Biochemicals) and 0.02 mg/ml DNase (Sigma-Aldrich) in DMEM supplemented with 2% (v/v) FBS. Cells were passed through a 70 μm cell strainer (BD Biosciences) twice and processed for flow cytometry. For the assessment of activated and exhausted T cells, cells were surface stained with fluorophore-conjugated antibodies against CD62L, CD44, PD-1 and TIM-3. For the detection of intracellular cytokines TNF and IFNγ, T cells were stimulated with PMA (20 ng/ml; Sigma-Aldrich) and ionomycin (1 μg/ml; Sigma-Aldrich) in the presence of GolgiPlug™ and GolgiStop™ (BD Biosciences) for 4 h at 37°C in complete T cell medium. For the intracellular detection of granzyme B, cells were processed untreated without the addition of GolgiPlug™ and GolgiStop™.

### Quantitative real-time PCR

RNA was extracted with TRIzol reagent (Thermo Fisher Scientific) and RNA quality and quantity determined using a NanoDrop 2000 (Thermo Fisher Scientific). mRNA was reverse transcribed with the High-Capacity cDNA Reverse Transcription Kit (Applied Biosystems) and processed for quantitative real-time PCR using the Fast SYBR™ Green Master Mix (Applied Biosystems). Primer sets from PrimePCR™ SYBR® Green Assay (Bio-Rad, #10025636) were utilized to perform quantitative PCR detecting *Ptpn1* (Bio-Rad PrimePCR™ SYBR^®^ Green Assay: qMmuCED0045967). Relative gene expression (ΔΔCt) was determined by normalisation to the house-keeping gene *Rps18*.

### Intracellular staining for p-STAT-5, BCL-2, BCL-xL in human T cells

Cells fixed in 4% (v/v) paraformaldehyde, permeabilised with methanol/acetone (50/50) and then stained intracellularly by incubating cells with rabbit antibody against p-(Y694) STAT-5 (D47E7) XP (Cell Signaling), mouse antibody against BCL-2 (100, Santa Cruz Biotechnology) or mouse antibody against BCL-xL (H5, Santa Cruz Biotechnology) for 60 min and then secondary antibodies against rabbit IgG (H+L) F(ab’)_2_ fragment conjugated to DyLight™ 649 (Jackson ImmunoResearch) or mouse IgG (H+L) conjugated to Alexa Fluor 647 (Thermo Fisher). Cells were then stained with anti-human CD8-FITC (BW135/80, Miltenyi biotec) and anti-human CD4-PE-Cy7 (SK3, BD Biosciences) and mean fluorescence intensities (MFIs) for p-(Y694) STAT5, BCL-xL and BCL-2 in CD4^+^ and CD8^+^ T cell subsets were determined by flow cytometry.

### Statistical analyses

Statistical analyses were performed with Graphpad Prism software 9 using the non-parametric using 2-tailed Mann-Whitney U Test, the parametric 2-tailed Student’s t test, the one-way or 2-way ANOVA-test using Kruskal-Wallis, Turkey, Bonferroni or Sidak post-hoc comparison or the Log-rank (Mantel-Cox test) where indicated. *p<0.05, **p<0.01, ***p<0.001 and ****p<0.0001 were considered as significant.

### Human and animal ethics

Human ethics approval for use of PBMCs was granted by both the Red Cross and the Peter MacCallum Cancer Centre Human Research and Ethics committee (HREC# 01/14). Informed consent was obtained from blood donors by the Red Cross.

All animal experiments were performed in accordance with the NHMRC Australian Code of Practice for the Care and Use of Animals. All protocols were approved by the Monash University School of Biomedical Sciences Animal Ethics Committee (Ethics numbers: MARP/2012/124, 23177 and 27792) or the Peter MacCallum Animal Ethics and Experimentation Committee (Ethics number: E604).

