## Supplementary material for "PTP1B is an intracellular checkpoint that limits T cell and CAR T cell anti-tumor immunity": Supp Methods and Data

***CRISPR RNP gene-editing***

PTP1B was deleted in AT3-OVA cells using CRISPR ribonucleoprotein (RNP)-based gene-editing. AT3-OVA tumor cells were transfected with recombinant Cas9 (74 pmol; Alt-R S.p. Cas9 Nuclease V3, IDT) pre-complexed with either short guide (sg) RNAs (600 pmol) targeting *Ptpn1* (5'UUUCAGUUGACCACAGU3'), or non-targeting sgRNAs (5'GCACUACCAGAGCUAACUCA3') as a control, using the P3 Primary Cell 4D-Nucleofector X™ Kit (Lonza Bioscience) and cultured in high-glucose DMEM supplemented with 10% (v/v) FBS, L-glutamine (2 mM), penicillin (100 units/mL)/streptomycin (100 μg/mL) and HEPES (10 mM).

For deleting PTP1B in human T cells, human peripheral blood mononuclear cells (PBMCs) were isolated from normal buffy coats using Ficoll centrifugation and cultured in complete human T cell medium [RPMI 1640 (Gibco Life Technologies) supplemented with 10% heat-inactivated FBS, L-glutamine (2 mM), MEM non-essential amino acids (0.1 mM), sodium-pyruvate (1 mM), 10 mM 4-(2-hydroxyethyl)-1-piperazineethanesulfonic acid (HEPES), 100 U/mL penicillin and 100 μg/mL streptomycin ] supplemented with human IL-2 (300 IU/ml) and stimulated with soluble anti-human CD3 (OKT3; 30 ng/ml, eBioscience) for 3 days at 37°C. Activated human T cells were then electroporated, as described for AT3-OVA cells, with recombinant Cas9 pre-complexed with either sgRNAs targeting human *PTPN1* (5'UAAAAAUGGAAGAAGCCCAA3') or non-targeting sgRNAs (5'GCACUACCA GAGCUAACUCA3') using the P3 Primary Cell 4D-Nucleofector X™ Kit according to the manufacturer's instructions.

***Human T cell activation***

To monitor for the TCR-mediated activation of human T cells, control or PTP1B-deleted T cells were cultured with plate-bound α-human CD3 (OKT3; 1 μg/ml, eBioscience) in complete human T cell media overnight and then stained with anti-human CD8-FITC (BW135/80, Miltenyi Biotec), anti-human CD4-PE-Cy7 (SK3, BD Biosciences) and anti-human CD69-APC (FN50, BD Biosciences) and analysed by flow cytometry. For the assessment of T cell proliferation by CellTrace™ Violet dilution, T cells were incubated with CTV in D-PBS supplemented with BSA (0.1%) at a final concentration of 2 μM for 10 min at 37ºC. Cells were then washed three times with D-PBS supplemented with 10% (v/v) FBS. Washed T cells (1 x $10^5$) were incubated with plate-bound α-human CD3 (0-1 μg/ml) antibody for 5 days at 37°C and then harvested and stained with anti-human CD8-FITC (BW135/80, Miltenyi Biotec) and anti-human CD4-PE-Cy7 (SK3, BD Biosciences) and proliferation assessed by monitoring for CTV dilution by flow cytometry.

### *Histology and immunohistochemistry*

For immunohistochemistry, tumors were formalin-fixed and paraffin-embedded. Sections were dewaxed in Histopure (CSA Pathology) for 3 x 5 min, dehydrated in ethanol (3 x 5 min) and antigens retrieved in Tris/EDTA pH 8.0 buffer for 5 min in a pressure cooker (70 kpa). Endogenous peroxidase activity was blocked in 3% $H_2O_2$ for 10 min and sections blocked with 1.5% (v/v) horse (for Ki67) or goat (for cleaved caspase 3) serum in phosphate-buffered saline (PBS) for 30 min at room temperature and incubated overnight (4°C) with primary antibodies anti-Ki67 (1:400; Clone 8D5, Cell Signaling) or anti-cleaved caspase 3 (Asp175) (1:2000; Clone 5A1E, Cell Signaling). Ki67- or cleaved caspase 3-positive cells were visualised using mouse (Ki67) or rabbit (cleaved caspase 3) IgG VECTORSTAIN ABC Elite and DAB (3,3′-diaminobenzidine) Peroxidase Substrate Kits (Vector Laboratories, UK) and counterstained with haematoxylin. TUNEL stauning was performed using the DeadEnd™Colorimetric TUNEL System (Promega) following the manufacturer's instructions.

Sections were visualised on an Olympus CX33 microscope (Olympus) and imaged at 20x magnification.

For immunofluorescence microscopy, tumors were formalin-fixed and paraffin-embedded. Sections were dewaxed in Histopure (CSA Pathology) for 3 x 5 min, dehydrated in ethanol (3 x 5 min) and antigens retrieved in Leica EDTA epitope pH 9 retrieval buffer (Leica Biosystems) for 5 min in a pressure cooker (70 kpa). Sections were blocked in 5% normal goat serum in Tris-buffered saline (TBS) for 60 min at room temperature and incubated overnight (4°C) with endomucin primary antibody (1:200; eBioscience, Thermo Fisher Scientific) and then with goat anti-rat IgG Alexa Fluor 555 (Invitrogen, Thermo Fisher Scientific) secondary antibody for 60 min at room temperature. Nuclei were counterstained with DAPI before mounting in Fluoromount-G mounting medium (Thermo Fisher Scientific). Fluorescence was visualised on a Zeiss Axioskop-2 Mot Plus fluorescence microscope (Zeiss) and 20x magnification.

For brain histology, mice were transcardially perfused with heparinized saline [10,000 units/l heparin in 0.9% (w/v) NaCl] followed by 4% (w/v) paraformaldehyde in phosphate buffered saline (PBS; 0.1 M, pH 7.4). Brains were post-fixed overnight at 4°C and then kept refrigerated for three days in 30% (w/v) sucrose in 0.1 M PBS to cryoprotect the tissue, before freezing on dry ice. Series of 30 μm sections (180 μm apart) were cut in the coronal plane (Leica CM1850 cryostat, Leica Biosystems) throughout the entire rostral-caudal extent of the cerebellum. A representative subset of sections were stained with hematoxylin and eosin, mounted with dibutylphthalate polystyrene xylene mounting media, and imaged on an Olympus CX33 microscope (Olympus) at 20x magnification.

***Behavioural assessments***

Food intake, ambulatory activity, wheel running and energy expenditure were assessed over 48 h after 24 h acclimation using a Promethion Metabolic Screening System (Sable Systems International, NV) fitted with indirect open circuit calorimetry, running wheels and food consumption and activity monitors.

Neuromotor function was assessed with rotarod tests. Mice were trained once per day for 4 days on a rotating rod with a lane width of 5 cm (Ugo Basile Rota-Rod 476000, France) spinning at 4 rpm for 5 min. After training, mice were subjected to an incremental protocol, where the speed was increased over 480s from 4 to 60 rpm. All animals were subjected to four independent trials separated by 1 h and the latency to fall (length of time that the mice remained on the rod) recorded and analysed.

**SUPPLEMENTAL DATA**

***PTP1B is an intracellular checkpoint that limits T cell and CAR T cell anti-tumor immunity***

Florian Wiede, Kun-Hui Lu , Xin Du, Mara N. Zeissig, Rachel Xu, Pei Kee Goh, Chrysovalantou E. Xirouchaki, Samuel J. Hogarth, Spencer Greatorex, Kevin Sek, Roger J. Daly, Paul A. Beavis, Phillip K. Darcy, Nicholas K. Tonks, Tony Tiganis

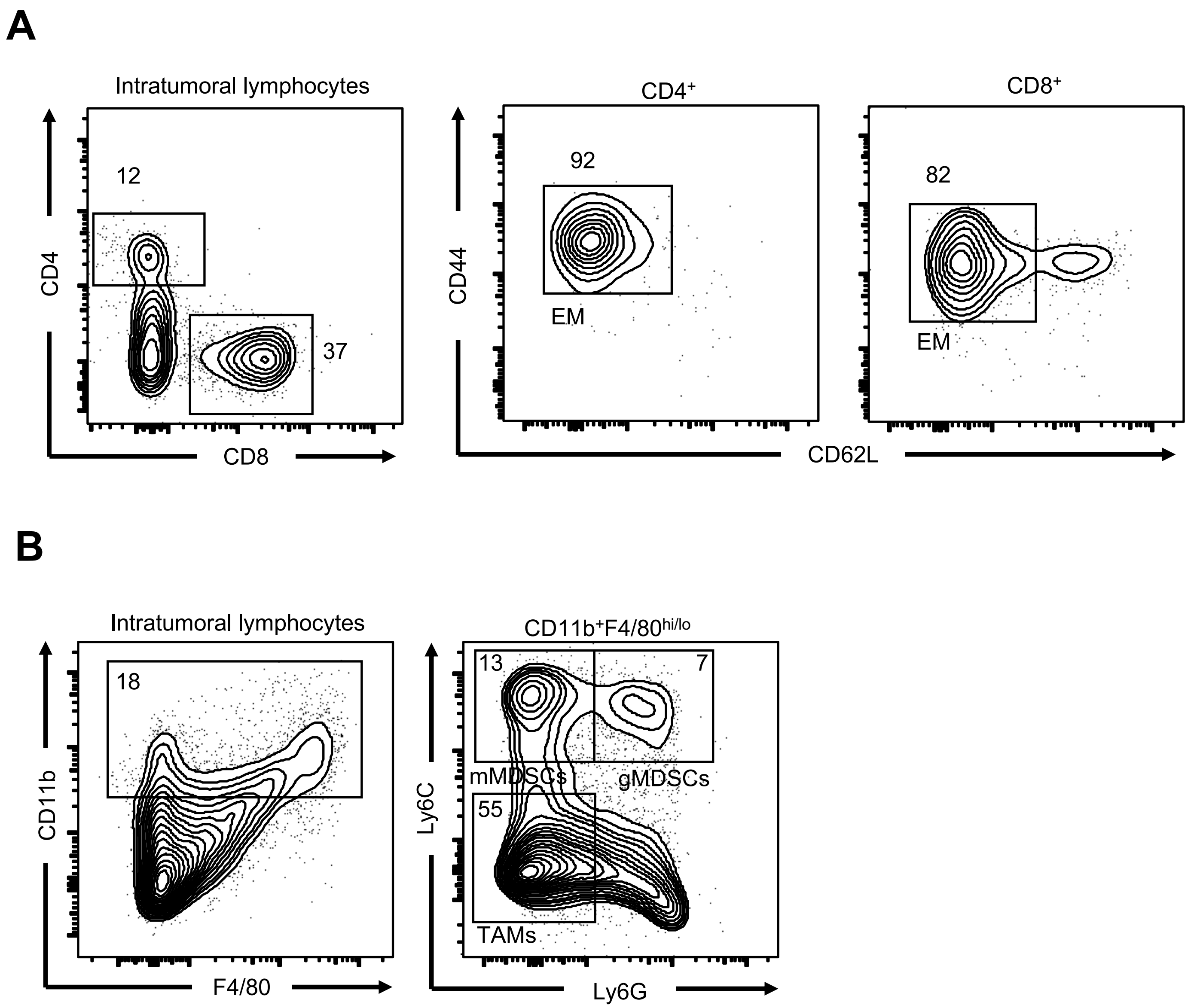
A
Intratumoral lymphocytes
CD4
CD8
12
37
CD4+
CD8+
CD44
CD62L
92
EM
82
EM
B
Intratumoral lymphocytes
CD11b
F4/80
18
CD11b+F4/80hi/lo
Ly6C
Ly6G
13
7
mMDSCs
gMDSCs
55
TAMs

Fig. S1

***Figure S1. Gating strategy for intratumoral lymphocytes*** (related to Fig. 1)**. a-b**) AT3-OVA tumor cells were implanted into the fourth inguinal mammary fat pads of *Ptpn1*$^{+/+}$ mice and tumor-infiltrating lymphocytes (TILs) including **a**) CD4$^{+}$ and CD8$^{+}$ effector/memory (EM; CD44$^{hi}$CD62L$^{lo}$) T cells and **b**) CD11b$^{+}$F4/80$^{hi}$Ly6C$^{–}$Ly6G$^{–}$ tumor-associated macrophages (TAMs), granulocytic CD11b$^{+}$F4/80$^{hi/lo}$Ly6C$^{int}$Ly6G$^{+}$ (gMDSCs) and monocytic CD11b$^{+}$F4/80$^{hi/lo}$ Ly6C$^{+}$Ly6G$^{–}$ (mMDSCs) myeloid-derived suppressor cells and were analysed by flow cytometry.

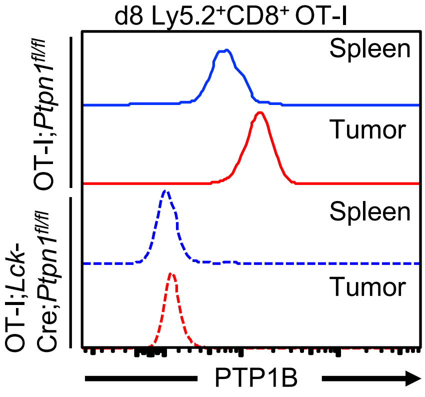

d8 Ly5.2⁺CD8⁺ OT-I
OT-I;Ptpn1fl/fl
OT-I;Lck-Cre;Ptpn1fl/fl
Spleen
Tumor
Spleen
Tumor
PTP1B


**Fig. S2**

***Figure S2. PTP1B protein levels in splenic and intratumoral T cells.*** Naïve ($CD44^{lo}CD62L^{hi}$) $Ly5.2^+CD8^+OT\text{-}I^+$ cells from OT-I;*Ptpn1*$^{fl/fl}$ versus OT-I;*Lck*-Cre;*Ptpn1*$^{fl/fl}$ mice were adoptively transferred into Ly5.1 mice bearing established (40-50 $mm^2$) AT3-OVA mammary tumors. PTP1B protein levels were determined by flow cytometry in splenic and intratumoral donor $Ly5.2^+CD8^+OT\text{-}I^+$ T cells eight days post adoptive transfer; staining in OT-I;*Lck*-Cre;*Ptpn1*$^{fl/fl}$ T cells is background staining. Representative results from at least two independent experiments are shown.

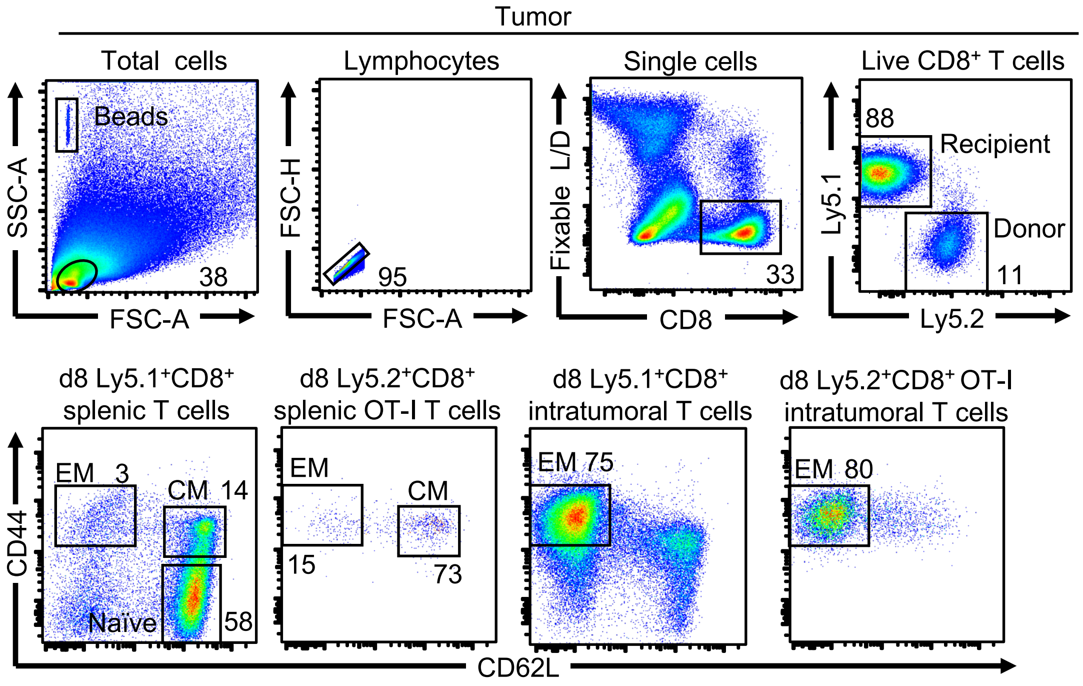
Tumor
Total cells
Beads
38
SSC-A
FSC-A
Lymphocytes
95
FSC-H
FSC-A
Single cells
33
Fixable L/D
CD8
Live CD8+ T cells
88
Recipient
Donor
11
Ly5.1
Ly5.2
d8 Ly5.1+CD8+ splenic T cells
EM 3
CM 14
Naive
58
d8 Ly5.2+CD8+ splenic OT-I T cells
EM
CM
15
73
d8 Ly5.1+CD8+ intratumoral T cells
EM 75
d8 Ly5.2+CD8+ OT-I intratumoral T cells
EM 80
CD44
CD62L

**Fig. S3**

***Figure S3. Gating strategy for adoptively transferred OT-I T cells.*** Naïve ($CD44^{lo}CD62L^{hi}$) $Ly5.2^{+}CD8^{+}OT\text{-}I^{+}$ cells from OT-I;*Ptpn1*$^{fl/fl}$ mice were adoptively transferred into Ly5.1 mice bearing established (40-50 $mm^2$) AT3-OVA mammary tumors. Eight days post adoptive transfer TILs and splenocytes were stained with fluorophore-conjugated antibodies against CD8, Ly5.1, Ly5.2, CD44 and CD62L and analysed by flow cytometry. Fixable live/dead (L/D) was used to exclude dead cells. To discriminate between donor $CD8^{+}$ OT-I T cells and recipient $CD8^{+}$ T cells, live $CD8^{+}$ were gated for $Ly5.1^{-}Ly5.2^{+}$ (donor) and $Ly5.1^{+}Ly5.2^{-}$ (recipient) T cells. Both donor and recipient $CD8^{+}$ T cells were further analysed for naïve ($CD44^{lo}CD62L^{hi}$), central/memory (CM; $CD44^{hi}CD62L^{hi}$) and effector/memory (EM; $CD44^{hi}CD62L^{lo}$) T cells. Representative results from at least two independent experiments are shown.

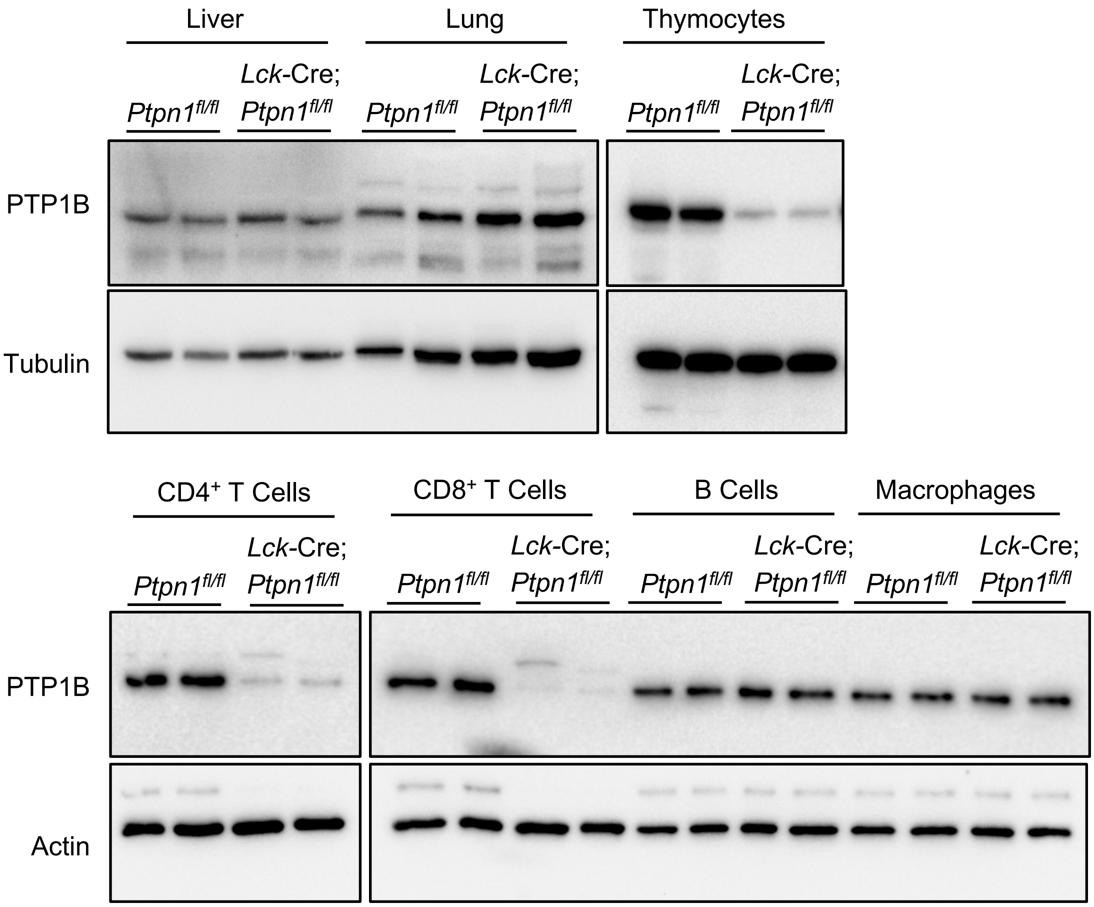
Liver
Lung
Thymocytes
Ptpn1fl/fl
Lck-Cre; Ptpn1fl/fl
PTP1B
Tubulin
CD4+ T Cells
CD8+ T Cells
B Cells
Macrophages
Actin

**Fig. S4**

***Figure S4. PTP1B protein levels in lymphoid and non-lymphoid organs in T cell-specific PTP1B-deficient mice.*** Livers and lungs from 7 week-old *Ptpn1*$^{fl/fl}$ and *Lck*-Cre;*Ptpn1*$^{fl/fl}$ were mechanically homogenised in ice-cold RIPA lysis buffer, clarified by centrifugation (16,000 x g, 10 min, 4$^{o}$C) and proteins resolved by SDS-PAGE and immunoblotted for PTP1B and tubulin. Alternatively, total thymocytes or FACS-purified CD4$^{+}$ and CD8$^{+}$ T cells, CD19$^{+}$ B cells and CD11b$^{+}$F4/80$^{+}$ macrophages from 7 week-old *Ptpn1*$^{fl/fl}$ and *Lck*-Cre;*Ptpn1*$^{fl/fl}$ were lysed in RIPA lysis buffer, clarified by centrifugation (16,000 x g, 10 min, 4$^{o}$C) and proteins resolved by SDS-PAGE and immunoblotted for PTP1B and actin.

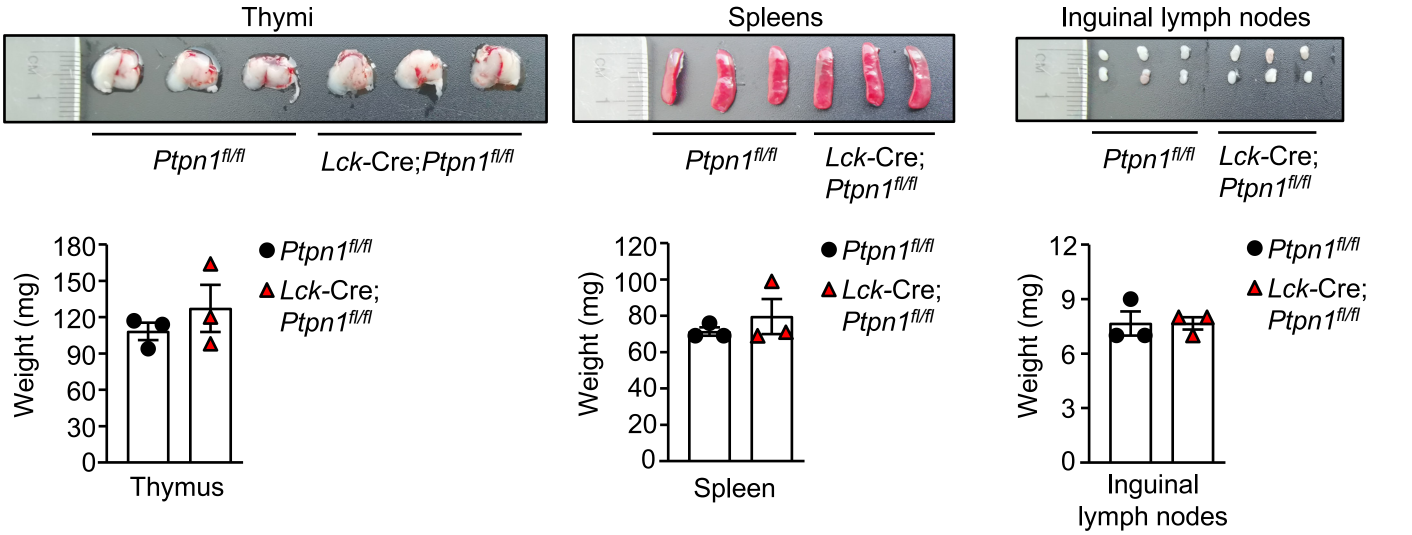

Thymi
Ptpn1^fl/fl
Lck-Cre;Ptpn1^fl/fl
Spleens
Ptpn1^fl/fl
Lck-Cre; Ptpn1^fl/fl
Inguinal lymph nodes
Ptpn1^fl/fl
Lck-Cre; Ptpn1^fl/fl
Weight (mg)
180
150
120
90
60
30
0
Thymus
Ptpn1^fl/fl
Lck-Cre; Ptpn1^fl/fl
120
100
80
60
40
20
0
Spleen
12
9
6
3
Inguinal lymph nodes


**Fig. S5**

***Figure S5. Thymi, spleens and inguinal lymph nodes in T cell-specific PTP1B-deficient mice.*** Thymi, spleens and inguinal lymph nodes from 7 week-old *Ptpn1*$^{fl/fl}$ and *Lck*-Cre;*Ptpn1*$^{fl/fl}$ mice were harvested and organ weights determined. Representative results (means ± SEM) from at least two independent experiments are shown.

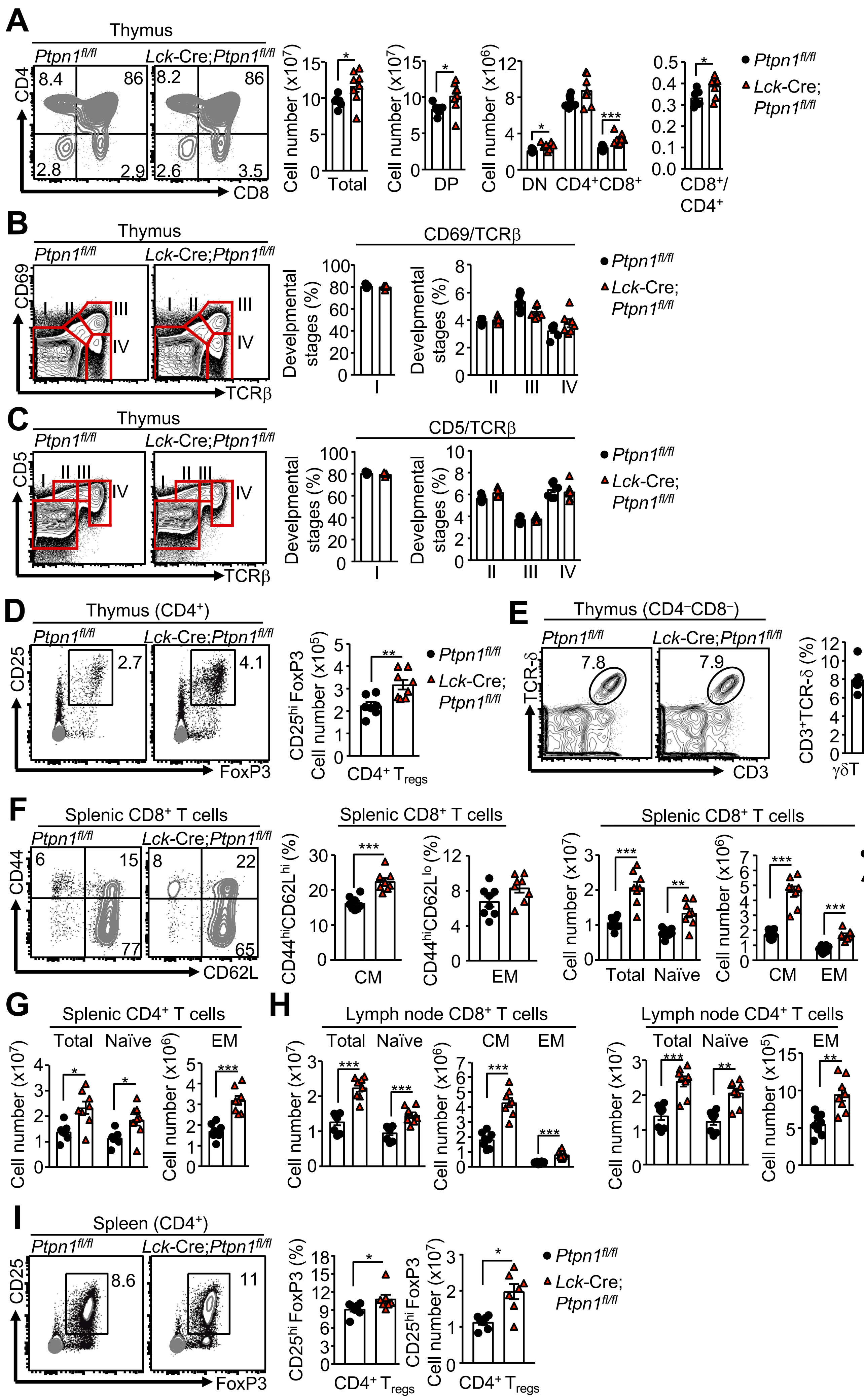

A
Thymus
Ptpn1fl/fl
Lck-Cre;Ptpn1fl/fl
CD4
CD8
Cell number (x10^7)
Total
DP
Cell number (x10^6)
DN
CD4+CD8+
CD8+/CD4+
B
Thymus
CD69
TCRβ
CD69/TCRβ
Develpmental stages (%)
C
Thymus
CD5
TCRβ
CD5/TCRβ
Develpmental stages (%)
D
Thymus (CD4+)
CD25
FoxP3
CD25hi FoxP3 Cell number (x10^5)
CD4+ Tregs
E
Thymus (CD4−CD8−)
TCR-δ
CD3
CD3+TCR-δ (%)
γδT cells
F
Splenic CD8+ T cells
CD44
CD62L
CD44hiCD62Lhi (%)
CD44hiCD62Llo (%)
CM
EM
Total
Naïve
G
Splenic CD4+ T cells
H
Lymph node CD8+ T cells
Lymph node CD4+ T cells
Cell number (x10^7)
Cell number (x10^6)
Cell number (x10^5)
I
Spleen (CD4+)
CD25
FoxP3
CD25hi FoxP3 (%)
CD25hi FoxP3 Cell number (x10^7)
CD4+ Tregs


Fig. S6

***Figure S6. T cell development in T cell-specific PTP1B-deficient mice.*** **a**) Total, $CD4^+CD8^+$ double positive (DP), $CD4^-CD8^-$ double negative (DN), and $CD4^+$ and $CD8^+$ single positive thymocyte numbers in 7 week-old *Ptpn1*$^{fl/fl}$ and *Lck*-Cre;*Ptpn1*$^{fl/fl}$ mice were determined by flow cytometry; $CD8^+$/$CD4^+$ thymocyte ratios were calculated. **b-c**) The four stages of thymocyte development were determined by assessing **b**) CD69 versus TCRβ or **c**) CD5 versus TCRβ expression on total thymocytes. **d**) The number of regulatory $CD4^+CD25^{hi}FoxP3^+$ T cells ($T_{regs}$) in thymi were determined by flow cytometry. **e**) The proportion of $CD4^-CD8^-CD3^+TCR\delta^+$ γδT cells in thymi were determined. **f**) The proportions and numbers of splenic $CD8^+$ central memory (CM; $CD44^{hi}CD62L^{hi}$) and effector memory (EM; $CD44^{hi}CD62L^{lo}$) T cells were determined. **g**) The number of total, naïve and EM $CD4^+$ T cells were determined in spleens. **h**) The number of total, naïve, CM and EM $CD8^+$ and $CD4^+$ T cells were determined in lymph nodes. **i**) The number of $T_{regs}$ in the spleen were determined by flow cytometry. Representative results (means ± SEM) from at least two independent experiments are shown. In (a, d-i) significance was determined using 2-tailed Mann-Whitney U Test.

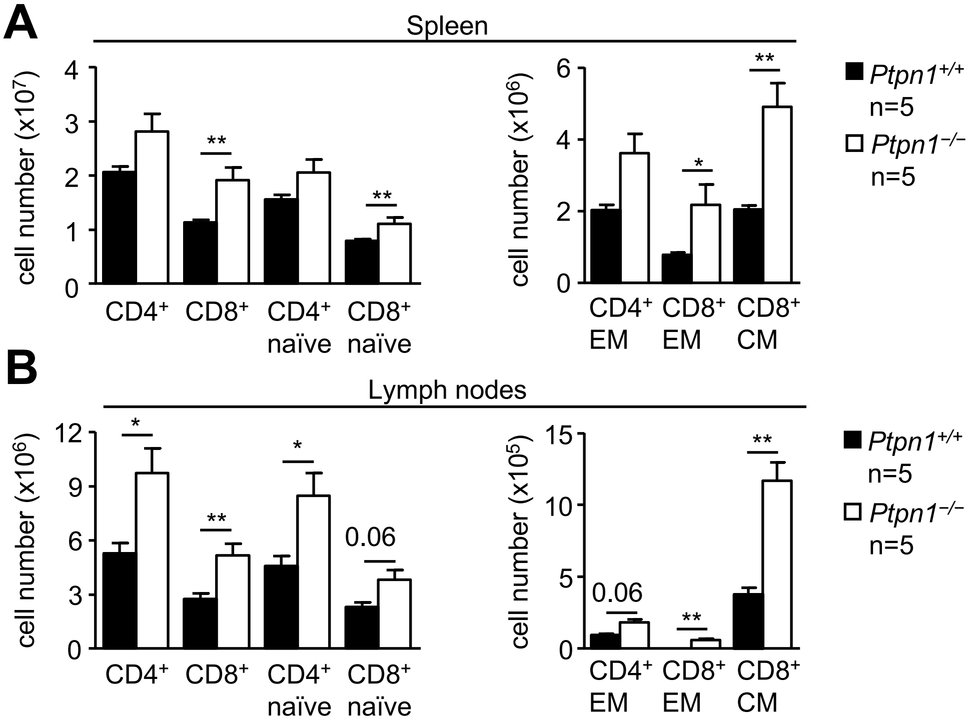
A
Spleen
cell number (x10^7)
4
3
2
1
0
**
**
CD4+
CD8+
CD4+ naïve
CD8+ naïve
cell number (x10^6)
6
4
2
0
*
**
CD4+ EM
CD8+ EM
CD8+ CM
Ptpn1+/+ n=5
Ptpn1−/− n=5
B
Lymph nodes
cell number (x10^6)
12
9
6
3
0
*
**
*
0.06
CD4+
CD8+
CD4+ naïve
CD8+ naïve
cell number (x10^5)
15
10
5
0
0.06
**
**
CD4+ EM
CD8+ EM
CD8+ CM
Ptpn1+/+ n=5
Ptpn1−/− n=5

**Fig. S7**

***Figure S7. T cell development in global PTP1B knockout mice.*** **a-b**) Total, naïve ($CD44^{lo}CD62L^{hi}$), central memory (CM; $CD44^{hi}CD62L^{hi}$) and effector memory (EM; $CD44^{hi}CD62L^{lo}$) $CD4^{+}$ and $CD8^{+}$ T cell numbers were determined in the **a**) spleens and **b**) lymph nodes of *Ptpn1*$^{+/+}$ and *Ptpn1*$^{-/-}$ mice. Representative results (means ± SEM) from two independent experiments (each with n ≥ 5) are shown. Significance was determined using 2-tailed Mann-Whitney U Test.

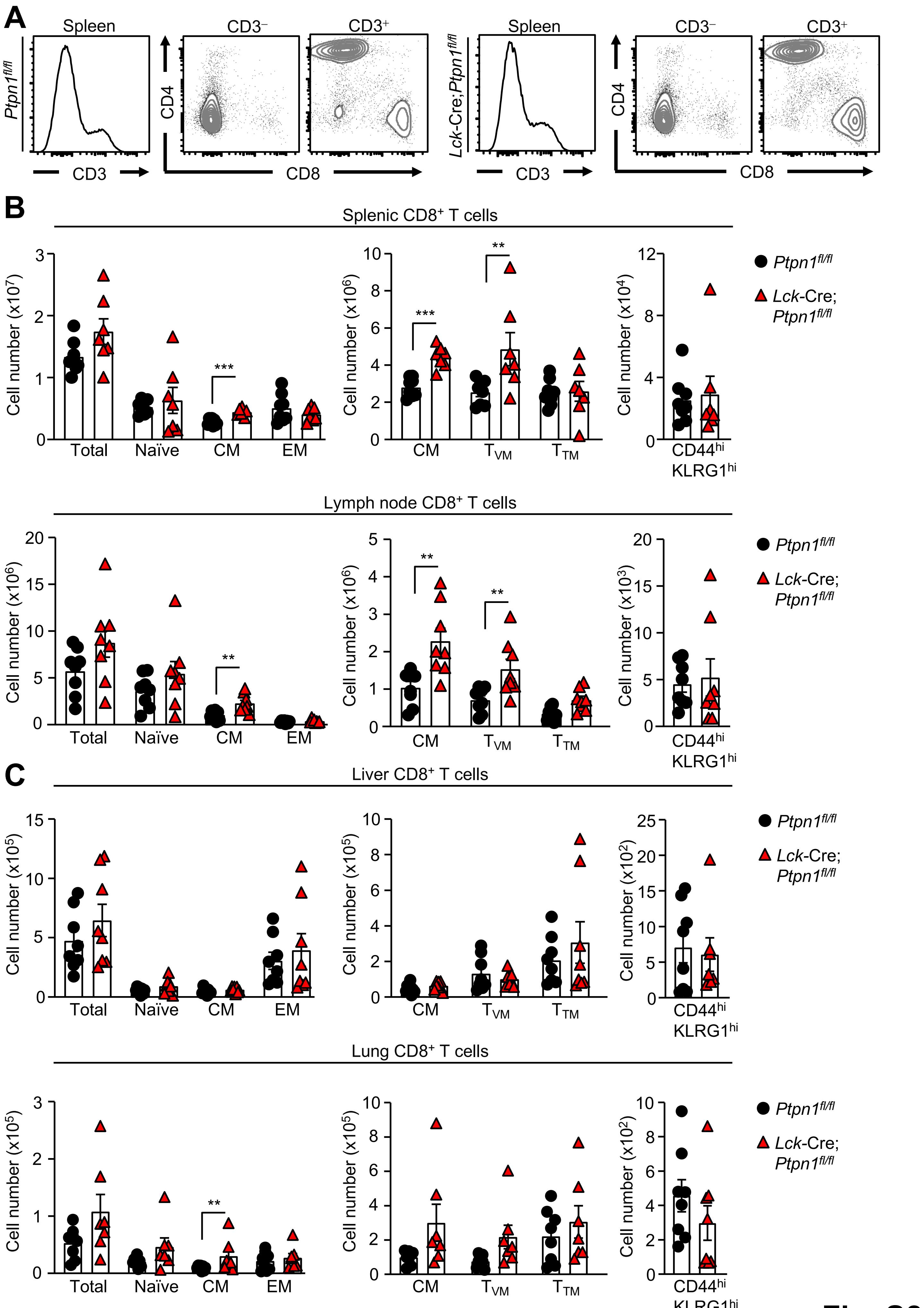
A
Spleen
CD3⁻
CD3⁺
Ptpn1fl/fl
CD3
CD4
CD8
Lck-Cre;Ptpn1fl/fl
B
Splenic CD8⁺ T cells
Cell number (x10⁷)
Total
Naïve
CM
EM
Cell number (x10⁶)
TVM
TTM
Cell number (x10⁴)
CD44hi KLRG1hi
Ptpn1fl/fl
Lck-Cre; Ptpn1fl/fl
Lymph node CD8⁺ T cells
Cell number (x10³)
C
Liver CD8⁺ T cells
Cell number (x10⁵)
Cell number (x10²)
Lung CD8⁺ T cells
***
**

Fig. S8

***Figure S8. Peripheral T cell development in one-year old T cell-specific PTP1B-deficient mice.*** **a**) Representative $CD4^+$ versus $CD8^+$ contour plots gated on $CD3^-$ and $CD3^+$ splenocytes from one-year old *Ptpn1$^{fl/fl}$* and *Lck*-Cre;*Ptpn1$^{fl/fl}$* mice. **b-c**) $CD8^+$ total, naïve ($CD44^{lo}CD62L^{hi}$), central/memory (CM; $CD44^{hi}CD62L^{hi}$), effector/memory (EM; $CD44^{hi}CD62L^{lo}$), virtual memory (VM; $CD44^{hi}CD49d^{lo}$; VM have a memory phenotype and arise from the engagement of self-antigen rather than foreign antigen), antigen-experienced true memory (TM; $CD44^{hi}CD49d^{hi}$; arise from engagement of foreign antigen) and $CD44^{hi}KLGR1^{hi}$ terminally-differentiated effector T cell numbers were determined in (b) spleens and lymph nodes and (c) livers and lungs from one-year old *Ptpn1$^{fl/fl}$* and *Lck*-Cre;*Ptpn1$^{fl/fl}$* mice. Representative results (means ± SEM) from two independent experiments are shown. Significance was determined using 2-tailed Mann-Whitney U Test.

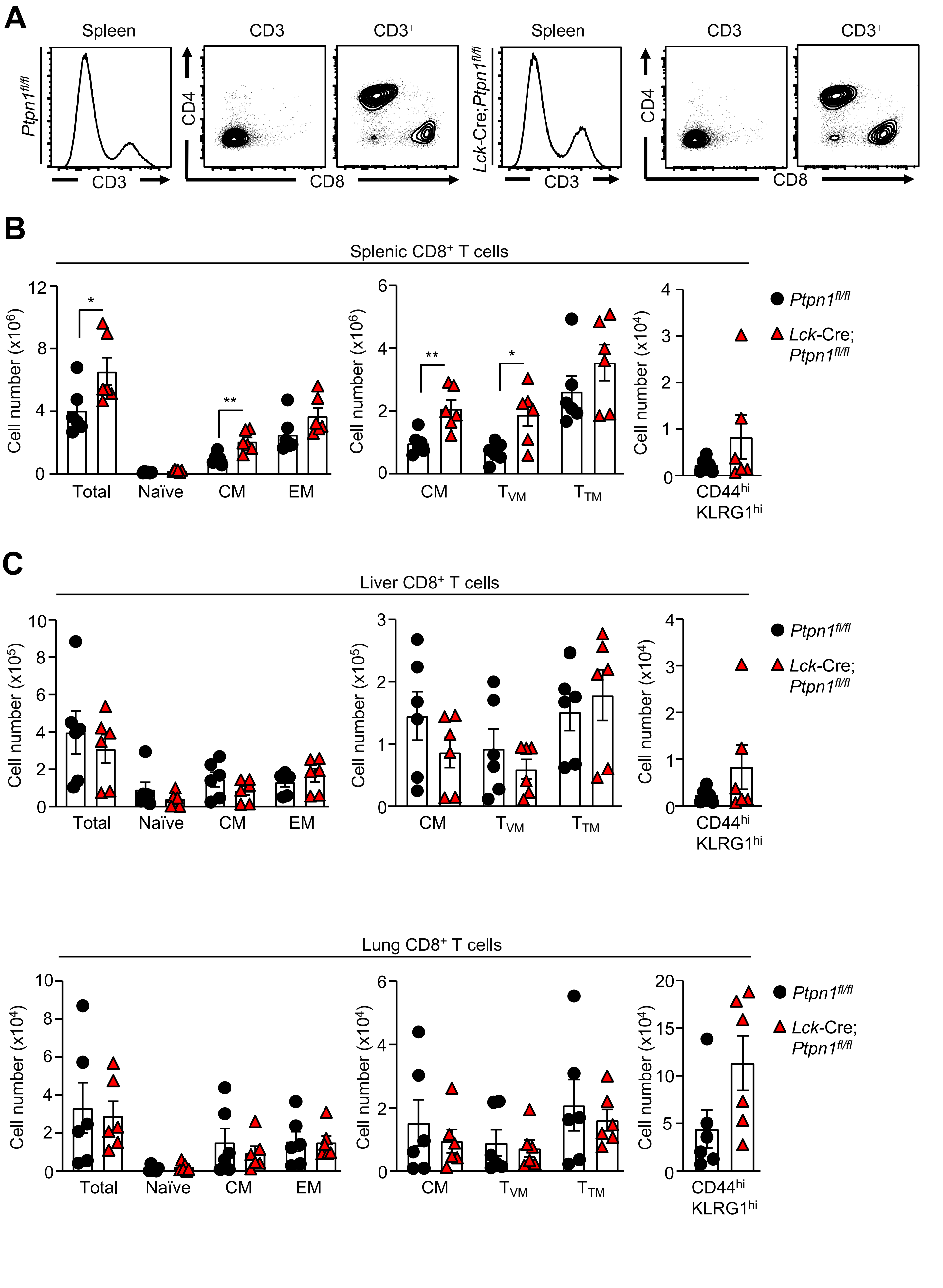
A
Spleen
CD3⁻
CD3⁺
Ptpn1fl/fl
CD3
CD4
CD8
Lck-Cre;Ptpn1fl/fl
B
Splenic CD8⁺ T cells
Cell number (x10⁶)
Cell number (x10⁴)
Total
Naïve
CM
EM
TVM
TTM
CD44hi KLRG1hi
Ptpn1fl/fl
Lck-Cre; Ptpn1fl/fl
C
Liver CD8⁺ T cells
Cell number (x10⁵)
Lung CD8⁺ T cells


Fig. S9

***Figure S9. Peripheral T cell development in two-year-old T cell-specific PTP1B-deficient mice.*** **a**) Representative $CD4^+$ versus $CD8^+$ contour plots gated on $CD3^-$ and $CD3^+$ splenocytes from two-year old *Ptpn1*$^{fl/fl}$ and *Lck*-Cre;*Ptpn1*$^{fl/fl}$ mice. **b-c**) $CD8^+$ total, naïve ($CD44^{lo}CD62L^{hi}$), central/memory (CM; $CD44^{hi}CD62L^{hi}$), effector/memory (EM; $CD44^{hi}CD62L^{lo}$), virtual memory (VM; $CD44^{hi}CD49d^{lo}$), antigen-experienced true memory (TM; $CD44^{hi}CD49d^{hi}$) and $CD44^{hi}KLGR1^{hi}$ terminally-differentiated effector T cell numbers were determined in the (b) spleens and (c) livers and lungs from two-year old *Ptpn1*$^{fl/fl}$ and *Lck*-Cre;*Ptpn1*$^{fl/fl}$ mice. Results shown are means ± SEM for the indicated number of mice. Significance was determined using 2-tailed Mann-Whitney U Test.

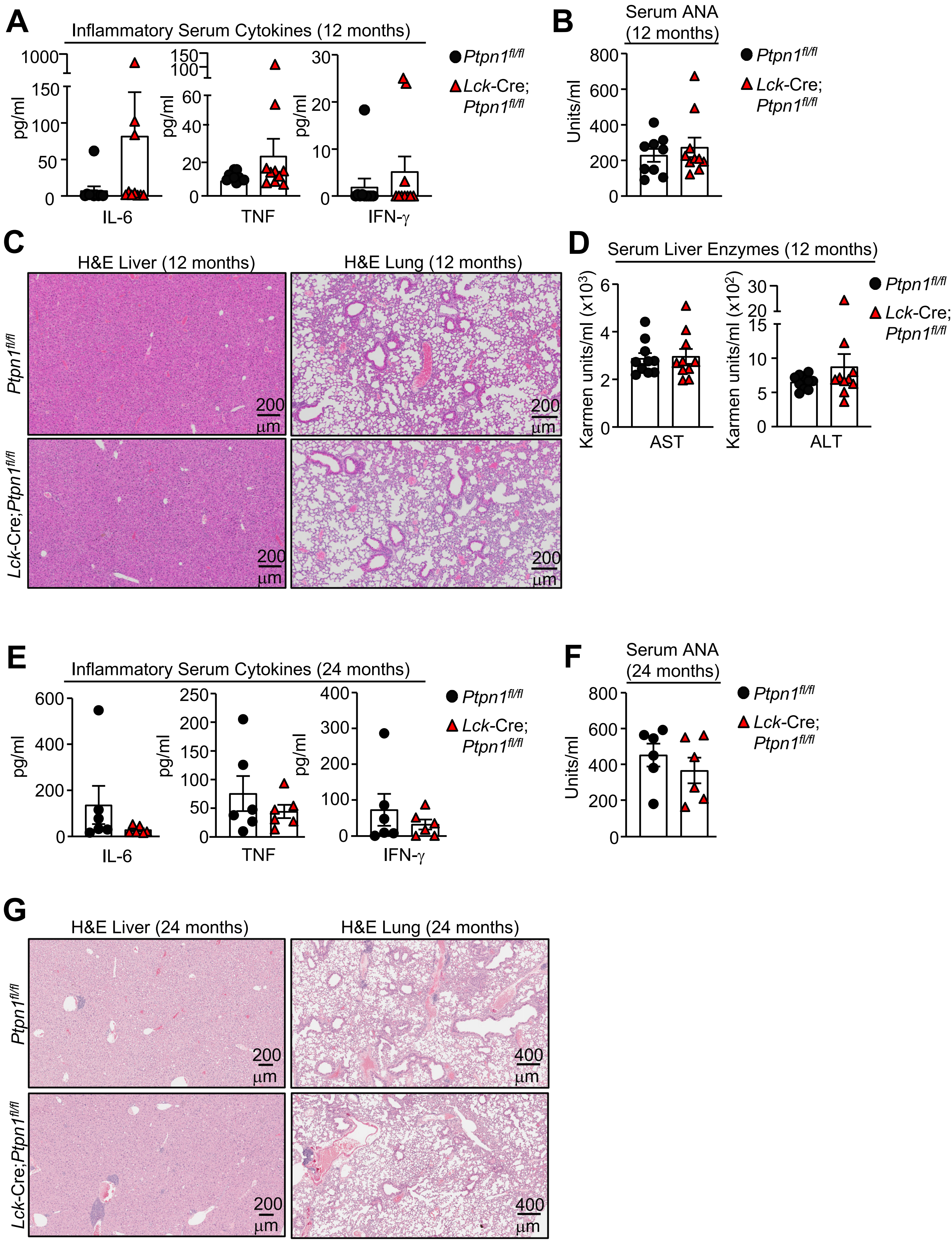

A
Inflammatory Serum Cytokines (12 months)
pg/ml
IL-6
TNF
IFN-γ
Ptpn1fl/fl
Lck-Cre; Ptpn1fl/fl
B
Serum ANA (12 months)
Units/ml
C
H&E Liver (12 months)
H&E Lung (12 months)
200 µm
D
Serum Liver Enzymes (12 months)
Karmen units/ml (x10³)
Karmen units/ml (x10²)
AST
ALT
E
Inflammatory Serum Cytokines (24 months)
F
Serum ANA (24 months)
G
H&E Liver (24 months)
H&E Lung (24 months)
400 µm


Fig. S10

***Figure S10. Inflammation and autoimmunity and are not evident in aged T cell-specific PTP1B-deficient mice.*** Serum cytokines in **a**) one-year old and **e**) two-year old *Ptpn1*$^{fl/fl}$ and *Lck*-Cre;*Ptpn1*$^{fl/fl}$ mice were determined by flow cytometry using a BD Cytokine Bead Array (BD Biosciences). Serum anti-nuclear antibodies (ANA) in **b**) one-year old and **f**) two-year old *Ptpn1*$^{fl/fl}$ and *Lck*-Cre;*Ptpn1*$^{fl/fl}$ mice. Livers and lungs from **c**) one-year old and **g**) two-year old *Ptpn1*$^{fl/fl}$ and *Lck*-Cre;*Ptpn1*$^{fl/fl}$ mice were fixed in formalin and processed for histological assessment (hematoxylin and eosin: H&E). **d**) Serum liver enzymes AST and ALT in one-year old *Ptpn1*$^{fl/fl}$ and *Lck*-Cre;*Ptpn1*$^{fl/fl}$ mice. Results shown are means ± SEM for the indicated number of mice.

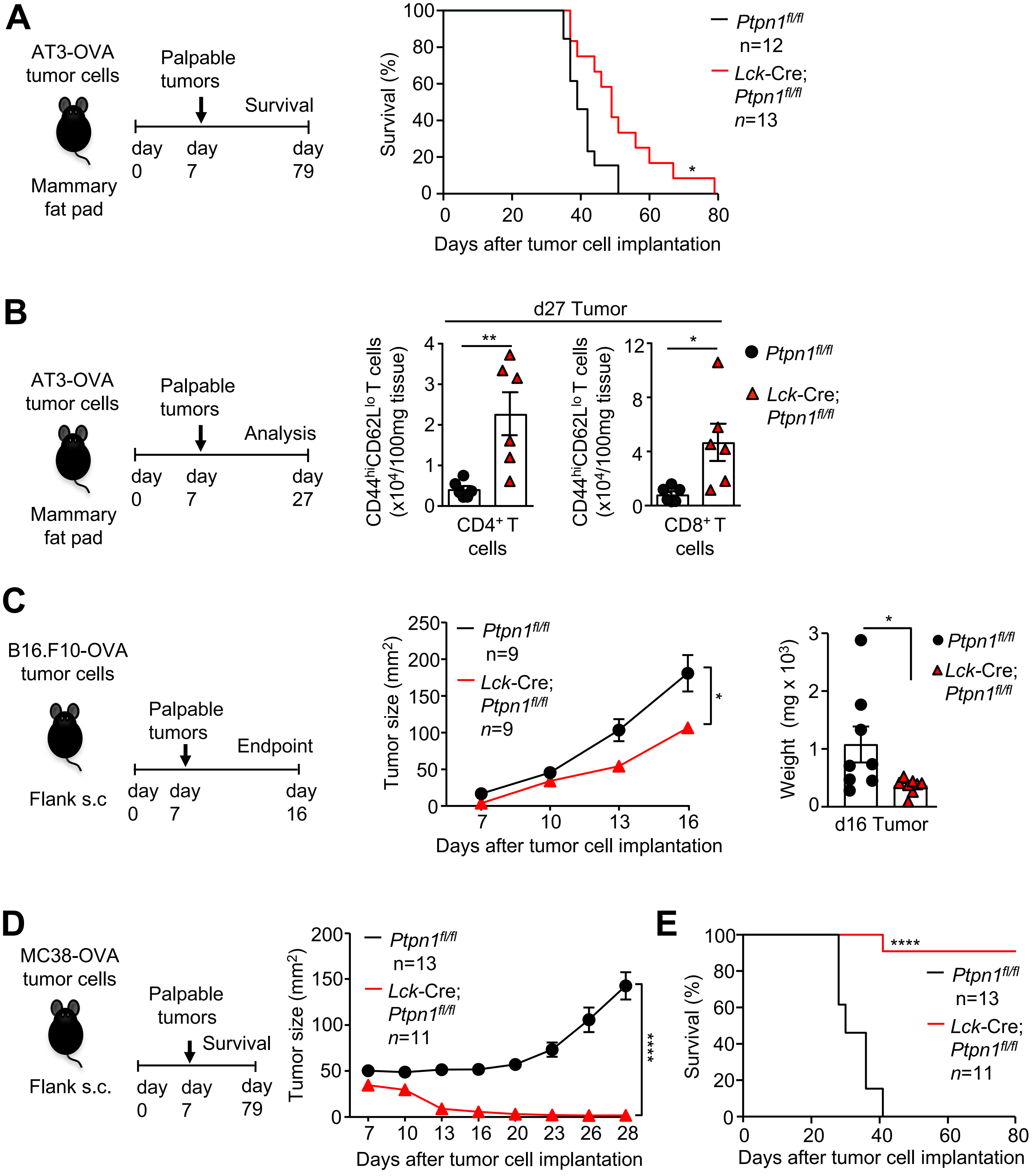
A
AT3-OVA tumor cells
Mammary fat pad
Palpable tumors
Survival
day 0
day 7
day 79
Survival (%)
Days after tumor cell implantation
Ptpn1fl/fl n=12
Lck-Cre; Ptpn1fl/fl n=13
*
B
AT3-OVA tumor cells
Mammary fat pad
Palpable tumors
Analysis
day 0
day 7
day 27
d27 Tumor
CD44hiCD62Llo T cells (x10⁴/100mg tissue)
CD4⁺ T cells
CD8⁺ T cells
**
*
Ptpn1fl/fl
Lck-Cre; Ptpn1fl/fl
C
B16.F10-OVA tumor cells
Flank s.c
Palpable tumors
Endpoint
day 0
day 7
day 16
Tumor size (mm²)
Days after tumor cell implantation
Ptpn1fl/fl n=9
Lck-Cre; Ptpn1fl/fl n=9
*
Weight (mg x 10³)
d16 Tumor
Ptpn1fl/fl
Lck-Cre; Ptpn1fl/fl
D
MC38-OVA tumor cells
Flank s.c.
Palpable tumors
Survival
day 0
day 7
day 79
Tumor size (mm²)
Days after tumor cell implantation
Ptpn1fl/fl n=13
Lck-Cre; Ptpn1fl/fl n=11
****
E
Survival (%)
Days after tumor cell implantation
Ptpn1fl/fl n=13
Lck-Cre; Ptpn1fl/fl n=11
****

**Fig. S11**

***Figure S11. PTP1B-deficiency enhances CD8$^{+}$ T cell-mediated anti-tumor immunity.*** **a**) AT3-OVA tumor cells were implanted into the fourth inguinal mammary fat pads of *Ptpn1*$^{fl/fl}$ or *Lck*-Cre;*Ptpn1*$^{fl/fl}$ mice and survival was monitored. **b**) Related to Fig. 3a: AT3-OVA tumor cells were implanted into the fourth inguinal mammary fat pads of *Ptpn1*$^{fl/fl}$ or *Lck*-Cre;*Ptpn1*$^{fl/fl}$ mice and tumor-infiltrating CD4$^{+}$ and CD8$^{+}$ T cells analysed by flow cytometry. **c**) B16F10-OVA tumor cells were xenografted into the flanks of C57BL/6 mice and tumor growth and final tumor weights were determined. **d-e**) MC38-OVA tumor cells were xenografted into the flanks of *Ptpn1*$^{fl/fl}$ or *Lck*-Cre;*Ptpn1*$^{fl/fl}$ mice and **d**) tumor growth and **e**) survival monitored. Representative results (means ± SEM) from at least two independent experiments are shown. For tumor growth curves in (c-d) significance was determined using 2-way ANOVA Test. Significances for T cell infiltrates and tumor weights in (b-c) were determined using 2-tailed Mann-Whitney U Test. In (a, d) significances were determined using Log-rank (Mantel-Cox) test.

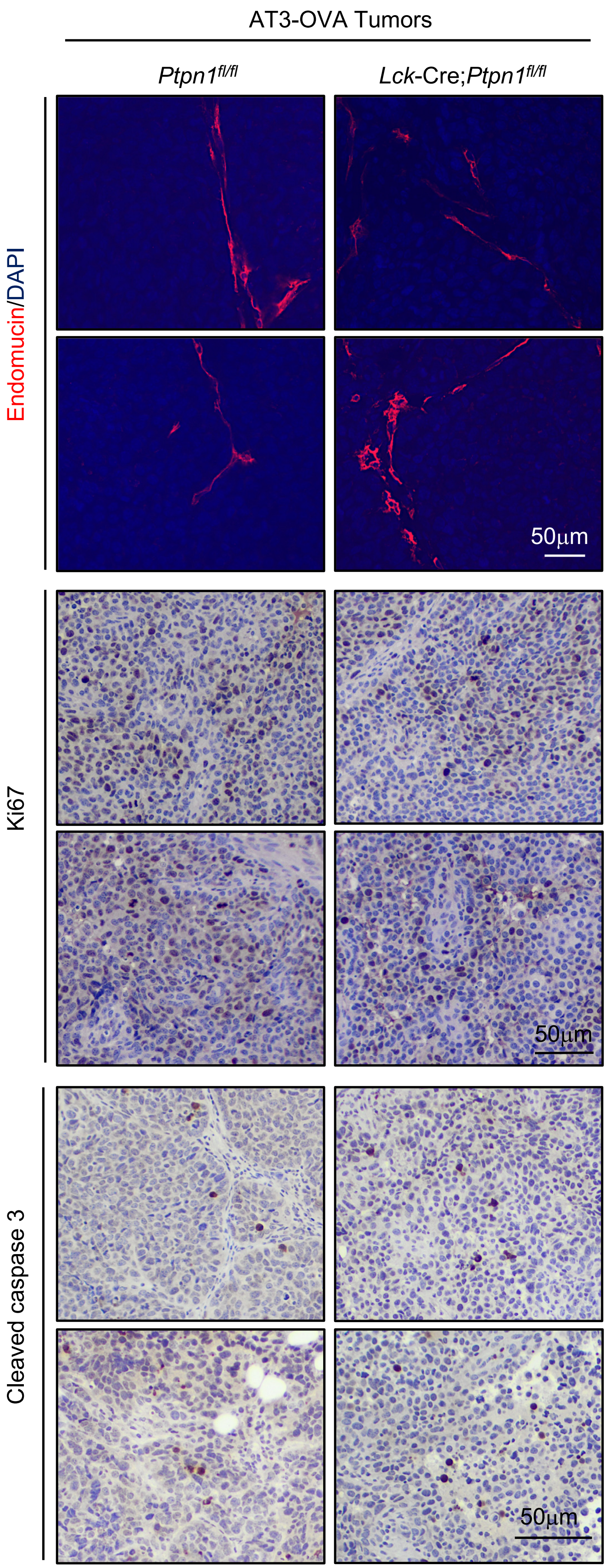
AT3-OVA Tumors
Ptpn1fl/fl
Lck-Cre;Ptpn1fl/fl
Endomucin/DAPI
Ki67
Cleaved caspase 3
50µm
50µm
50µm

Fig. S12

***Figure S12. T cell-specific PTP1B-deficiency does not affect tumor cell proliferation, apoptosis or angiogenesis.*** AT3-OVA tumor cells were implanted into the fourth inguinal mammary fat pads of *Ptpn1*$^{fl/fl}$ or *Lck*-Cre;*Ptpn1*$^{fl/fl}$ mice. Tumors were extracted on day 19 and processed for immunohistochemistry to monitor for angiogenesis (endomucin) and tumor cell proliferation (Ki67) and apoptosis (cleave caspase 3). Representative results are shown.

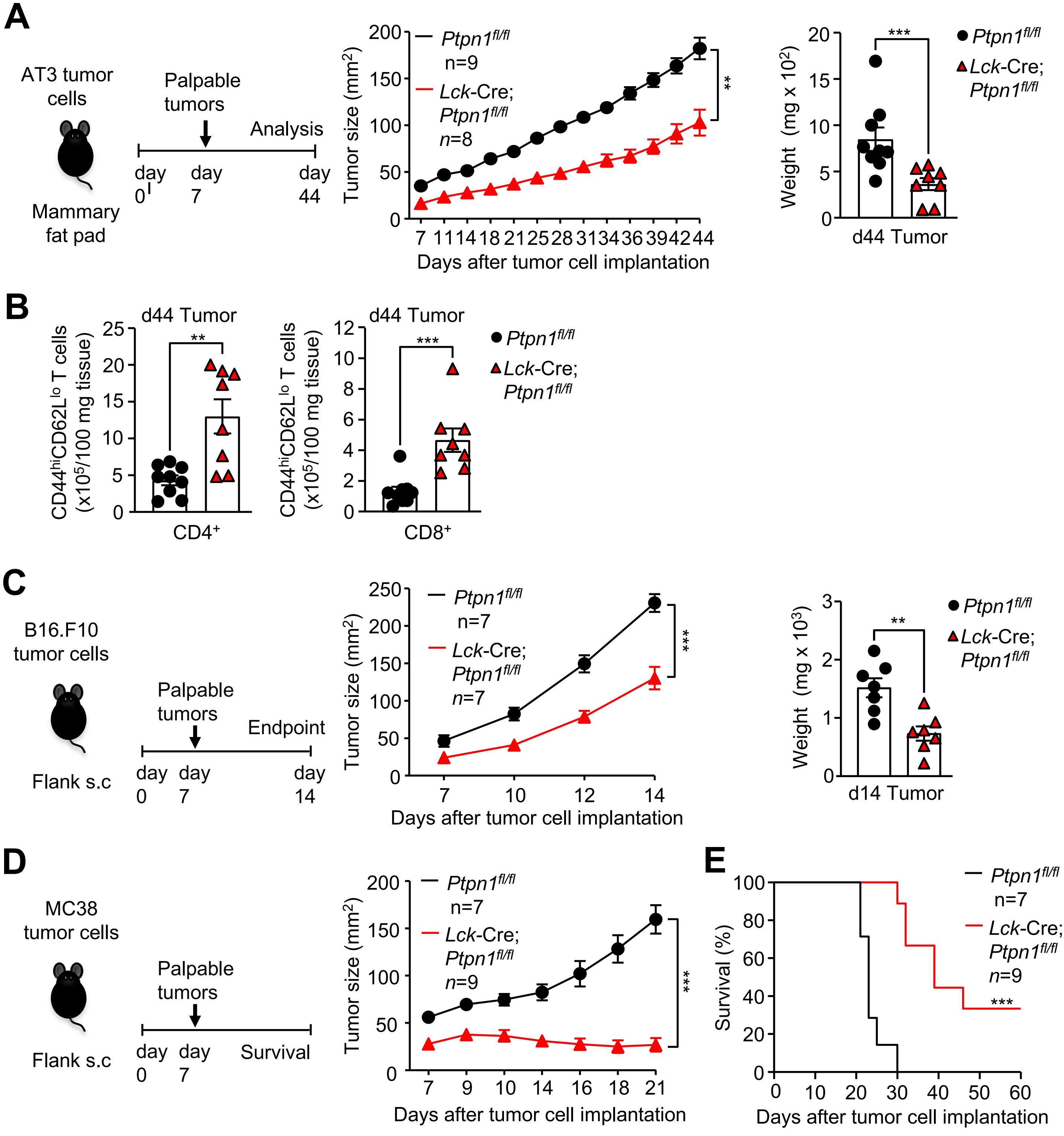
A
AT3 tumor cells
Mammary fat pad
Palpable tumors
Analysis
day 0
day 7
day 44
Tumor size (mm2)
Ptpn1fl/fl n=9
Lck-Cre; Ptpn1fl/fl n=8
Days after tumor cell implantation
Weight (mg x 102)
d44 Tumor
B
d44 Tumor
CD44hiCD62Llo T cells (x105/100 mg tissue)
CD4+
CD8+
C
B16.F10 tumor cells
Flank s.c
Palpable tumors
Endpoint
day 0
day 7
day 14
Ptpn1fl/fl n=7
Lck-Cre; Ptpn1fl/fl n=7
Weight (mg x 103)
d14 Tumor
D
MC38 tumor cells
Flank s.c
Survival
Lck-Cre; Ptpn1fl/fl n=9
E
Survival (%)

Fig. S13

***Figure S13. PTP1B-deficient CD8$^+$ T cells repress the growth of OVA negative tumors.*** **a**) AT3 tumor cells were implanted into the fourth inguinal mammary fat pads of *Ptpn1*$^{fl/fl}$ or *Lck*-Cre;*Ptpn1*$^{fl/fl}$ mice and tumor growth was monitored and final tumor weights determined. **b**) Tumor-infiltrating CD4$^+$ and CD8$^+$ T cells from (a) were analysed by flow cytometry. **c**) B16F10 tumor cells were xenografted into the flanks of C57BL/6 mice and tumor growth and final tumor weights determined. **d-e**) MC38 tumor cells were xenografted into the flanks of *Ptpn1*$^{fl/fl}$ or *Lck*-Cre;*Ptpn1*$^{fl/fl}$ mice and **d**) tumor growth and **e**) survival monitoredRepresentative results (means ± SEM) from at least two independent experiments (a-e) and one experiment (f) are shown. For tumor growth curves in (a, c, d) significance was determined using 2-way ANOVA Test. Significances for T cell infiltrates and tumor weights in (a-c) was determined using 2-tailed Mann-Whitney U Test. In (e) significance was determined using Log-rank (Mantel-Cox) test.

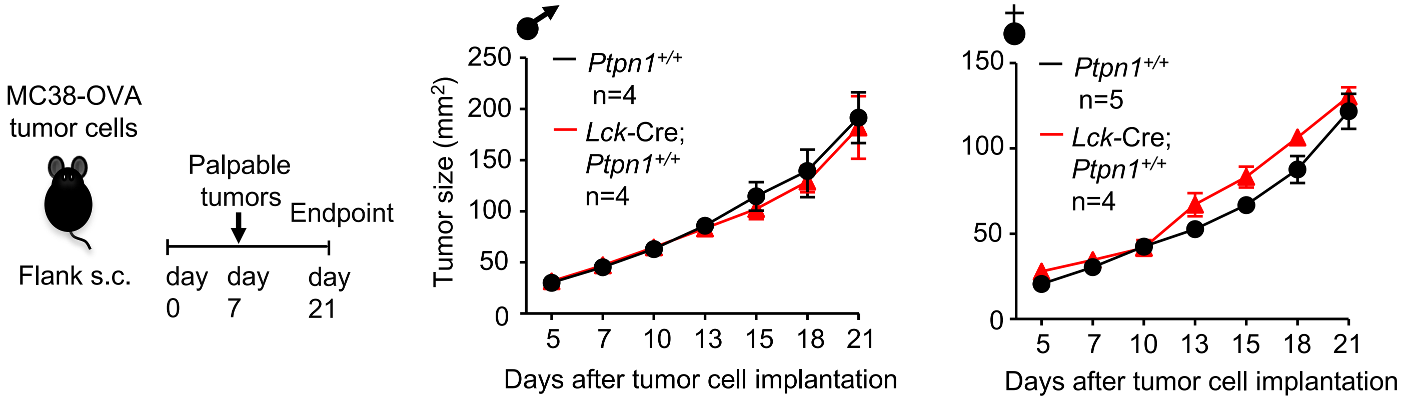
MC38-OVA
tumor cells
Flank s.c.
Palpable
tumors
Endpoint
day 0
day 7
day 21
Tumor size (mm²)
Ptpn1+/+
n=4
Lck-Cre;
Ptpn1+/+
n=4
Ptpn1+/+
n=5
Lck-Cre;
Ptpn1+/+
n=4
Days after tumor cell implantation
Days after tumor cell implantation

**Fig. S14**

***Figure S14. The Lck-Cre transgene does not affect the growth of syngeneic tumors.*** MC38-OVA tumor cells were xenografted into the flanks of male or female *Ptpn1*$^{+/+}$ and *Lck*-Cre mice and tumor growth monitored. Results shown are means ± SEM for the indicated number of mice.

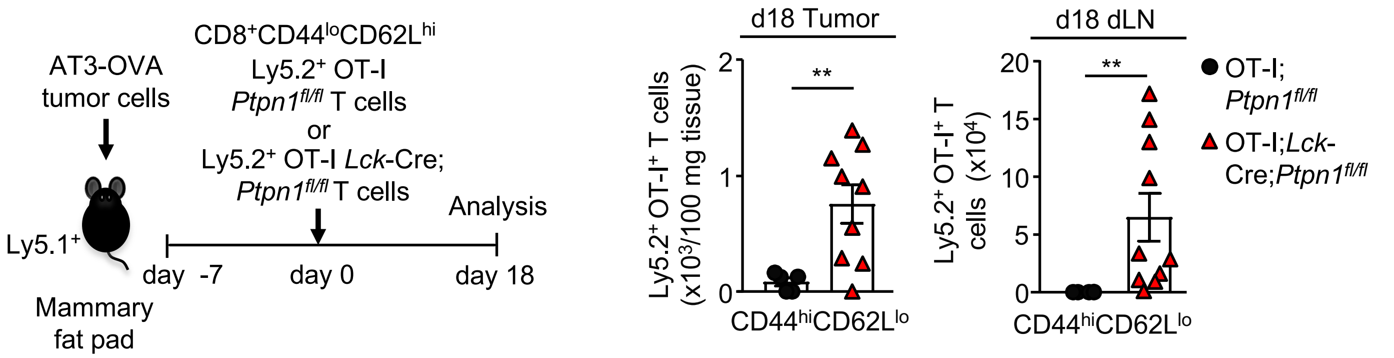
AT3-OVA
tumor cells
CD8+CD44loCD62Lhi
Ly5.2+ OT-I
Ptpn1fl/fl T cells
or
Ly5.2+ OT-I Lck-Cre;
Ptpn1fl/fl T cells
Analysis
Ly5.1+
day -7
day 0
day 18
Mammary
fat pad
d18 Tumor
Ly5.2+ OT-I+ T cells
(x10³/100 mg tissue)
CD44hiCD62Llo
d18 dLN
Ly5.2+ OT-I+ T
cells (x10⁴)
CD44hiCD62Llo
**
OT-I;
Ptpn1fl/fl
OT-I;Lck-
Cre;Ptpn1fl/fl

Fig. S15

***Figure S15. PTP1B-deficiency increases the abundance of intratumoral OT-I CD8$^{+}$ T cells*** (related to Fig. 3f)**.** Naïve (CD44$^{lo}$CD62L$^{hi}$) Ly5.2$^{+}$CD8$^{+}$OT-I$^{+}$ T cells from OT-I;*Ptpn1$^{fl/fl}$* or OT-I;*Lck*-Cre;*Ptpn1$^{fl/fl}$* mice were adoptively transferred into Ly5.1 mice bearing established (40-50 mm$^{2}$) AT3-OVA mammary tumors and CD44$^{hi}$CD62L$^{lo}$ effector/memory Ly5.2$^{+}$CD8$^{+}$OT-I$^{+}$ T cells monitored in the tumors and draining lymph nodes (dLN). Representative results (means ± SEM) from at least two independent experiments are shown. Significance was determined using 2-tailed Mann-Whitney U Test.

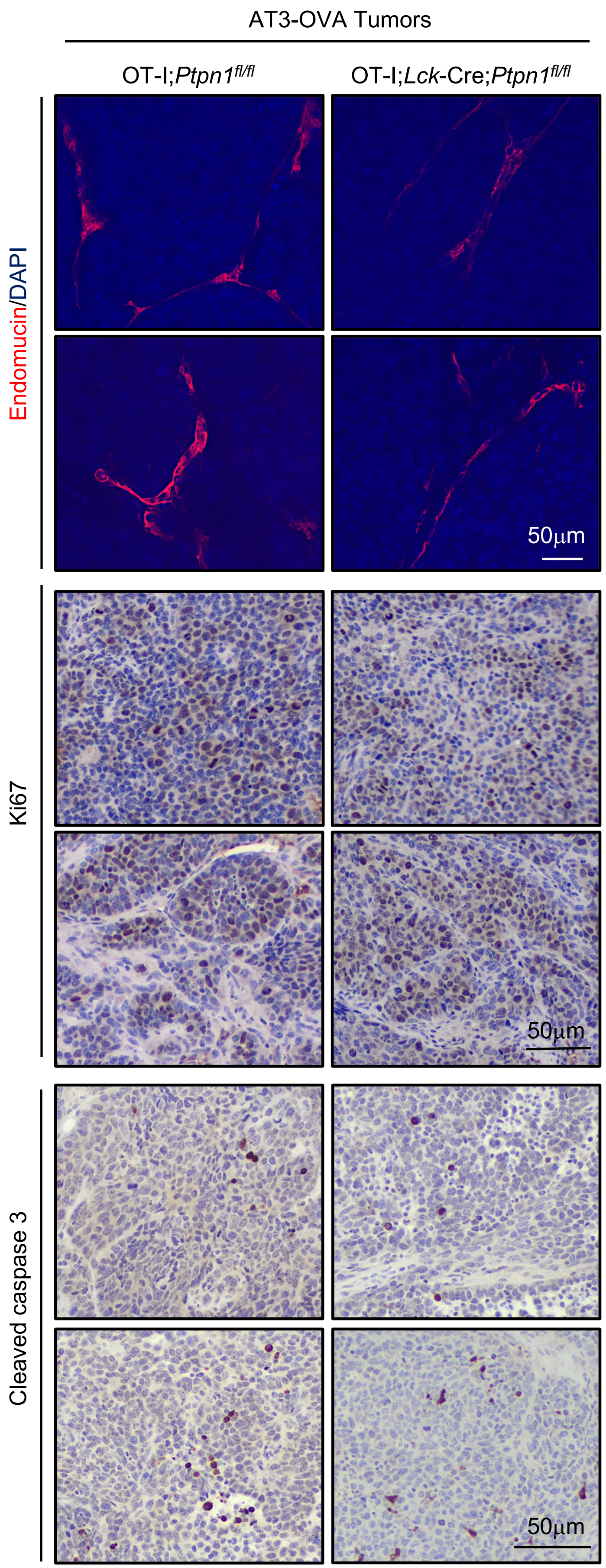
AT3-OVA Tumors
OT-I;Ptpn1fl/fl
OT-I;Lck-Cre;Ptpn1fl/fl
Endomucin/DAPI
Ki67
Cleaved caspase 3
50µm
50µm
50µm

Fig. S16

***Figure S16. PTP1B-deficiency in adoptively transferred OT-I CD8⁺ T cells does not alter tumor cell proliferation, apoptosis or angiogenesis.*** Naïve ($CD44^{lo}CD62L^{hi}$) $Ly5.2^{+}CD8^{+}OT$-$I^{+}$ T cells from OT-I;*Ptpn1*$^{fl/fl}$ or OT-I;*Lck*-Cre;*Ptpn1*$^{fl/fl}$ mice were adoptively transferred into Ly5.1 mice bearing established (40-50 $mm^2$) AT3-OVA mammary tumors. Tumors were extracted on day 16 post adoptive transfer and then processed for immunohistochemistry to monitor for angiogenesis (endomucin) and tumor cell proliferation (Ki67) and apoptosis (cleave caspase 3). Representative results are shown.

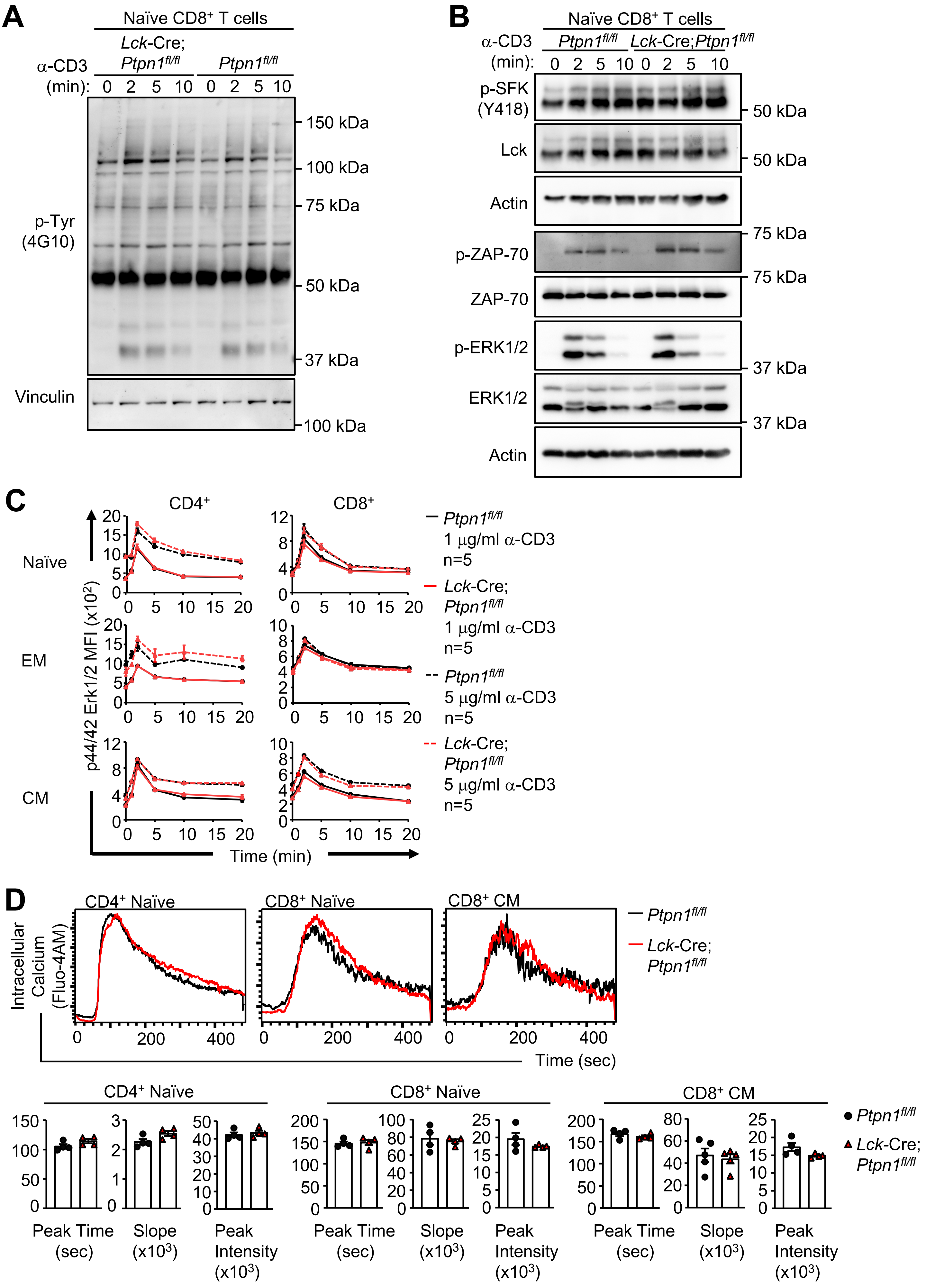

A
Naïve CD8⁺ T cells
Lck-Cre; Ptpn1fl/fl
Ptpn1fl/fl
α-CD3 (min): 0 2 5 10 0 2 5 10
p-Tyr (4G10)
150 kDa
100 kDa
75 kDa
50 kDa
37 kDa
Vinculin
100 kDa
B
Naïve CD8⁺ T cells
α-CD3 (min):
Ptpn1fl/fl
Lck-Cre;Ptpn1fl/fl
0 2 5 10 0 2 5 10
p-SFK (Y418)
Lck
Actin
p-ZAP-70
ZAP-70
p-ERK1/2
ERK1/2
Actin
50 kDa
50 kDa
75 kDa
75 kDa
37 kDa
37 kDa
C
CD4⁺
CD8⁺
Naïve
EM
CM
p44/42 Erk1/2 MFI (x10²)
Time (min)
Ptpn1fl/fl 1 µg/ml α-CD3 n=5
Lck-Cre; Ptpn1fl/fl 1 µg/ml α-CD3 n=5
Ptpn1fl/fl 5 µg/ml α-CD3 n=5
Lck-Cre; Ptpn1fl/fl 5 µg/ml α-CD3 n=5
D
CD4⁺ Naïve
CD8⁺ Naïve
CD8⁺ CM
Intracellular Calcium (Fluo-4AM)
Time (sec)
Ptpn1fl/fl
Lck-Cre; Ptpn1fl/fl
CD4⁺ Naïve
CD8⁺ Naïve
CD8⁺ CM
Peak Time (sec)
Slope (x10³)
Peak Intensity (x10³)


Fig. S17

***Figure S17. T cell receptor signaling is not altered in T cell-specific PTP1B-deficient mice.*** **a-b**) Naïve $CD8^+$ T cells from *Ptpn1*$^{fl/fl}$ and *Lck*-Cre;*Ptpn1*$^{fl/fl}$ mice were left untreated or stimulated with α-CD3 (5 μg/ml) and crosslinked with α-hamster IgG (20 μg/ml) for the indicated times. Cell lysates were resolved by SDS-PAGE and immunoblotted with antibodies for **a**) phosphotyrosine (p-Tyr) and **b**) phosphorylated SFKs [p-SFK (Y418)], phosphorylated ZAP-70 [p-ZAP-70 (Y493)/Syk (Y526)] and phosphorylated and activated ERK1/2 (p-ERK1/2) and re-probed for Lck, ZAP-70, ERK2, actin or vinculin. **c**) $CD8^+$ T cells from *Ptpn1*$^{fl/fl}$ and *Lck*-Cre;*Ptpn1*$^{fl/fl}$ mice were left untreated or stimulated with α-CD3 (1 μg/ml or 5 μg/ml) and crosslinked with α-hamster IgG (20 μg/ml) for the indicated times. p-ERK1/2 was assessed by flow cytometry. **d**) Lymph node T cells from *Ptpn1*$^{fl/fl}$ and *Lck*-Cre;*Ptpn1*$^{fl/fl}$ mice were stained with fluorophore-conjugated antibodies against CD4, CD8 and CD44 and loaded with Fluo-4AM and then stimulated with α-CD3 (1 μg/ml) and crosslinked with α-hamster IgG (20 μg/ml). Intracellular calcium release was measured by flow cytometry and peak time, slope and peak intensity determined in naïve ($CD44^{lo}$) $CD4^+$ and $CD8^+$ T cells and in memory ($CD44^{hi}$) $CD8^+$ T cells. Representative results (means ± SEM) from two independent experiments are shown.

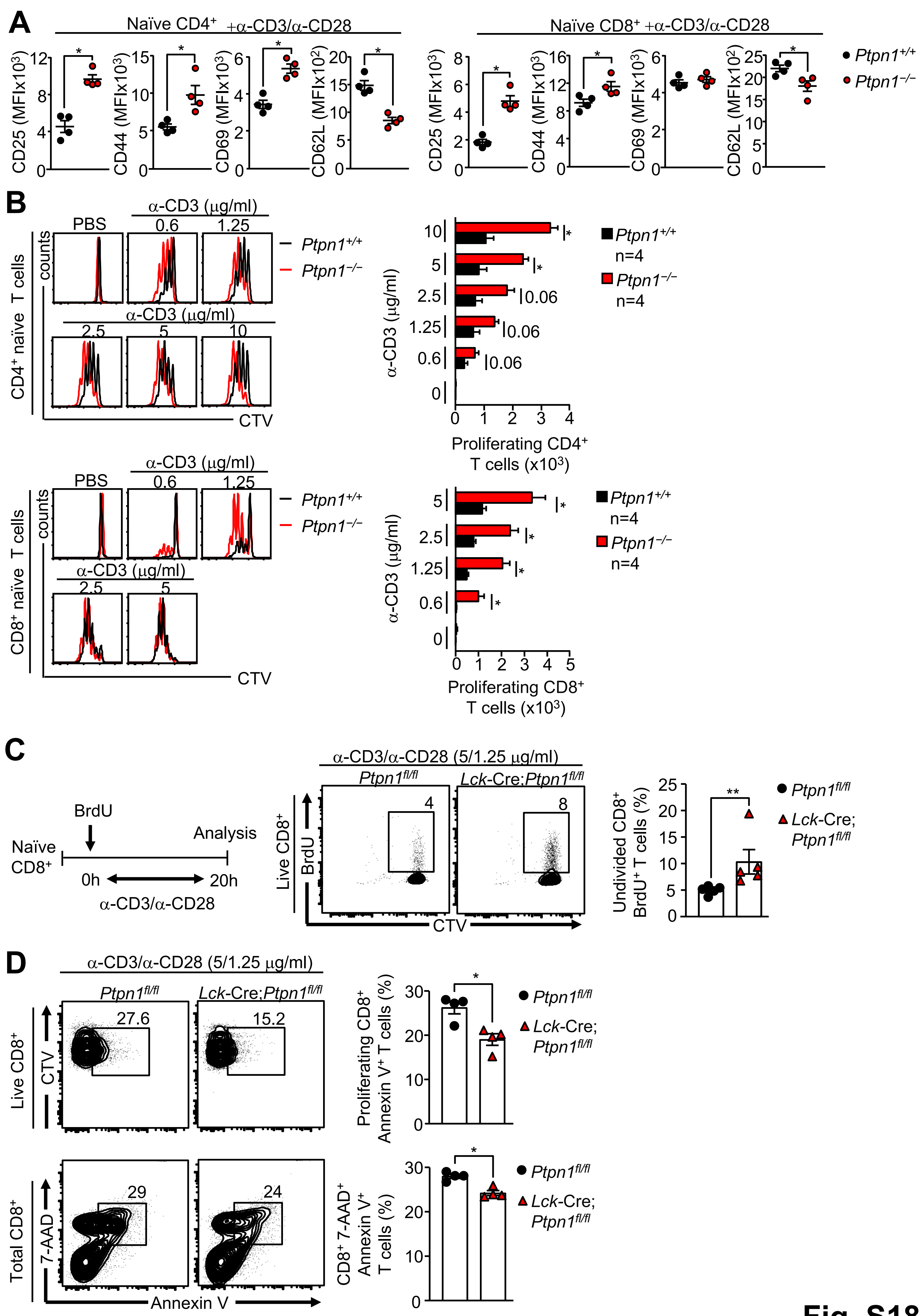

A
Naïve CD4+ +α-CD3/α-CD28
Naïve CD8+ +α-CD3/α-CD28
CD25 (MFIx10³)
CD44 (MFIx10³)
CD69 (MFIx10³)
CD62L (MFIx10²)
Ptpn1+/+
Ptpn1−/−
B
PBS
α-CD3 (μg/ml)
CD4+ naïve T cells
CD8+ naïve T cells
counts
CTV
Proliferating CD4+ T cells (x10³)
Proliferating CD8+ T cells (x10³)
n=4
0.06
C
BrdU
Naïve CD8+
Analysis
0h
20h
α-CD3/α-CD28
α-CD3/α-CD28 (5/1.25 μg/ml)
Ptpn1fl/fl
Lck-Cre;Ptpn1fl/fl
Live CD8+
Undivided CD8+ BrdU+ T cells (%)
D
Proliferating CD8+ Annexin V+ T cells (%)
Total CD8+
7-AAD
Annexin V
CD8+ 7-AAD+ Annexin V+ T cells (%)


Fig. S18

***Figure S18. PTP1B deletion enhances T cell activation, proliferation and survival.*** **a**) Naïve $CD8^+$ T cells from *Ptpn1*$^{+/+}$ and *Ptpn1* $^{-/-}$ mice were stimulated with plate-bound α-CD3 (2.5 μg/ml) and α-CD28 (1.25 μg/ml) for 48 h and CD25, CD44, CD69 and CD62L MFIs determined by flow cytometry. **b**) CTV-labelled naïve $CD4^+$ and $CD8^+$ T cells from *Ptpn1*$^{+/+}$ and *Ptpn1* $^{-/-}$ mice were stimulated with the indicated concentrations of plate-bound α-CD3 and α-CD28 (1.25 μg/ml) for 72 h and T cell proliferation (CTV dilution) was determined by flow cytometry. Total cell numbers were determined by adding a known amount of Flow-Count Fluorospheres before analysis. **c**) CTV-labelled naïve $CD8^+$ T cells from *Ptpn1*$^{fl/fl}$ and *Lck*-Cre;*Ptpn1*$^{fl/fl}$ mice were stimulated with plate-bound α-CD3 (5 μg/ml) plus α-CD28 (1.25 μg/ml) for 20 h in the presence of BrdU (10 μM) and the proportion of undivided $BrdU^+$ T cells was determined by flow cytometry. **d**) CTV-labelled naïve $CD8^+$ T cells from *Ptpn1*$^{fl/fl}$ and *Lck*-Cre;*Ptpn1*$^{fl/fl}$ mice were stimulated with plate-bound α-CD3 (5 μg/ml) plus α-CD28 (1.25 μg/ml) for 72 h and the proportions of proliferating apoptotic 7-$AAD^-$Annexin $V^+$ and necrotic/apoptotic 7-$AAD^+$Annexin $V^+$ T cells were determined by flow cytometry. Representative results (means ± SEM) from two independent experiments are shown. Significance in (a-d) was determined using 2-tailed Mann-Whitney U Test.

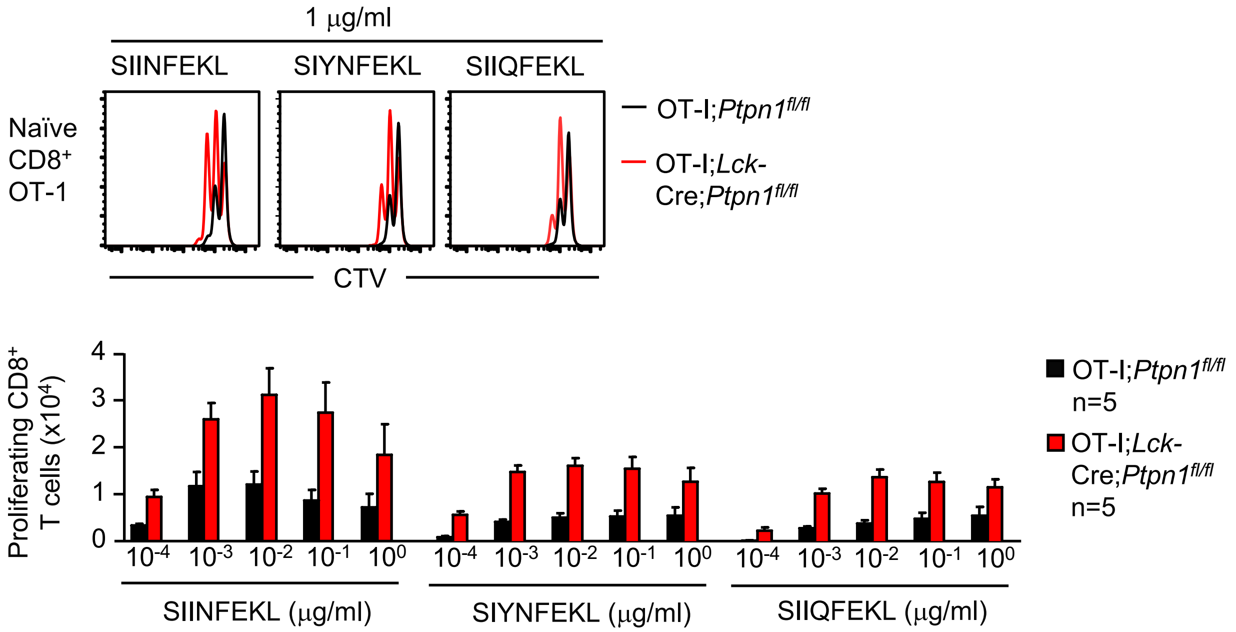
1 μg/ml
SIINFEKL
SIYNFEKL
SIIQFEKL
Naïve CD8+ OT-1
OT-I;Ptpn1fl/fl
OT-I;Lck-Cre;Ptpn1fl/fl
CTV
Proliferating CD8+ T cells (x10^4)
4
3
2
1
0
10^-4
10^-3
10^-2
10^-1
10^0
SIINFEKL (μg/ml)
SIYNFEKL (μg/ml)
SIIQFEKL (μg/ml)
OT-I;Ptpn1fl/fl n=5
OT-I;Lck-Cre;Ptpn1fl/fl n=5

**Fig. S19**

***Figure S19. PTP1B deletion enhances the activation of OT-I CD8*** $^{+}$ ***T cells by altered peptide ligands.*** CTV-labelled naïve CD8$^{+}$ T cells from OT-I;*Ptpn1*$^{fl/fl}$ and OT-I;*Lck*-Cre;*Ptpn1*$^{fl/fl}$ mice were stimulated with SIINFEKL (N4), SIYNFEKL (Y3) and SIIQFEKL (Q4) for 48 h and T cell proliferation (CTV dilution) determined by flow cytometry. Total cell numbers were determined by adding a known amount of Flow-Count Fluorospheres before analysis. Representative results (means ± SEM) from two independent experiments are shown.

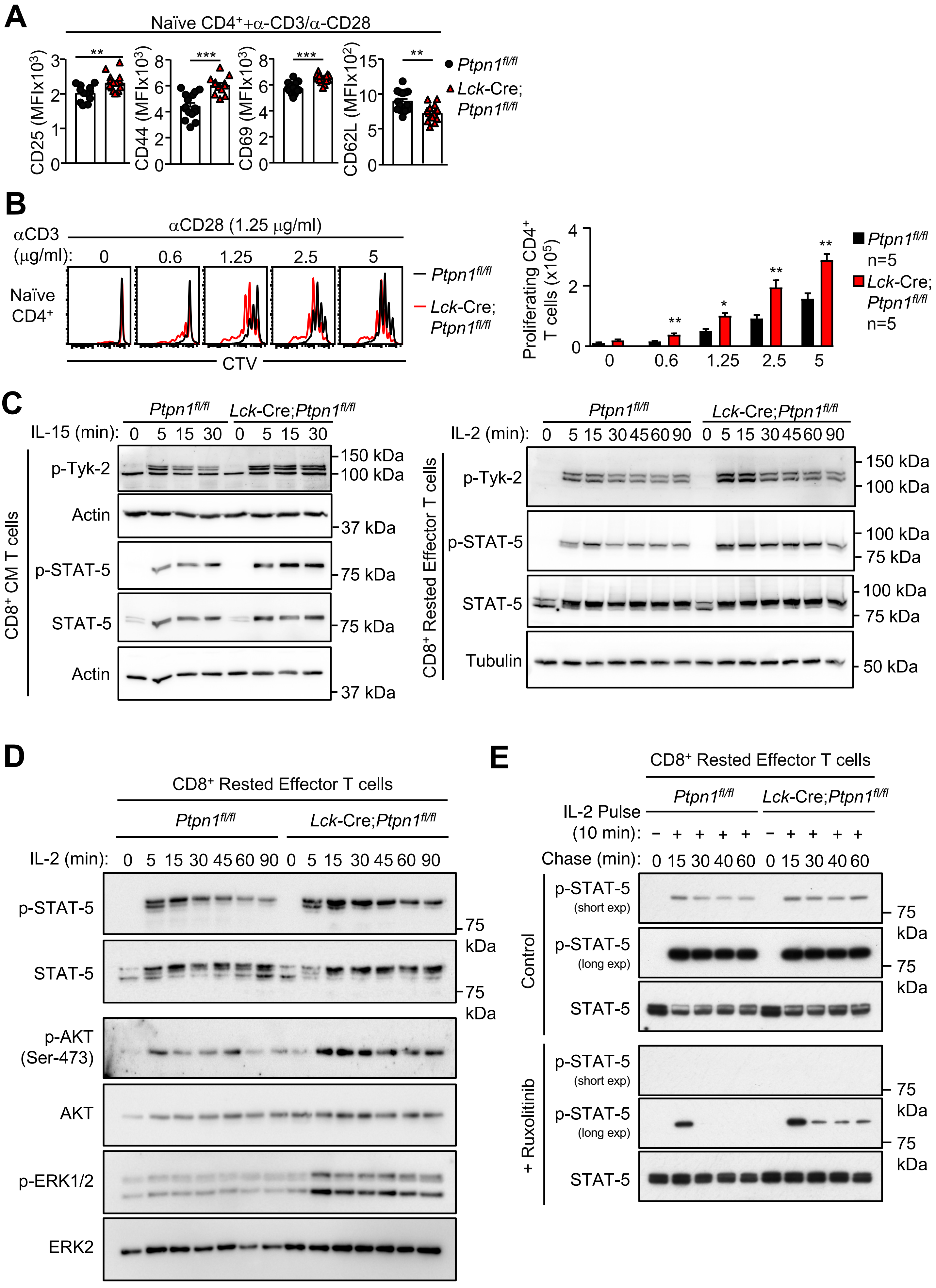
A
Naïve CD4+ + α-CD3/α-CD28
CD25 (MFIx10^3)
CD44 (MFIx10^3)
CD69 (MFIx10^3)
CD62L (MFIx10^2)
**
***
***
**
Ptpn1fl/fl
Lck-Cre; Ptpn1fl/fl
B
αCD3 (μg/ml):
αCD28 (1.25 μg/ml)
0
0.6
1.25
2.5
5
Naïve CD4+
CTV
Ptpn1fl/fl
Lck-Cre; Ptpn1fl/fl
Proliferating CD4+ T cells (x10^5)
Ptpn1fl/fl n=5
Lck-Cre; Ptpn1fl/fl n=5
C
Ptpn1fl/fl
Lck-Cre;Ptpn1fl/fl
IL-15 (min): 0 5 15 30 0 5 15 30
CD8+ CM T cells
p-Tyk-2
Actin
p-STAT-5
STAT-5
Actin
150 kDa
100 kDa
37 kDa
75 kDa
75 kDa
37 kDa
IL-2 (min): 0 5 15 30 45 60 90 0 5 15 30 45 60 90
CD8+ Rested Effector T cells
p-Tyk-2
p-STAT-5
STAT-5
Tubulin
150 kDa
100 kDa
100 kDa
75 kDa
100 kDa
75 kDa
50 kDa
D
CD8+ Rested Effector T cells
Ptpn1fl/fl
Lck-Cre;Ptpn1fl/fl
IL-2 (min): 0 5 15 30 45 60 90 0 5 15 30 45 60 90
p-STAT-5
STAT-5
p-AKT (Ser-473)
AKT
p-ERK1/2
ERK2
75 kDa
75 kDa
E
CD8+ Rested Effector T cells
Ptpn1fl/fl
Lck-Cre;Ptpn1fl/fl
IL-2 Pulse (10 min): − + + + + − + + + +
Chase (min): 0 15 30 40 60 0 15 30 40 60
Control
p-STAT-5 (short exp)
p-STAT-5 (long exp)
STAT-5
+ Ruxolitinib
p-STAT-5 (short exp)
p-STAT-5 (long exp)
STAT-5
75 kDa
75 kDa
75 kDa
75 kDa

**Fig. S20**

***Figure S20. PTP1B deletion enhances T cell activation, proliferation, survival and STAT-5 signaling*** (related to Fig. 4)**.** **a**) Naïve ($CD44^{lo}CD62L^{hi}$) $CD4^+$ T cells from *Ptpn1*$^{fl/fl}$ and *Lck*-Cre;*Ptpn1*$^{fl/fl}$ were stimulated with plate-bound α-CD3 (2.5 μg/ml) and α-CD28 (1.25 μg/ml) for 48 h and CD25, CD44, CD69 and CD62L levels determined by flow cytometry. **b**) CTV-labelled naïve $CD4^+$ ($CD44^{lo}CD62L^{hi}$) T cells from *Ptpn1*$^{fl/fl}$ and *Lck*-Cre;*Ptpn1*$^{fl/fl}$ mice were stimulated with the indicated concentrations of plate-bound α-CD3 and α-CD28 (1.25 μg/ml) for 72 h and T cell proliferation (CTV dilution) and the number of resulting T cells determined by flow cytometry. **c**) FACS-purified central memory ($CD44^{hi}CD62L^{hi}$) and *in vitro* generated effector $CD8^+$ T cells were stimulated with IL-15 and IL-2 respectively for the indicated times. Tyk-2 Y1054/Y1055 phosphorylation (p-Tyk-2) and p-STAT-5 were assessed by immunoblotting. **d**) *In vitro* generated effector $CD8^+$ T cells were stimulated with IL-2 for the indicated times and. p-STAT-5, p-ERK1/2 and p-AKT were assessed by immunoblotting. **e**) *In vitro* generated effector $CD8^+$ T cells were pulsed with IL-2 for 10 min. Cells were washed and incubated with or without Ruxolitinib (250 nM) for the indicated times and p-STAT-5 assessed by immunoblotting. Representative results (means ± SEM) from at least two independent experiments are shown. Significance in (a-b) was determined using 2-tailed Mann-Whitney U Test.

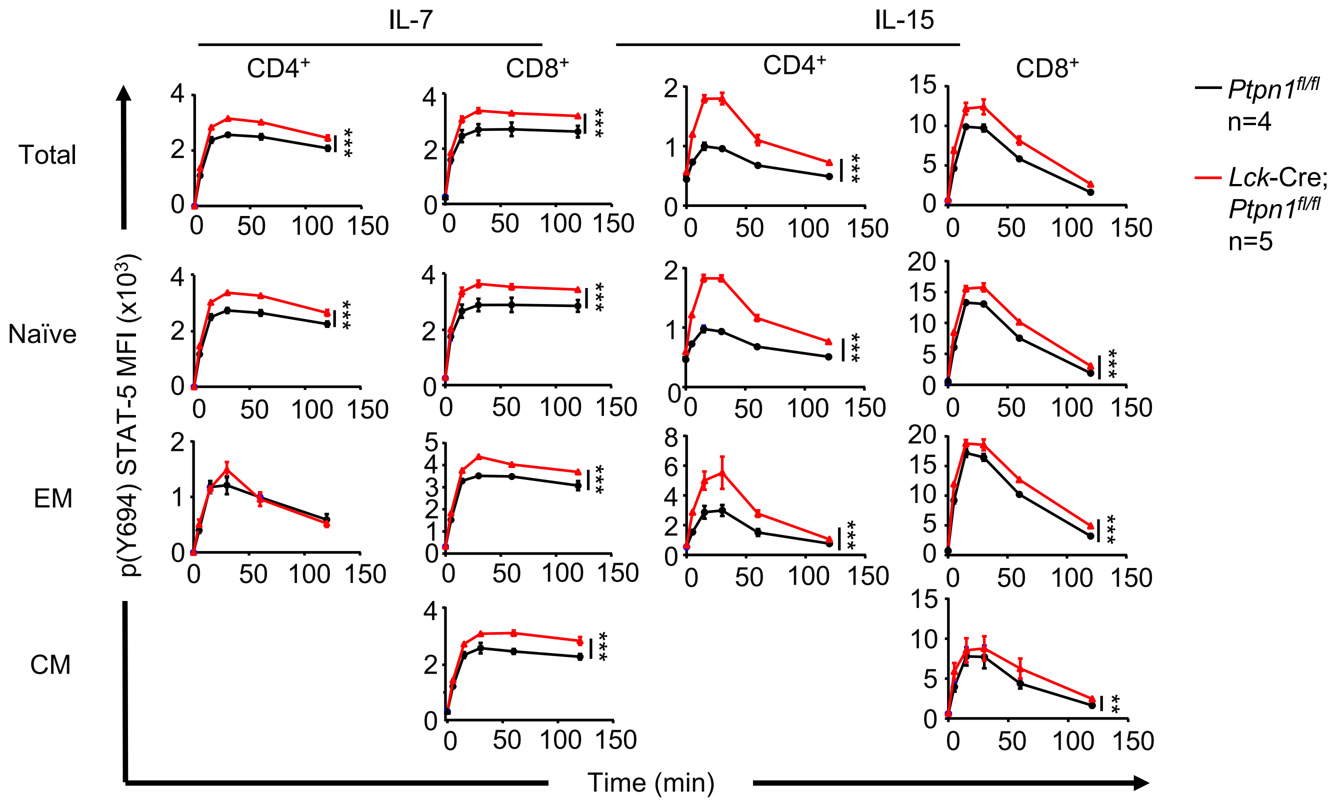
IL-7
IL-15
CD4+
CD8+
CD4+
CD8+
Total
Naïve
EM
CM
p(Y694) STAT-5 MFI (x10³)
Time (min)
Ptpn1fl/fl
n=4
Lck-Cre;
Ptpn1fl/fl
n=5
***
**

Fig. S21

***Figure S21. PTP1B-deficiency enhances IL-7 and IL-15-induced STAT-5 signaling in T cells.*** T cells were stimulated with IL-7 (5 ng/ml) or IL-15 (20 ng/ml) for the indicated times and intracellular STAT-5 Y694 phosphorylation (p-STAT-5) in total $CD4^+$ and $CD8^+$ T cells, naïve ($CD44^{lo}CD62L^{hi}$), central memory (CM; $CD44^{hi}CD62L^{hi}$) and effector memory (EM; $CD44^{hi}CD62L^{lo}$) T cells was assessed by flow cytometry. Significance was determined using 2-way ANOVA Test.

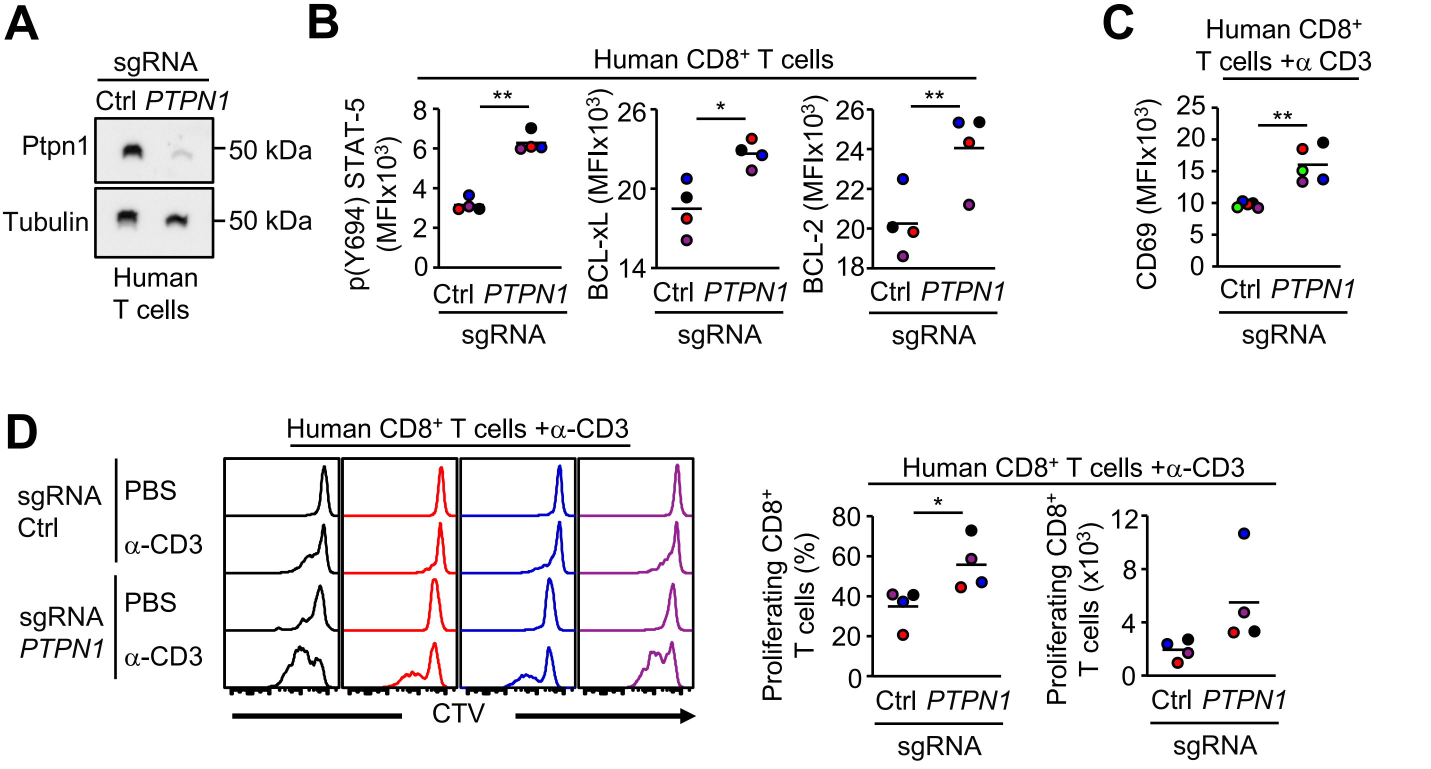
A
sgRNA
Ctrl PTPN1
Ptpn1
Tubulin
50 kDa
50 kDa
Human T cells
B
Human CD8+ T cells
p(Y694) STAT-5 (MFIx10^3)
BCL-xL (MFIx10^3)
BCL-2 (MFIx10^3)
Ctrl PTPN1
sgRNA
C
Human CD8+ T cells +α CD3
CD69 (MFIx10^3)
Ctrl PTPN1
sgRNA
D
Human CD8+ T cells +α-CD3
sgRNA Ctrl
sgRNA PTPN1
PBS
α-CD3
CTV
Proliferating CD8+ T cells (%)
Proliferating CD8+ T cells (x10^3)
Ctrl PTPN1
sgRNA

Fig. S22

***Figure S22. PTP1B deletion promotes STAT-5 signaling and the activation and proliferation of human T cells.*** CRISPR RNP-based gene-editing was used to delete PTP1B in human PBMC-derived T cells (PBMCs stimulated with OKT3 and IL-2 for 72 h). Control and PTP1B-deficient human T cells were processed for **a**) immunoblotting and **b**) intracellular p-STAT-5, BCL-xL or BCL-2 analysis by flow cytometry. **c**) Control and PTP1B-deficient PBMC-derived human T cells were stimulated with plate-bound α-CD3 (OKT3) overnight and CD69 (MFIs) levels analysed by flow cytometry. **d**) CTV-labelled control and PTP1B-deficient PBMC-derived human T cells were stimulated with plate-bound α-CD3 (OKT3) for 5 days and T cell proliferation (CTV dilution) was assessed by flow cytometry. Representative results (means ± SEM) from at least two independent experiments are shown. In (b-d) significance was determined using paired Student's t test.

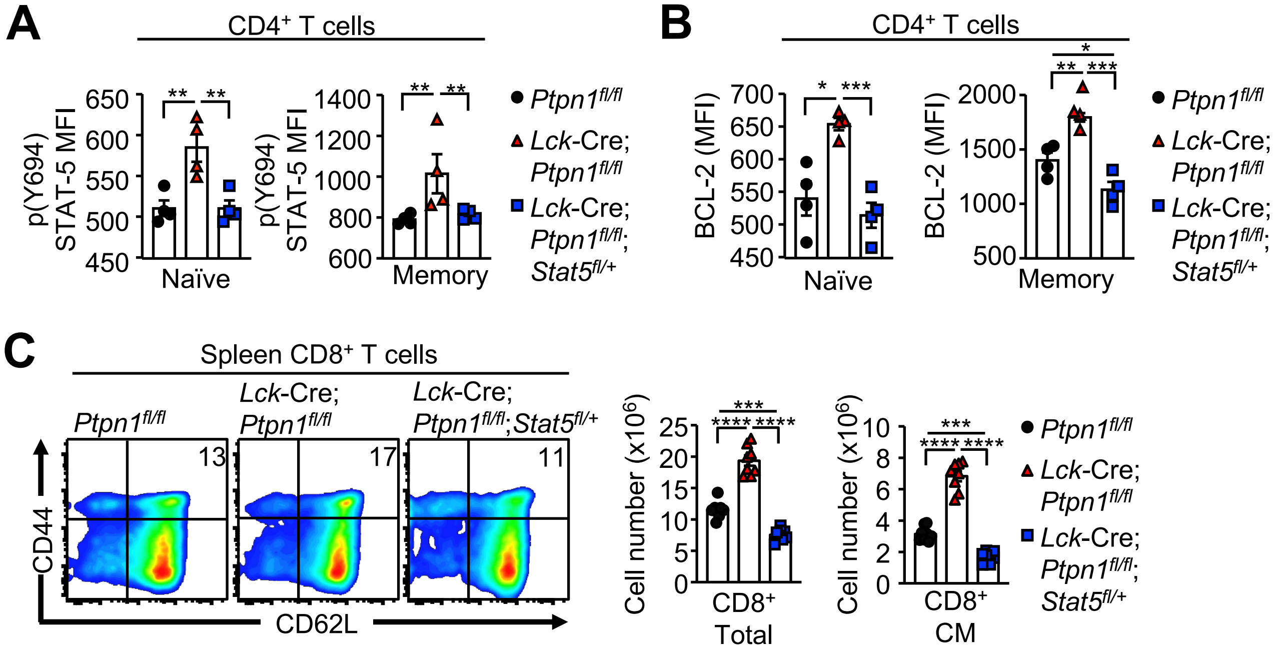
A
CD4+ T cells
p(Y694) STAT-5 MFI
Naïve
Memory
B
CD4+ T cells
BCL-2 (MFI)
Naïve
Memory
C
Spleen CD8+ T cells
Ptpn1fl/fl
Lck-Cre; Ptpn1fl/fl
Lck-Cre; Ptpn1fl/fl;Stat5fl/+
13
17
11
CD44
CD62L
Cell number (x10^6)
CD8+ Total
CD8+ CM
Ptpn1fl/fl
Lck-Cre; Ptpn1fl/fl
Lck-Cre; Ptpn1fl/fl; Stat5fl/+

**Fig. S23**

***Figure S23. PTP1B-deficiency enhances STAT-5-dependent BCL-2 expression and T cell development*** (related to Fig. 5)**.** **a**) Basal intracellular p-STAT-5 and **b**) BCL-2 were assessed in $CD44^{lo}$ and $CD44^{hi}$ $CD4^{+}$ T cells from *Ptpn1*$^{fl/fl}$, *Lck*-Cre;*Ptpn1*$^{fl/fl}$ and *Lck*-Cre;*Ptpn1*$^{fl/fl}$*Stat5*$^{fl/+}$ mice. **c**) The proportion and number of total and $CD8^{+}$ central memory (CM; $CD44^{hi}CD62L^{hi}$) T cells in the spleens from *Ptpn1*$^{fl/fl}$, *Lck*-Cre;*Ptpn1*$^{fl/fl}$ and *Lck*-Cre;*Ptpn1*$^{fl/fl}$*Stat5*$^{fl/+}$ mice were determined by flow cytometry. Representative results (means ± SEM) from at least two independent experiments are shown. In (a-c) significance was determined using 1-way ANOVA Test.

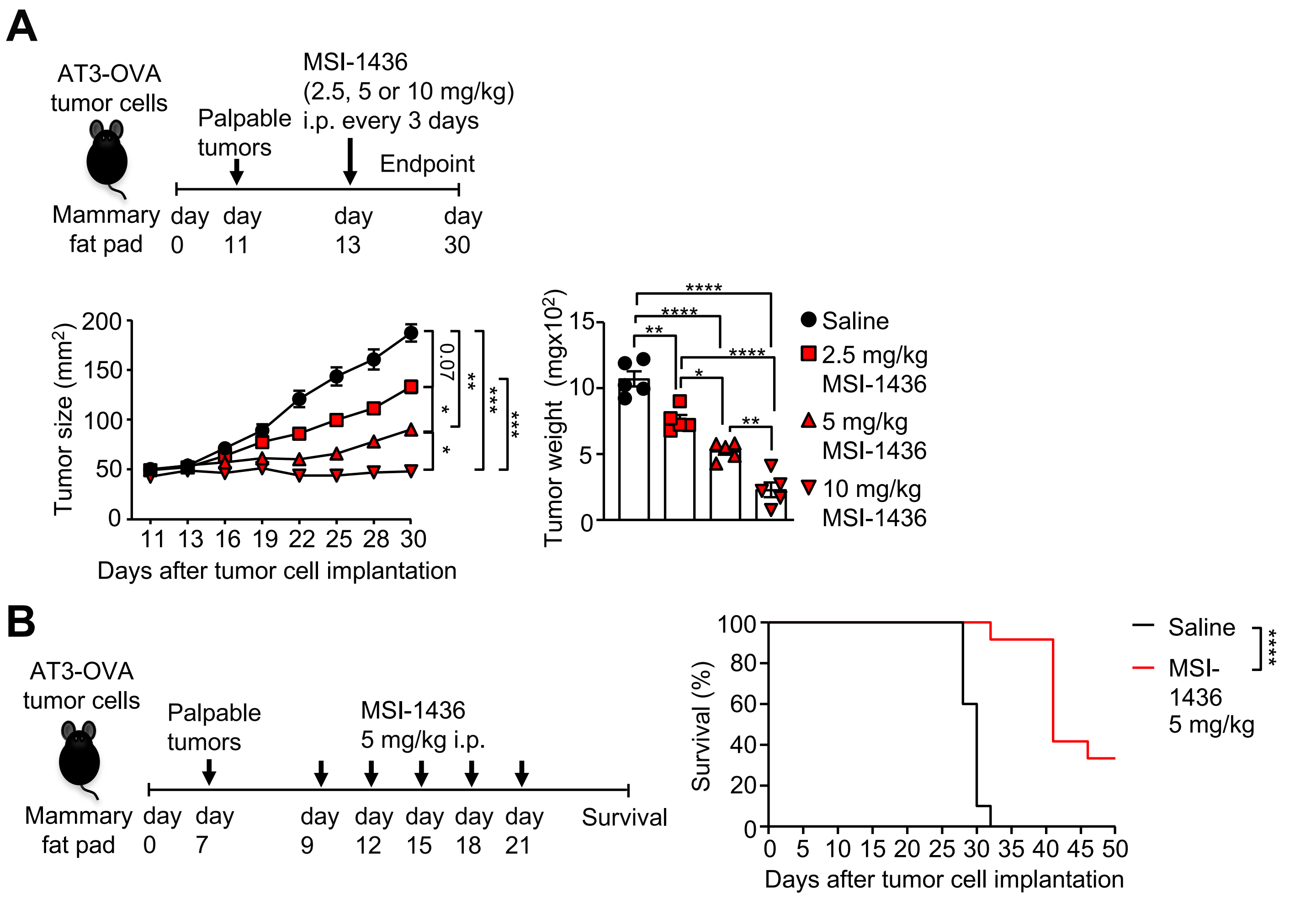
A
AT3-OVA
tumor cells
Mammary
fat pad
Palpable
tumors
MSI-1436
(2.5, 5 or 10 mg/kg)
i.p. every 3 days
Endpoint
day
0
day
11
day
13
day
30
Tumor size (mm²)
Days after tumor cell implantation
0.07
Tumor weight (mgx10²)
Saline
2.5 mg/kg
MSI-1436
5 mg/kg
MSI-1436
10 mg/kg
MSI-1436
B
AT3-OVA
tumor cells
Mammary
fat pad
Palpable
tumors
MSI-1436
5 mg/kg i.p.
day
0
day
7
day
9
day
12
day
15
day
18
day
21
Survival
Survival (%)
Days after tumor cell implantation
Saline
MSI-
1436
5 mg/kg

Fig. S24

***Figure S24. MSI-1436 represses tumor growth and promotes survival*** (related to Fig. 6)**. a**) AT3-OVA tumor cells were implanted into the fourth inguinal mammary fat pads of C57BL/6 mice. Mice were treated with MSI-1436 (2.5, 5 and 10 mg/kg intraperitoneally) or saline on days 13, 16, 19, 22, 25 and 28 after tumor cell implantation. Tumor growth and weights were measured. **b**) AT3-OVA tumor cells were implanted into the fourth inguinal mammary fat pads of C57BL/6 mice. Mice were treated with MSI-1436 (5 mg/kg intraperitoneally) or saline on days 9, 12, 15, 18 and 21 after tumor cell implantation and survival monitored. Representative results (means ± SEM) from at least two independent experiments are shown. For tumor sizes in (a) significance was determined using 2-way ANOVA Test and for tumor weights in (a) significance was determined using 1-way ANOVA Test. In (b) significance was determined using Log-rank (Mantel-Cox) test.

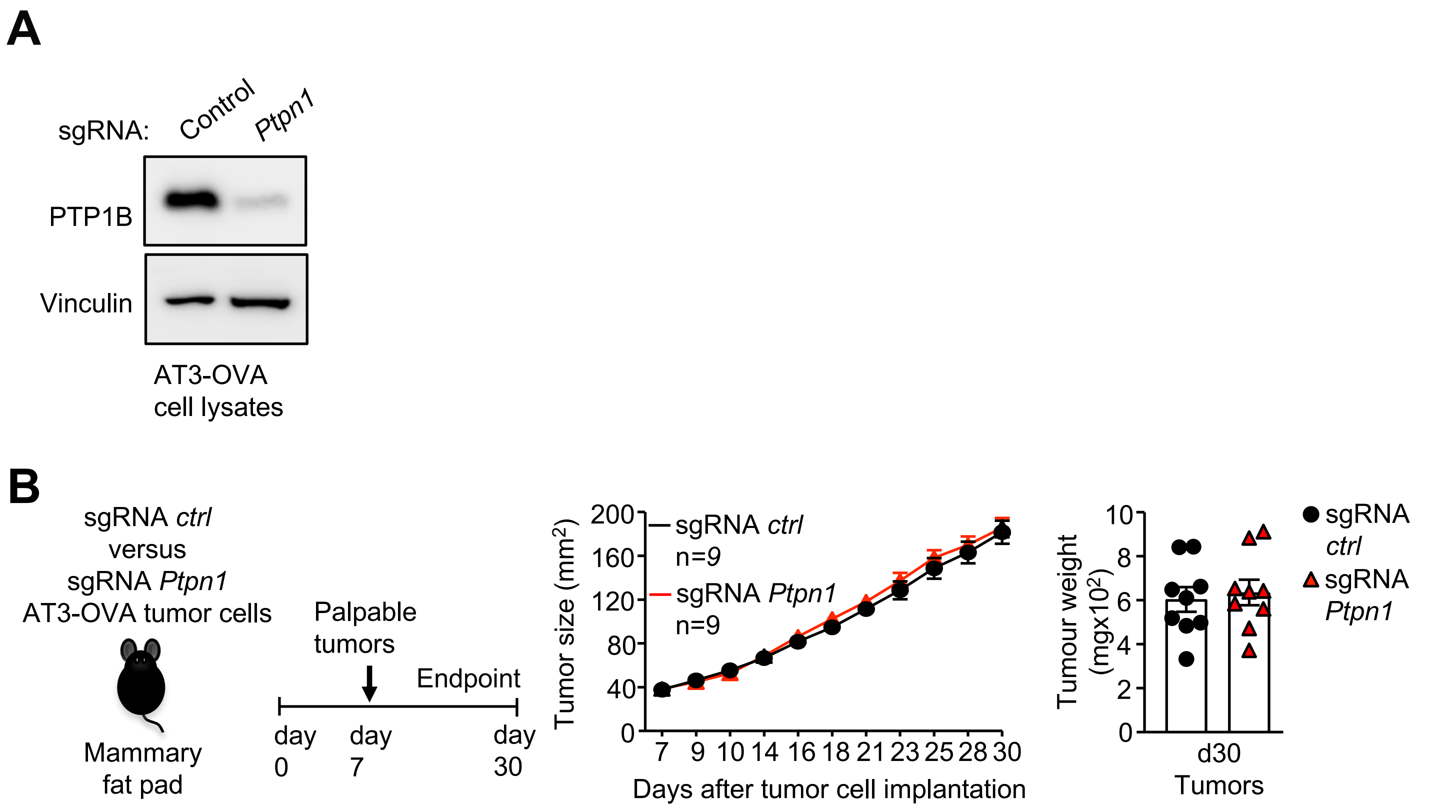
A
sgRNA:
Control
Ptpn1
PTP1B
Vinculin
AT3-OVA
cell lysates
B
sgRNA ctrl
versus
sgRNA Ptpn1
AT3-OVA tumor cells
Mammary
fat pad
Palpable
tumors
Endpoint
day
0
day
7
day
30
Tumor size (mm²)
sgRNA ctrl
n=9
sgRNA Ptpn1
n=9
Days after tumor cell implantation
Tumour weight
(mgx10²)
d30
Tumors
sgRNA
ctrl
sgRNA
Ptpn1

Fig. S25

***Figure S25. PTP1B deletion does not affect the growth of AT3-OVA tumors***. CRISPR RNP-based gene-editing was used to delete PTP1B in AT3-OVA cells. AT3-OVA cells were transfected with recombinant Cas9 pre-complexed with control (ctrl) non-targeting sgRNAs or those targeting *Ptpn1*. **a**) PTP1B deletion was assessed by immunoblotting. **b**) Control AT3-OVA cells or those in which PTP1B had been deleted by CRISPR-Cas9 gene editing were injected into the fourth inguinal mammary fat pads of female C57BL/6 mice and tumor growth monitored. Representative results (means ± SEM) from at least two independent experiments are shown.

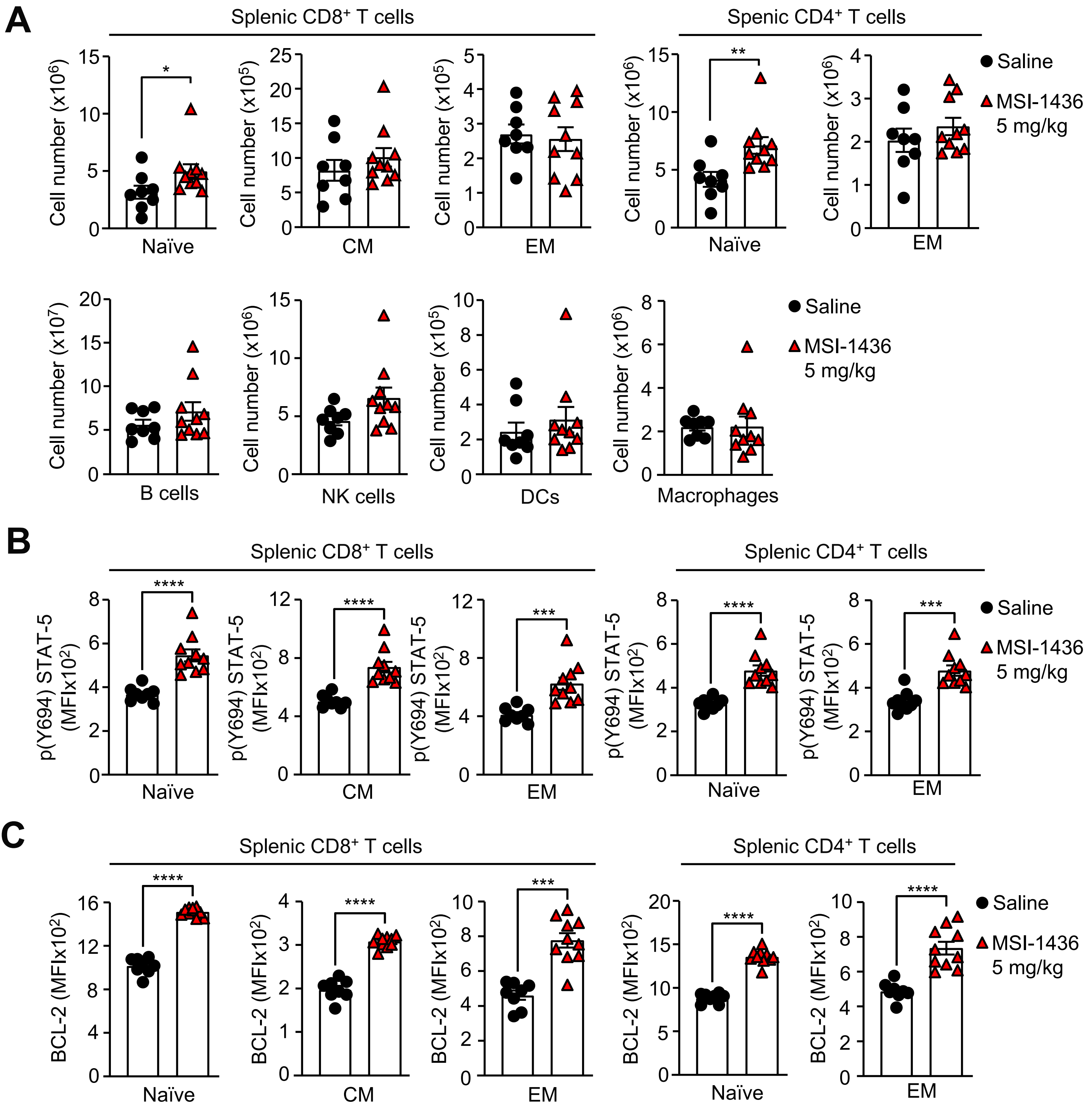
A
Splenic CD8+ T cells
Spenic CD4+ T cells
Cell number (x10^6)
Cell number (x10^5)
Cell number (x10^7)
Naïve
CM
EM
B cells
NK cells
DCs
Macrophages
Saline
MSI-1436
5 mg/kg
B
Splenic CD8+ T cells
Splenic CD4+ T cells
p(Y694) STAT-5
(MFIx10^2)
C
BCL-2 (MFIx10^2)

Fig. S26

***Figure S26. MSI-1436 increases naïve T cell numbers and promotes STAT-5 signaling in naïve, central memory and effector/memory T cells.*** **a-c**) AT3-OVA tumor cells were implanted into the fourth inguinal mammary fat pads of C57BL/6 mice. Mice were treated with MSI-1436 (5 mg/kg intraperitoneally) or saline on days 13, 16, 19 and 22. On day 23 splenocytes were harvested and **a**) the numbers of naïve ($CD44^{lo}CD62L^{hi}$), central memory (CM; $CD44^{hi}CD62L^{hi}$) and effector memory (EM; $CD44^{hi}CD62L^{lo}$) $CD8^+$ and $CD4^+$ T cells , as well as $B220^+$ B cells, $NK1.1^+TCR\beta^-$ (NK) cells, $CD11c^+$ dendritic cells (DCs) and $CD11b^+F4/80^+$ macrophages were determined by flow cytometry. Alternatively, **b**) intracellular p-STAT-5 and BCL-2 levels were determined in naïve, CM and EM $CD8^+$ and $CD4^+$ T cells by flow cytometry. Results are means ± SEM) for the indicated number of mice. Significance was determined using 2-tailed Mann-Whitney U Test.

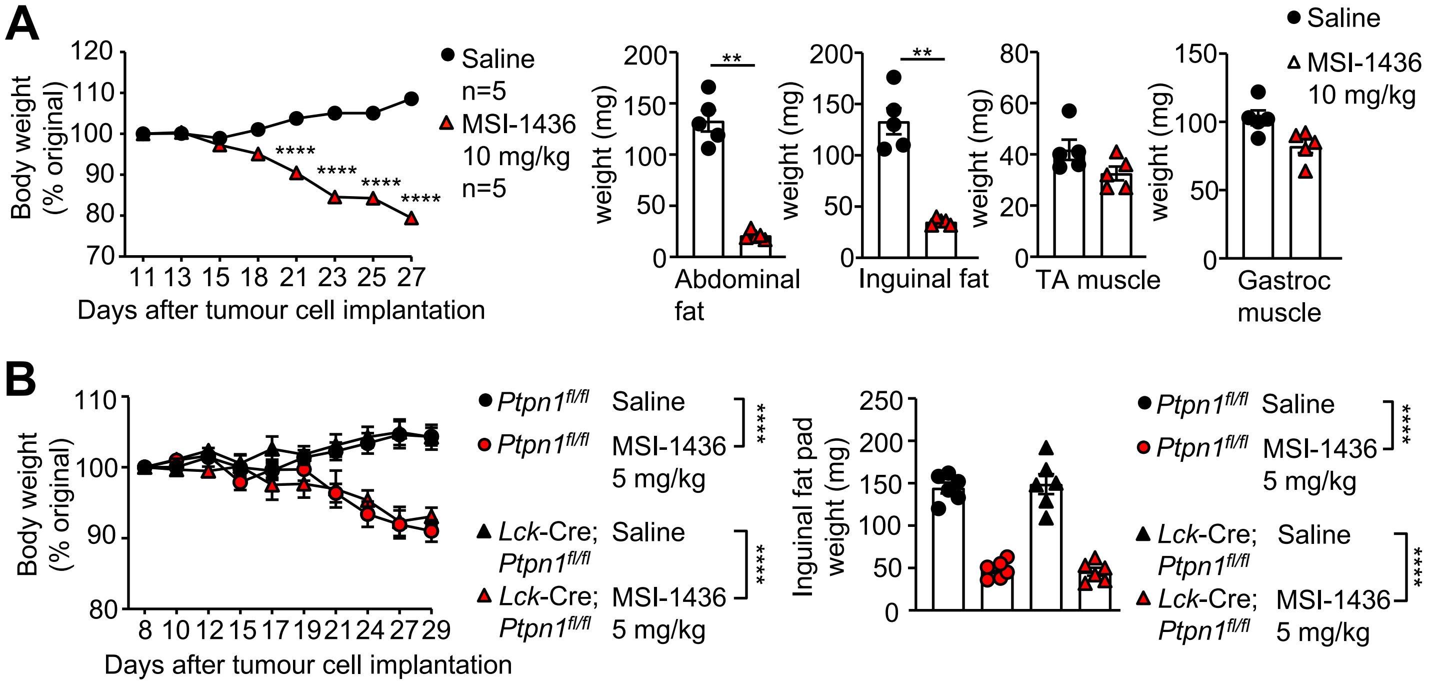
A
Body weight (% original)
Days after tumour cell implantation
Saline n=5
MSI-1436 10 mg/kg n=5
weight (mg)
Abdominal fat
Inguinal fat
TA muscle
Gastroc muscle
Saline
MSI-1436 10 mg/kg
B
Body weight (% original)
Days after tumour cell implantation
Ptpn1fl/fl Saline
Ptpn1fl/fl MSI-1436 5 mg/kg
Lck-Cre; Ptpn1fl/fl Saline
Lck-Cre; Ptpn1fl/fl MSI-1436 5 mg/kg
Inguinal fat pad weight (mg)

Fig. S27

***Figure S27. MSI-1436 promotes weight loss and decreases adiposity*** (related to Fig. 6)**. a**) AT3-OVA tumor cells were implanted into the fourth inguinal mammary fat pads of C57BL/6 mice. Mice were treated with MSI-1436 (10 mg/kg intraperitoneally) or saline on days 13, 16, 19, 22 and 25 after tumor cell implantation and body weights determined. At the completion of the experiment abdominal fat, inguinal fat, tibialis anterior (TA) muscle and gastrocnemius (Gastroc) muscle weights determined. **b**) AT3-OVA tumor cells were implanted orthotopically into the fourth inguinal mammary fat pads of *Ptpn1*$^{fl/fl}$ or *Lck*-Cre;*Ptpn1*$^{fl/fl}$ mice. After tumors were established, mice were treated every 3 days with saline or MSI-1436 (5 mg/kg intraperitoneally on days 14, 17, 20, 23 and 26) and body weights determined. At the completion of the experiment inguinal fat pad weights were determined. Representative results (means ± SEM) from at least two independent experiments are shown. Significances for body weights in (a, b) were determined using 2-way ANOVA Test. Significance for fat pad weights in (a, b) was determined using 2-tailed Mann-Whitney U Test.

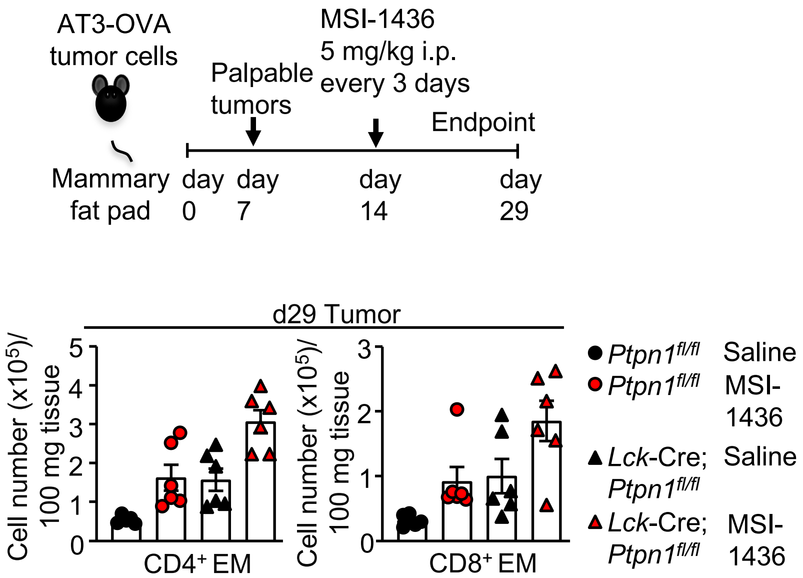
AT3-OVA
tumor cells
Mammary
fat pad
Palpable
tumors
MSI-1436
5 mg/kg i.p.
every 3 days
Endpoint
day 0
day 7
day 14
day 29
d29 Tumor
Cell number (x10^5)/
100 mg tissue
CD4+ EM
CD8+ EM
Ptpn1fl/fl Saline
Ptpn1fl/fl MSI-1436
Lck-Cre; Ptpn1fl/fl Saline
Lck-Cre; Ptpn1fl/fl MSI-1436

**Fig. S28**

***Figure S28. The inhibition of PTP1B with MSI-1436 promotes the infiltration of activated T cells into tumors*** (related to Fig. 6)**.** AT3-OVA tumor cells were implanted into the fourth inguinal mammary fat pads of *Ptpn1*$^{fl/fl}$ versus *Lck*-Cre;*Ptpn1*$^{fl/fl}$ mice. After tumors were established, mice were treated every 3 days with saline or MSI-1436 (5 mg/kg intraperitoneally on days 14, 17, 20, 23, and 26) and tumor-infiltrating $CD4^+$ and $CD8^+$ effector memory (EM; $CD44^{hi}CD62L^{lo}$) T cell numbers were determined by flow cytometry. Representative results (means ± SEM) from at least two independent experiments are shown.

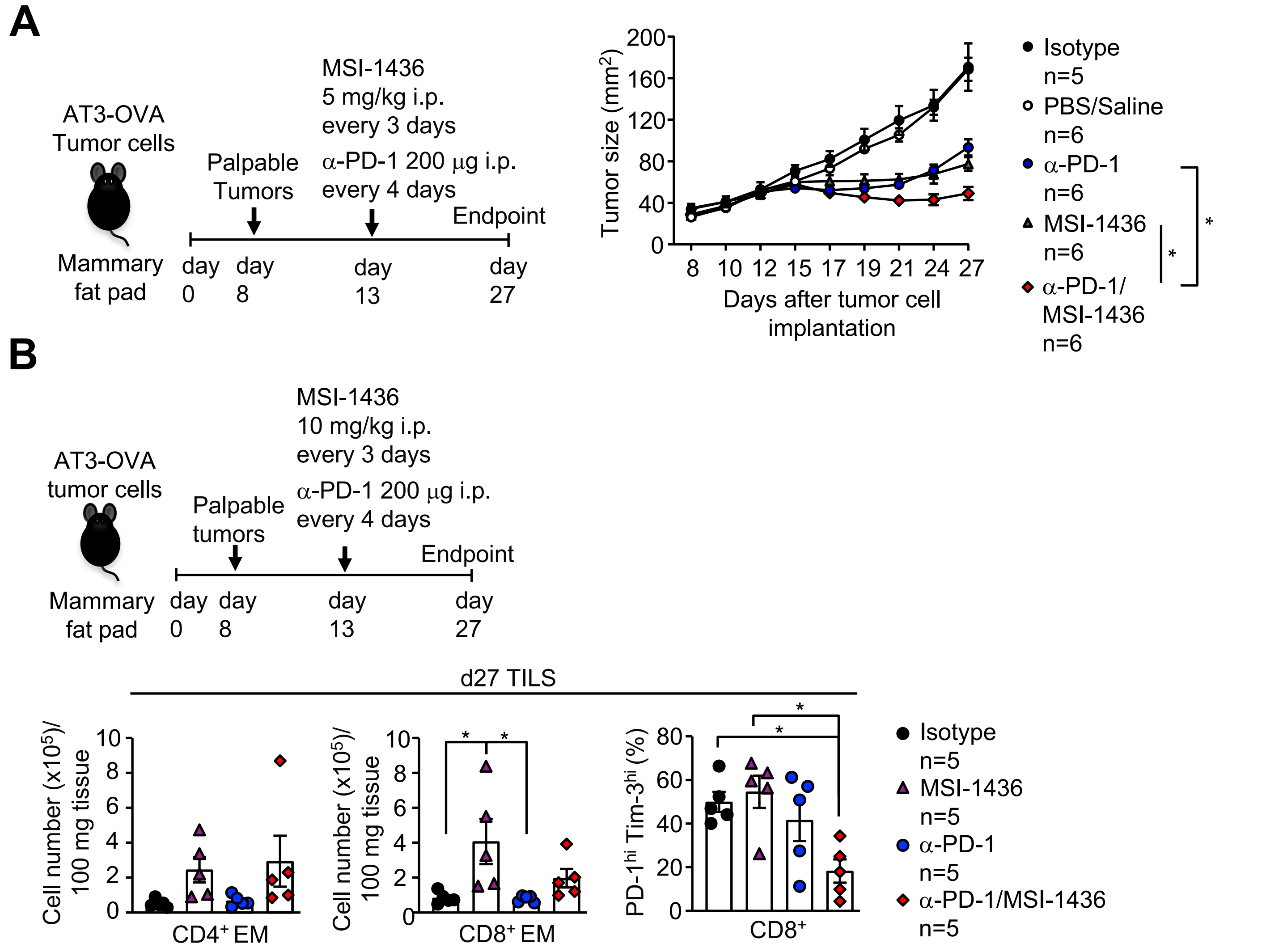
A
AT3-OVA
Tumor cells
Mammary
fat pad
Palpable
Tumors
MSI-1436
5 mg/kg i.p.
every 3 days
α-PD-1 200 μg i.p.
every 4 days
Endpoint
day 0
day 8
day 13
day 27
Tumor size (mm2)
Days after tumor cell implantation
Isotype n=5
PBS/Saline n=6
α-PD-1 n=6
MSI-1436 n=6
α-PD-1/MSI-1436 n=6
B
AT3-OVA
tumor cells
Palpable
tumors
MSI-1436
10 mg/kg i.p.
every 3 days
α-PD-1 200 μg i.p.
every 4 days
Endpoint
Mammary
fat pad
day 0
day 8
day 13
day 27
d27 TILS
Cell number (x105)/100 mg tissue
CD4+ EM
CD8+ EM
PD-1hi Tim-3hi (%)
CD8+
Isotype n=5
MSI-1436 n=5
α-PD-1 n=5
α-PD-1/MSI-1436 n=5

**Fig. S29**

***Figure S29. PTP1B inhibition with MSI-1436 enhances the response to PD-1 checkpoint blockade*** (related to Fig. 6)**.** **a**) AT3-OVA tumor cells were implanted into the fourth inguinal mammary fat pads of C57BL/6 mice. Mice were treated with MSI-1436 (5 mg/kg intraperitoneally on days 13, 16, 19, 22, and 25) and either α-PD-1 or isotype control (200 μg intraperitoneally in each case on days 13, 17, 21 and 24) alone or MSI-1436 plus α-PD-1 and tumor growth monitored. **b**) AT3-OVA tumor cells were implanted into the fourth inguinal mammary fat pads of C57BL/6 mice. Mice were treated with MSI-1436 (10 mg/kg intraperitoneally on days 13, 16, 19, 22, and 25) and either α-PD-1 or isotype control (200 μg intraperitoneally in each case on days 13, 17, 21 and 24) alone or MSI-1436 plus α-PD-1 and the number of tumor-infiltrating $CD4^+$ and $CD8^+$ effector memory (EM; $CD44^{hi}CD62L^{lo}$) T cells and the frequency of $CD8^+PD\text{-}1^{hi}Tim\text{-}3^{hi}$ T cells was determined by flow cytometry. Representative results (means ± SEM) from at least two independent experiments are shown. Significance in (a) was determined using 2-way ANOVA Test.

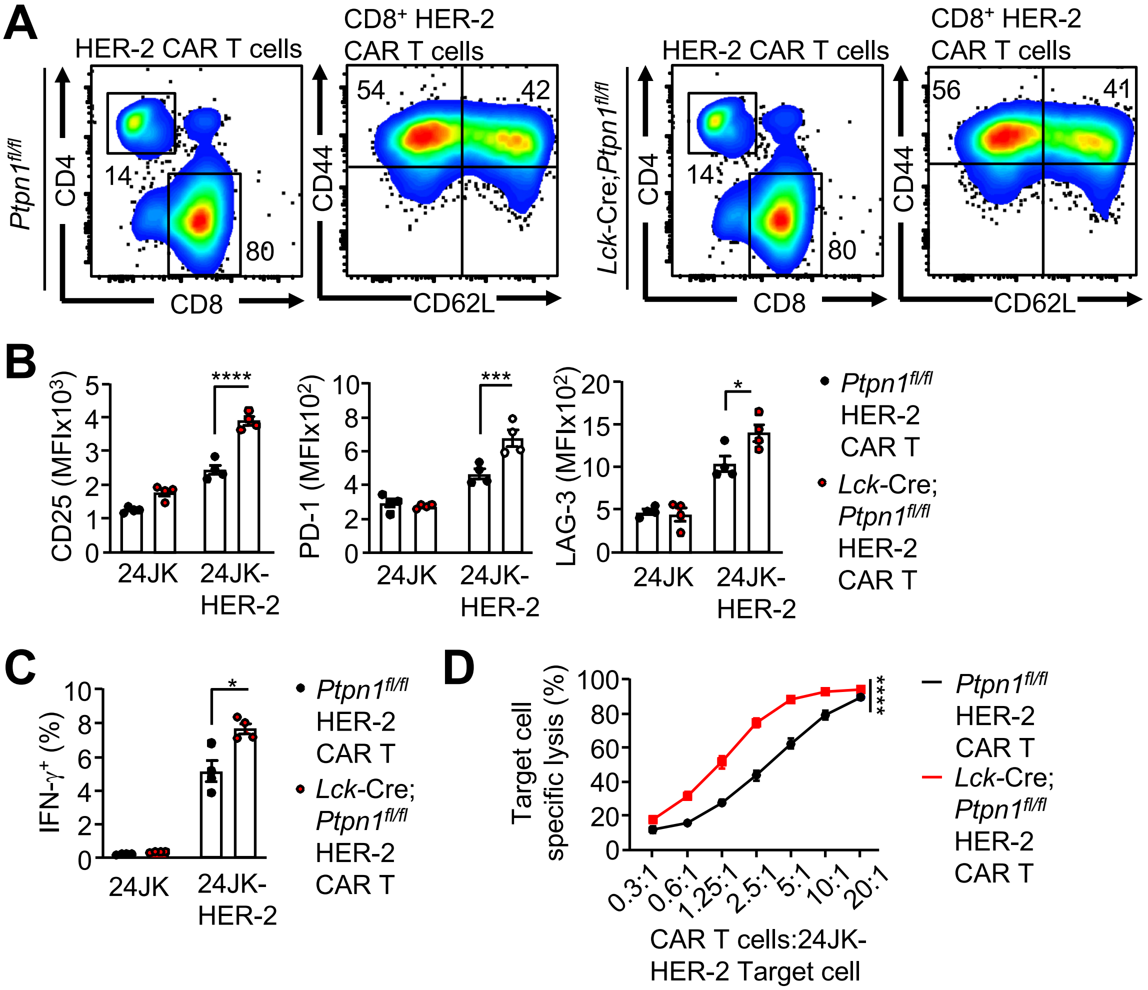
A
HER-2 CAR T cells
CD8⁺ HER-2 CAR T cells
Ptpn1fl/fl
CD4
CD8
14
80
CD44
CD62L
54
42
Lck-Cre;Ptpn1fl/fl
14
80
56
41
B
CD25 (MFIx10³)
PD-1 (MFIx10²)
LAG-3 (MFIx10²)
24JK
24JK-HER-2
****
***
*
Ptpn1fl/fl HER-2 CAR T
Lck-Cre; Ptpn1fl/fl HER-2 CAR T
C
IFN-γ⁺ (%)
*
D
Target cell specific lysis (%)
0.3:1
0.6:1
1.25:1
2.5:1
5:1
10:1
20:1
CAR T cells:24JK-HER-2 Target cell
****

Fig. S30

***Figure S30. PTP1B-deficiency enhances CAR T cell cytotoxicity ex vivo.*** **a**) HER-2 CAR T cells generated from *Ptpn1*$^{fl/fl}$ versus *Lck*-Cre;*Ptpn1*$^{fl/fl}$ splenocytes were assessed for the proportions of $CD4^+$ and $CD8^+$ total T cells as well as $CD44^{hi}CD62L^{hi}$ central memory versus $CD44^{hi}CD62L^{lo}$ effector/memory $CD8^+$ T cells by flow cytometry. **b**) HER-2-specific *Ptpn1*$^{fl/fl}$ versus *Lck*-Cre;*Ptpn1*$^{fl/fl}$ CAR T cells were incubated overnight with HER-2 expressing 24JK sarcoma cells (24JK-HER-2) or HER-2-negative 24JK sarcoma cells and CD25, PD-1 and LAG-3 MFIs on $CD8^+$ CAR T cells determined by flow cytometry. **c**) HER-2-specific *Ptpn1*$^{fl/fl}$ versus *Lck*-Cre;*Ptpn1*$^{fl/fl}$ CAR T cells were incubated for 4 h with HER-2 expressing 24JK sarcoma cells (24JK-HER-2) or HER-2 negative 24JK sarcoma cells and intracellular IFN-γ was determined by flow cytometry. **d**) *Ptpn1*$^{fl/fl}$ versus *Lck*-Cre;*Ptpn1*$^{fl/fl}$ HER-2 CAR T cells were incubated with 5 μM CTV-labelled ($CTV^{bright}$) 24JK-HER-2 and 0.5 μM CTV-labelled ($CTV^{dim}$) 24JK sarcoma cells. Antigen-specific target cell lysis was assessed after 4 h by monitoring for the depletion of $CTV^{bright}$ 24JK-HER-2 cells by flow cytometry. Representative results (means ± SEM) from at least two independent experiments are shown. Significance in (c) was determined using 2-tailed Mann-Whitney U Test. Significance in (b) was determined using one-way ANOVA Test. Significance in (d) was determined using 2-way ANOVA Test.

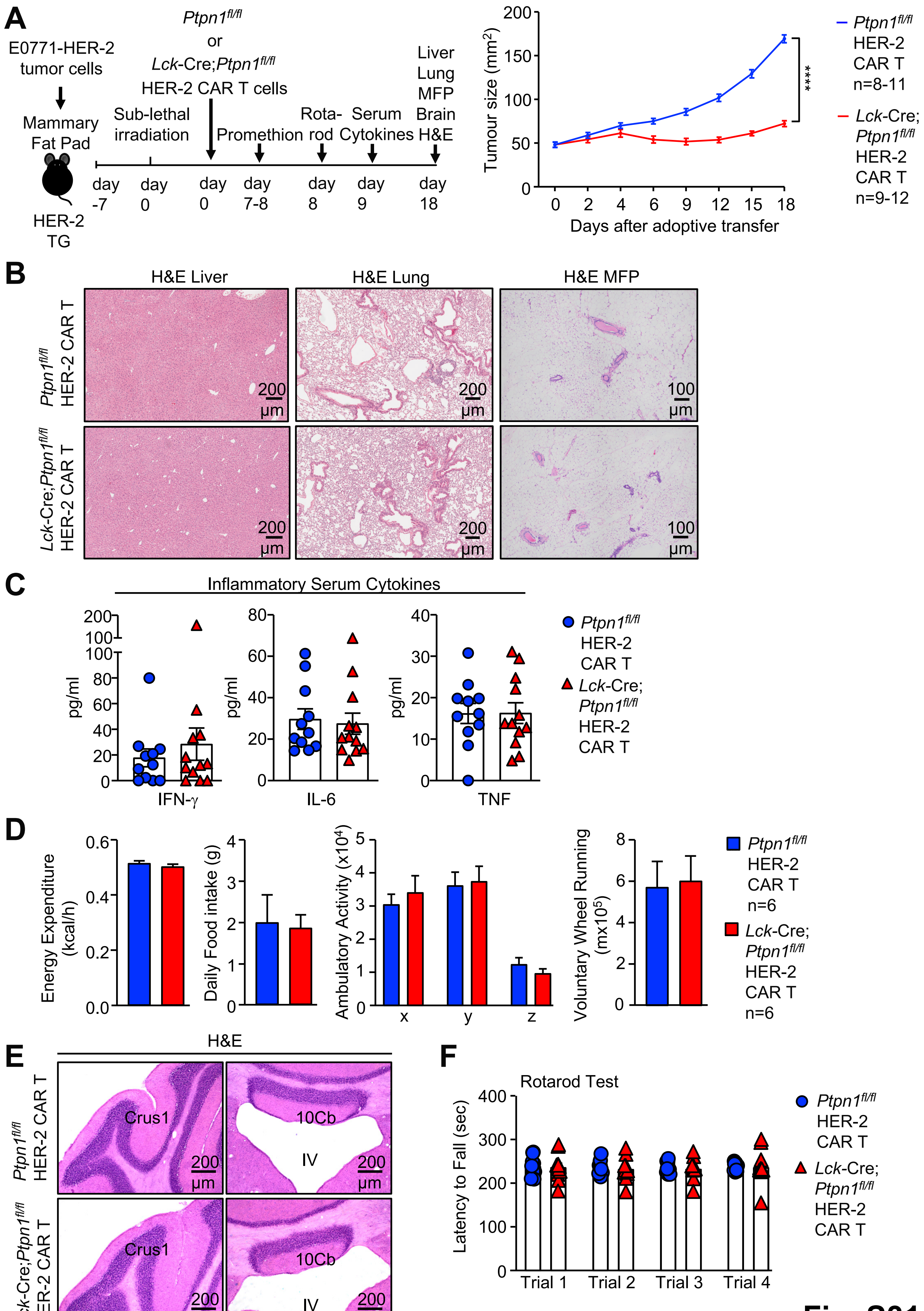
A
E0771-HER-2 tumor cells
Mammary Fat Pad
HER-2 TG
Sub-lethal irradiation
Ptpn1fl/fl or Lck-Cre;Ptpn1fl/fl HER-2 CAR T cells
Promethion
Rota-rod
Serum Cytokines
Liver Lung MFP Brain H&E
day -7
day 0
day 0
day 7-8
day 8
day 9
day 18
Tumour size (mm2)
Days after adoptive transfer
****
Ptpn1fl/fl HER-2 CAR T n=8-11
Lck-Cre; Ptpn1fl/fl HER-2 CAR T n=9-12
B
H&E Liver
H&E Lung
H&E MFP
Ptpn1fl/fl HER-2 CAR T
Lck-Cre;Ptpn1fl/fl HER-2 CAR T
200 µm
100 µm
C
Inflammatory Serum Cytokines
pg/ml
IFN-γ
IL-6
TNF
Ptpn1fl/fl HER-2 CAR T
Lck-Cre; Ptpn1fl/fl HER-2 CAR T
D
Energy Expenditure (kcal/h)
Daily Food intake (g)
Ambulatory Activity (x104)
Voluntary Wheel Running (mx105)
Ptpn1fl/fl HER-2 CAR T n=6
Lck-Cre; Ptpn1fl/fl HER-2 CAR T n=6
E
H&E
Crus1
10Cb
IV
200 µm
F
Rotarod Test
Latency to Fall (sec)
Trial 1
Trial 2
Trial 3
Trial 4
Ptpn1fl/fl HER-2 CAR T
Lck-Cre; Ptpn1fl/fl HER-2 CAR T

Fig. S31

***Figure S31. PTP1B deficiency in CAR T cells does not result in systemic inflammation or morbidity.*** **a-f**) HER-2-E0771 mammary tumor cells were injected into the fourth inguinal mammary fat pads of female HER-2 TG mice. Seven days after tumor injection, HER-2 TG mice received total body irradiation (4 Gy) followed by the adoptive transfer of 20x$10^6$ HER-2 CAR T cells generated from *Ptpn1*$^{fl/fl}$ or *Lck*-Cre;*Ptpn1*$^{fl/fl}$ splenocytes. **a**) Tumor growth was monitored. **b**) The livers, lungs and the contralateral mammary fat pads (MFP) from HER-2 TG mice administered *Ptpn1*$^{fl/fl}$ or *Lck*-Cre;*Ptpn1*$^{fl/fl}$ HER-2 CAR T cells were processed for histological assessment (hematoxylin and eosin: H&E). **c**) Serum cytokines in HER-2 TG mice administered *Ptpn1*$^{fl/fl}$ or *Lck*-Cre;*Ptpn1*$^{fl/fl}$ HER-2 CAR T cells were determined by flow cytometry using a BD Cytokine Bead Array (BD Biosciences). **d**) Energy expenditure, daily food intake, ambulatory activity and voluntary wheel running were assessed in HER-2 TG mice administered *Ptpn1*$^{fl/fl}$ or *Lck*-Cre;*Ptpn1*$^{fl/fl}$ HER-2 CAR T cells. **e**) Brains from HER-2 TG mice administered *Ptpn1*$^{fl/fl}$ or *Lck*-Cre;*Ptpn1*$^{fl/fl}$ HER-2 CAR T cells were fixed in paraformaldehyde and cerebella sections processed for histology (H&E) to assess gross tissue architecture. **f**) HER-2 TG mice administered *Ptpn1*$^{fl/fl}$ or *Lck*-Cre;*Ptpn1*$^{fl/fl}$ HER-2 CAR T cells were subjected to a rotarod test and the latency to fall determined in consecutive trials. Representative results (means ± SEM) from one experiment are shown. Significance in (c) was determined using 2-tailed Mann-Whitney U Test. Significance in (a) was determined using 2-way ANOVA Test.

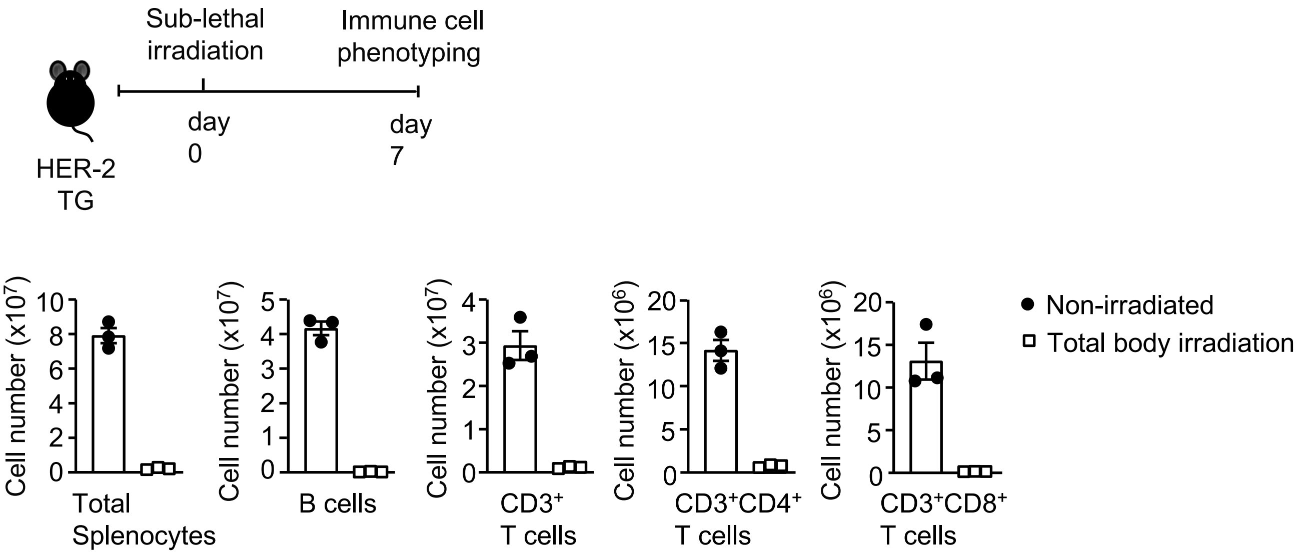

Sub-lethal irradiation
Immune cell phenotyping
HER-2 TG
day 0
day 7
Cell number (x10^7)
Total Splenocytes
B cells
CD3+ T cells
Cell number (x10^6)
CD3+CD4+ T cells
CD3+CD8+ T cells
Non-irradiated
Total body irradiation


**Fig. S32**

***Figure S32. Sublethal irradiation promotes lymphopenia in HER-2 TG mice.*** HER-2 TG mice were sublethally irradiated (4 Gy) and the number of total live splenocytes, $CD19^+$ B cells, $CD3^+$ T cells, $CD3^+CD4^+$ T cells and $CD3^+CD8^+$ T cells were determined after 7 days by flow cytometry. Representative results (means ± SEM) from one experiment are shown.
